# Analysis of multi-way chromatin interactions at sub-nucleosome resolution identifies synergistic nano-scale interactions within super-enhancers

**DOI:** 10.64898/2026.05.21.726779

**Authors:** Shyam Ramasamy, Yi Zhu, Magdalena A. Karpinska, Ilya M. Flyamer, Johannes Söding, A. Marieke Oudelaar

## Abstract

In-depth understanding of the 3D organization of the genome and its role in the regulation of gene expression requires techniques to measure complex chromosome conformations at single alleles at high resolution. Here, we present multi-way Micro-Capture-C (mwMCC), a method to detect concurrent interactions between multiple chromatin regions at single alleles at sub-nucleosome resolution. mwMCC resolves the internal 3D structure of super-enhancers and identifies synergistic multi-way interactions between their constituent elements and gene promoters, providing a structural basis for cooperation within super-enhancers in nano-scale chromatin hubs. The dynamic formation and dissolution of these hubs during cellular differentiation correlates with up- and down-regulation of gene expression. Although large-scale chromatin hubs are dependent on the architectural protein CTCF, nano-scale hubs are robust to CTCF depletion. Instead, integration of footprinting analysis and protein occupancy data indicates a role for transcription factors and the co-activator Mediator in the formation of synergistic enhancer-promoter interactions within nano-scale chromatin hubs.

## INTRODUCTION

Precise spatio-temporal regulation of gene expression is controlled by the *cis*-regulatory elements of the genome, which include enhancers and gene promoters. Active enhancers recruit transcription factors and co-activators, which stimulate assembly and activation of the transcription machinery at gene promoters^1^. In mammals, most genes are regulated by multiple enhancers, which can be separated by large genomic distances from the genes they control. To activate genes, enhancers interact with their target gene promoter(s) in specific 3D chromatin structures^2-6^. Therefore, a molecular understanding of enhancer-mediated gene regulation requires in-depth characterization of 3D genome organization.

Over the last two decades, Chromosome Conformation Capture (3C) methods have been instrumental in uncovering principles of 3D genome organization. 3C methods are based on digestion and subsequent proximity ligation of crosslinked chromatin to detect spatial proximity between DNA sequences using high-throughput sequencing^7,8^. The application of 3C to study 3D gene regulation at the level of individual *cis*-regulatory elements and transcription factor binding sites requires analysis with very high resolution and sensitivity. The resolution and sensitivity of 3C is predominantly determined by the digestion strategy and sequencing depth. Traditional 3C methods use restriction enzymes for chromatin digestion; their resolution is therefore limited by the frequency and distribution of restriction sites in the genome, as these are the only sites at which ligation junctions (i.e., interactions) can be detected^9^. This limitation has been overcome by implementing digestion with micrococcal nuclease (MNase), which digests the genome largely independent of DNA sequence and therefore allows for 3C analysis with improved resolution^10^. Micro-C^10-13^, the first MNase-based 3C method, is a genome-wide approach, and is therefore limited by sequencing depth. The resolution of 3C can be further increased by implementing MNase digestion in targeted 3C methods, in which 3C libraries are enriched for selected genomic regions to allow for more targeted and thus deeper sequencing. Although these methods do not generate genome-wide data, they have the advantage that they can generate very high-resolution data in selected genomic regions of interest^14-18^. Targeted methods include Micro Capture-C (MCC)^15^, which targets capture oligonucleotides to multiplexed individual viewpoints and can generate high-resolution interaction profiles for those viewpoints, as well as Tiled MCC (TMCC)^14^, Region Capture Micro-C (RCMC)^16^, and MCC ultra (MCCu)^17^, which “tile” capture oligonucleotides across larger regions to generate high-resolution contact matrices for those regions.

Although MNase-based 3C methods allow for high-resolution data generation, a remaining limitation is that they cannot resolve how multiple *cis*-regulatory elements interact together in higher-order 3D chromatin structures at individual alleles. Current MNase-based 3C methods predominantly detect pair-wise interactions in populations of cells and therefore cannot distinguish simultaneous, cooperative interactions in individual cells from mutually exclusive interactions that occur independently in different cells. This hampers our ability to dissect how 3D enhancer-promoter interactions influence gene expression, since accurate gene regulation relies on cooperation between multiple *cis*-regulatory elements^19-22^. 3D relationships between multiple interacting *cis-*regulatory elements can be disentangled by 3C-like techniques based on physical separation and labeling of crosslinked chromatin interactions^23-26^, as well as by multi-way 3C methods, which use (relatively) long sequencing reads to capture multiple interactions derived from individual alleles^27-33^. However, current methods do not allow for analysis of multi-way interactions at the level of individual *cis*-regulatory elements, since they are based on restriction enzyme digestion and therefore limited in resolution. Furthermore, due to technical limitations, it is not straightforward to adapt current MNase-based 3C protocols for multi-way interaction analysis.

Here, we present an optimized targeted MNase-based 3C protocol that allows for the generation of single-viewpoint interaction profiles as well as regional contact matrices with improved efficiency and throughput at reduced sequencing costs. This protocol enables further development into a multi-way MNase-based 3C method (multi-way MCC, mwMCC) that allows for robust analysis of multi-way chromatin interactions at sub-nucleosome resolution. In addition, we present an automated computational pipeline for pair-wise and multi-way MNase-based 3C data analysis. Using mwMCC, we resolve the internal 3D structure of super-enhancers, and identify extensive multi-way interactions between constituent super-enhancer elements and gene promoters within nano-scale chromatin hubs. In addition, we characterize nano-scale hubs during cellular differentiation and their relationship with gene expression. Furthermore, by integrating mathematical modelling, we analyze synergy of multi-way chromatin interactions and provide estimates for their absolute probabilities. Finally, by integrating mwMCC data with protein perturbation, footprinting analysis, and protein occupancy data, we provide insight into the mechanisms that underly the formation of synergistic enhancer-promoter interactions within nano-scale chromatin hubs.

## RESULTS

### Optimization of targeted MNase-based 3C

3C methods based on MNase digestion offer superior resolution compared to 3C methods based on digestion with restriction enzymes, but are associated with technical challenges that limit their usability and adaptability. These challenges are twofold: (1) MNase levels need to be carefully titrated in order to achieve sufficient chromatin digestion but avoid over-digestion, after which linkers are too short for efficient ligation; (2) ligation after MNase digestion is inherently inefficient because MNase digestion results in heterogeneous, frayed DNA ends that require extensive end-repair and rely on blunt ligation. Inefficient ligation results in a low proportion of DNA fragments with a “valid” ligation junction that informs on chromatin interactions. It is possible to increase the proportion of DNA fragments with valid junctions in 3C sequencing libraries by incorporating biotinylated nucleotides during end repair and using streptavidin to enrich for DNA fragments with internal biotin that likely contain a valid junction. This has the advantage that the majority of sequencing reads are informative. Biotinylation is incorporated in current Micro-C and RCMC protocols^11-13,16^. A disadvantage of biotinylation is that the additional steps increase loss of material, which reduces library complexity, and interfere with the efficiency of subsequent biotin-mediated enrichment for regions of interest in targeted 3C methods. For this reason, the MCC, TMCC, and MCCu methods do not include biotinylation and benefit from potential lower input requirements and a higher proportion of reads in targeted genomic regions, at the cost of a low proportion of reads with a valid junction^14,15,17^.

Multi-way 3C methods rely on capturing multiple valid ligation junctions that are derived from single alleles in individual sequencing reads. Such methods therefore require efficient ligation and/or the ability to enrich for valid ligation junctions. In addition, they require very high library complexity to allow for analysis of multi-way contacts at sufficient depth in order to robustly identify patterns of enrichment and depletion in targeted regions. To enable the development of a targeted MNase-based multi-way 3C method and improve the conventional pair-wise methods, we heavily optimized the current MCC protocol and incorporated the following main changes: (1) binding of crosslinked cells to magnetic beads coated with Concanavalin A (ConA), which reduces losses during 3C library preparation and therefore increases library complexity and minimizes input requirements, and has the additional benefit that it makes chromatin less prone to over-digestion with MNase; (2) incorporation of biotinylated nucleotides in the digested DNA during end-repair; and (3) optimization of the target enrichment protocol for biotinylated 3C libraries (**Figure 1a**). The optimized procedure results in 3C libraries with: (1) high library complexity relative to input material; (2) a large proportion of valid ligation junctions; and (3) a high proportion of reads on target. A detailed overview of all parameters that we systematically tested and optimized is included in **Supplementary Note 1**. Of note, during the preparation of this manuscript, a low-input Micro-C method (Pico-C) was published, which also reports on benefits of the use of ConA beads in MNase-based 3C protocols^34^.

**Figure 1:**
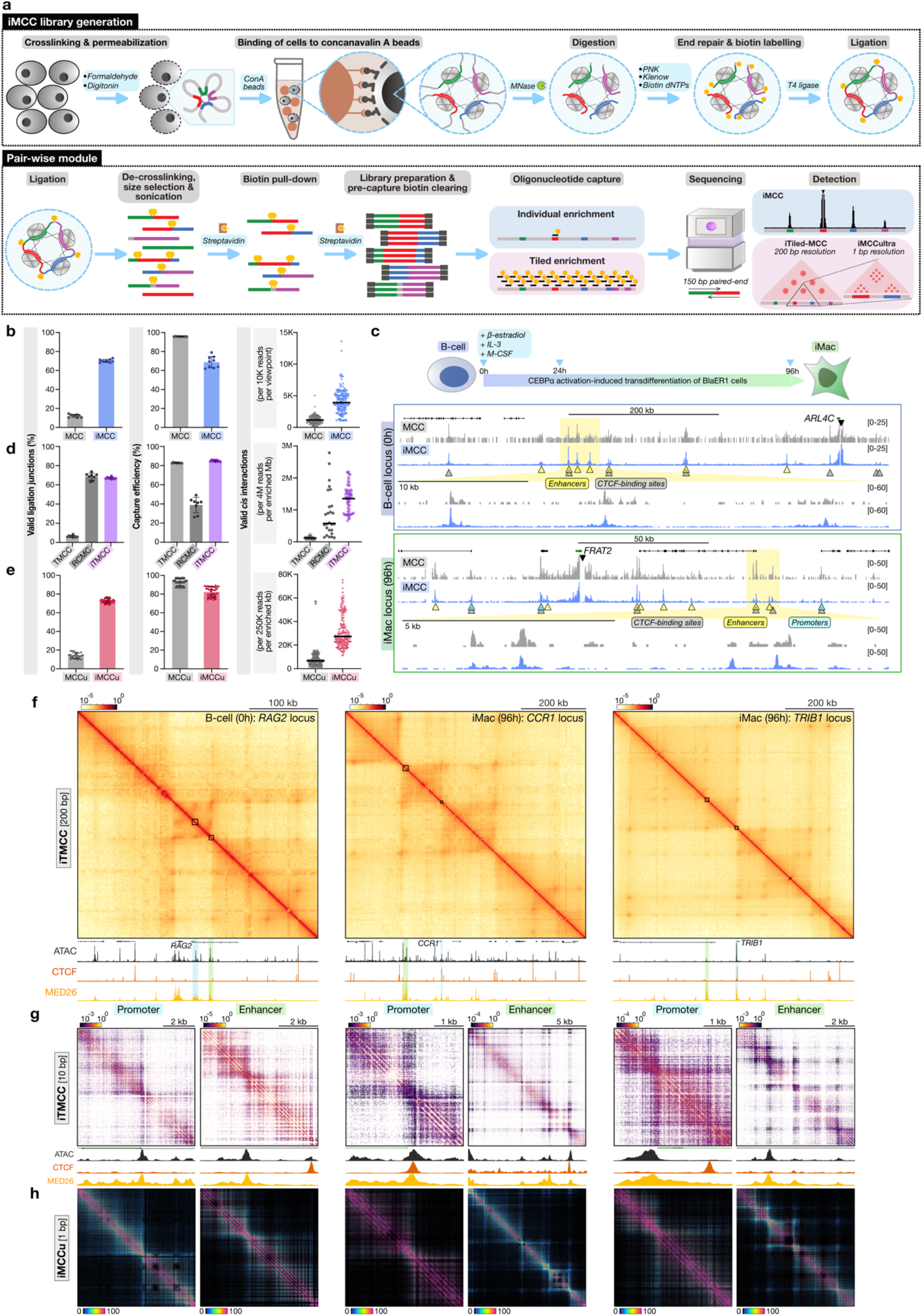
Improved targeted MNase-based 3C. **a**, Outline of the experimental procedure of the improved Micro-Capture-C (iMCC)-based methods. **b**, Percentage of valid ligation junctions in reads (left), percentage of reads on target (middle), and number of valid ligation junctions (interactions) in cis per 10,000 sequenced reads in conventional Micro-Capture-C (MCC) and improved Micro-Capture-C (iMCC) data in BLaER1 cells. **c**, Top: Outline of the BLaER1 lymphoid-to-myeloid transdifferentiation system. Bottom: MCC and iMCC interaction profiles of the ARL4C and FRAT2 promoters, generated from data down-sampled to an average of 10,000 reads per viewpoint, in BLaER1 cells. Black triangles indicate the viewpoint; yellow, blue, and grey triangles indicate enhancers, promoters, and CTCF-binding sites, respectively. Coordinates (hg38): ARL4C locus, chr2:233908830-234563849; FRAT2 locus, chr2:234131765-234169572. **d**, Comparison of conventional Tiled Micro-Capture-C (TMCC; BLaER1 cells), Region Capture Micro-C (RCMC; mES cells), and improved Tiled Micro-Capture-C (iTMCC; BLaER1 cells) data, as in **b**. The right graph shows the number of valid cis interactions per enriched Mb per 4M sequencing reads. **e**, Comparison of conventional Micro-Capture-C ultra (MCCu; mES cells) and improved Micro-Capture-C ultra (iMCCu; BLaER1 cells) data, as in **b**. The right graph shows the number of detected cis interactions per enriched kb per 250k sequencing reads. **f**, iTMCC large-scale contact matrices (200 bp resolution) of the RAG2, CCR1, and TRIB1 loci in BLaER1 cells. Small black squares and shadings in blue and green indicate regions shown at higher resolution in **g** and **h**. Profiles below show ATAC-seq, CTCF ChIP-seq, and MED26 ChIPmentation. Axes scales: RAG2 locus, 0-2130 (ATAC-seq), 0-10994 (CTCF), 0-2527 (MED26); CCR1 locus, 0-873 (ATAC-seq), 0-6000 (CTCF), 0-3766 (MED26); TRIB1 locus, 0-5963 (ATAC-seq), 0-6348 (CTCF), 0-4182 (MED26). Coordinates (hg38): RAG2 locus, chr11: 36426478-36776478; CCR1 locus, chr3:45911998-46651998; TRIB1 locus, chr8:125071000-125771000. **g**, iTMCC nano-scale contact matrices (10 bp resolution) of the RAG2, CCR1, and TRIB1 loci in BLaER1 cells. Profiles below show ATAC-seq, CTCF ChIP-seq, and MED26 ChIPmentation. Axes scales: RAG2 promoter locus, 0-1718 (ATAC-seq), 0-2000 (CTCF), 0-1233 (MED26); RAG2 enhancer locus, 0-1826 (ATAC-seq), 0-2360 (CTCF), 0-2831 (MED26); CCR1 promoter locus, 0-542 (ATAC-seq), 0-6626 (CTCF), 0-1660 (MED26); CCR1 enhancer locus, 0-1209 (ATAC-seq), 0-720 (CTCF), 0-6075 (MED26); TRIB1 promoter locus, 0-8268 (ATAC-seq), 0-9086 (CTCF), 0-3320 (MED26); TRIB1 enhancer locus, 0-1437 (ATAC-seq), 0-1500 (CTCF), 0-5546 (MED26). Coordinates (hg38): RAG2 promoter locus, chr8:125071000-125771000; RAG2 enhancer locus, chr11:36617349-36623306; CCR1 promoter locus, chr3:46206211-46210210; CCR1 enhancer locus, chr3:46089378-46102846; TRIB1 promoter locus, chr8:125428824-125432981; TRIB1 enhancer locus, chr8:125340255-125348155. **h**, iMCCu contact matrices (1 bp resolution) of the RAG2, CCR1, and TRIB1 loci in BLaER1 cells. Coordinates as in **g**.

The optimized procedure forms the basis for a user-friendly, improved MCC (iMCC) protocol (**Figure 1a, Supplementary Note 2**). To further promote the usability of the improved workflow, we built Capture MNase-based 3C Analysis Pipeline (C-MAP), an analysis pipeline for targeted MNase-based 3C data that is fast, modular, flexible, and user-friendly (**Extended Data Figure 1a**). The iMCC procedure allows for relatively low-input material (**Supplementary Note 1**) and enables generation of high-quality 3C libraries with a ∼6-fold higher proportion of valid ligation junctions, as demonstrated by comparison of iMCC and MCC data in BLaER1 cells (**Figure 1b, left panel**). As expected, the integration of biotinylation to enrich for ligation junctions reduces the capture efficiency, but with the optimized target enrichment procedure, the proportion of reads on target remains ∼70% (**Figure 1b, middle panel**). As a result, iMCC identifies substantially more informative interactions (defined as valid ligation junctions *in cis*) with the same number of sequencing reads compared to the conventional MCC method (**Figure 1b, right panel**) and therefore allows for the generation of high-quality interaction profiles with single-base-pair resolution with as little as 10,000 reads per viewpoint (**Figure 1c**).

We further adapted the iMCC workflow to develop an improved TMCC (iTMCC) protocol that allows for efficient data generation with a larger proportion of valid junctions compared to TMCC and a larger proportion of reads on target compared to RCMC (**Figure 1d**). The iTMCC method is therefore more efficient compared to current methods (**Figure 1d**) and can generate high-quality contact matrices (**Figure 1f**) at 500 bp resolution with <10 M reads per targeted Mb (**Methods**). An additional advantage over RCMC, is that iTMCC allows for direct identification of the exact positions of the ligation junctions because the libraries are sonicated to ∼200 bp and sequenced with 150 bp paired-end reads (**Figure 1a**). In contrast, RCMC libraries are comprised of extracted di-nucleosomal DNA fragments (300-400 bp), which are usually sequenced with 50 bp paired-end reads. This means that the exact positions of the ligation junctions cannot be determined and are inferred with an error of 200-300 bp, which therefore limits the resolution. Because (i)TMCC directly identifies ligation junctions, there is no absolute resolution limit, which allows for analysis of nano-scale features of genome folding in regions close to the diagonal (**Figure 1g**). Since (i)TMCC resolution is only limited by sequencing depth, it is possible to generate contact matrices with single-base-pair resolution by reducing the size of the targeted regions, as has been demonstrated by the MCCu method (**Figure 1a**). Notably, the improved MCC workflow targeted at relatively small regions (iMCCu) can also generate single-base-pair contact matrixes but needs substantially fewer sequencing reads compared to MCCu due to the improved efficiency (**Figure 1e,h**). In summary, the optimized iMCC, iTMCC, and iMCCu workflows generate high-quality data with greater efficiency compared to current targeted MNase-based 3C methods, resulting in data with higher resolution and sensitivity relative to input material and sequencing depth. As a result, the improved workflows allow for the identification of both large- and nano-scale cell-type-specific chromatin structures (**Extended Data Figure 1b-e**).

### Development of a targeted MNase-based multi-way 3C method

The increased efficiency of the iMCC workflow allows for the development of a targeted multi-way MCC method, based on the detection of multiple ligation junctions (i.e., multi-way interactions) within single, (relatively) long sequencing reads (**Figure 2a**). We tailored mwMCC to be compatible with 300 bp paired-end sequencing platforms from Illumina and Element Biosciences, since 600 sequencing cycles are sufficient to detect junctions between 3-4 interacting MNase-digested fragments, and Illumina and Element Biosciences offer benefits with respect to throughput, accuracy, and costs over traditional long-read sequencing platforms. To enrich libraries for fragments with a length of 500-600 bp, we optimized the fragmentation and size selection steps in the iMCC workflow. We found that gel extraction allows for the most robust enrichment for fragments in the desired size range and most efficient detection of multi-way interactions, and thereby outperforms sonication and bead-based size selection approaches (**Supplementary Note 1**).

**Figure 2:**
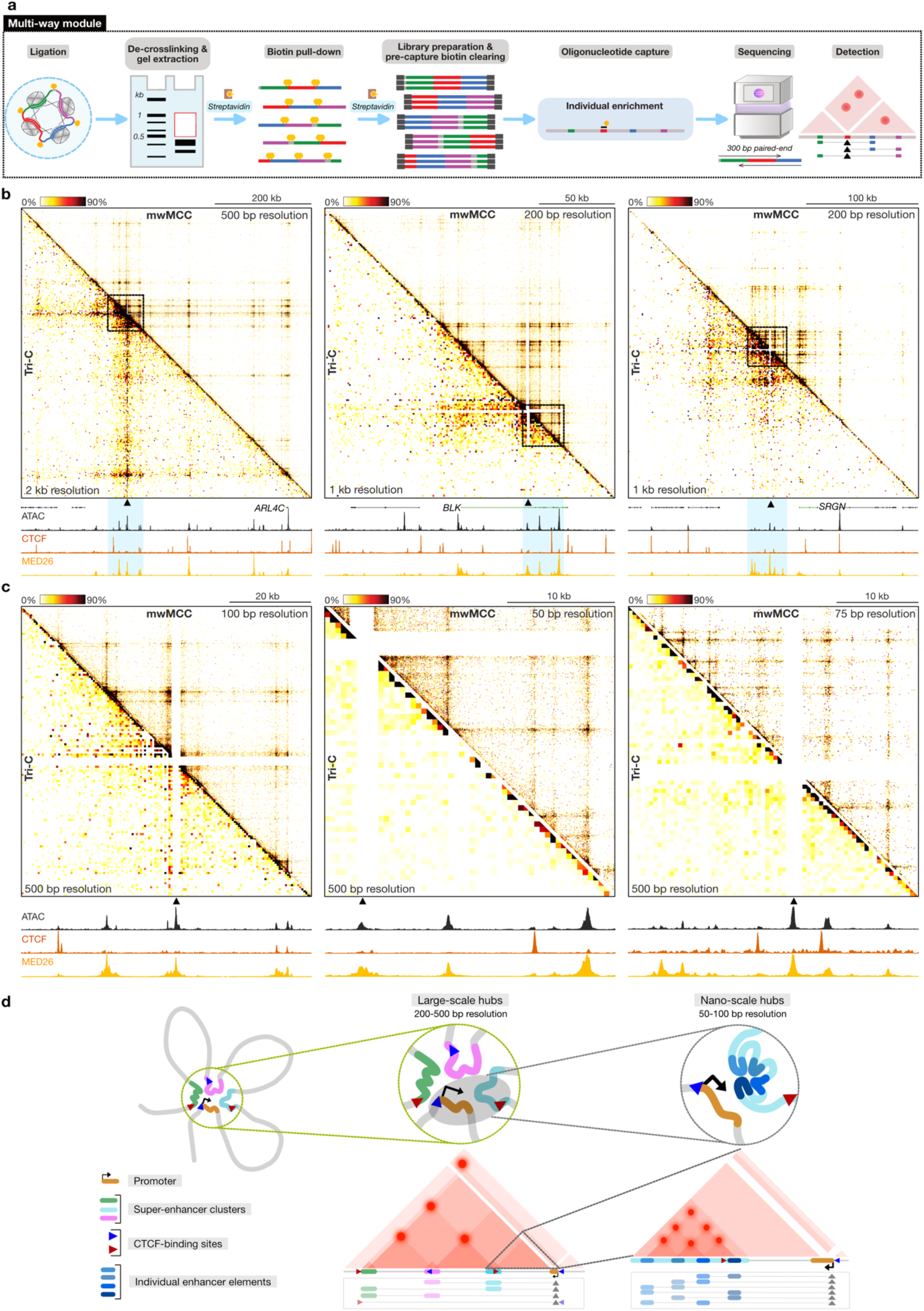
Experimental procedure and benchmarking of multi-way Micro-Capture-C. **a**, Outline of the experimental procedure of the multi-way Micro-Capture-C (mwMCC) method. **b**, Comparison of large-scale mwMCC (top-right; 200-500 bp resolution) and Tri-C (bottom-left; 1-2 kb resolution) data from enhancer viewpoints (VPs) in BlaER1 cells at 0h (ARL4C and BLK) and 96h (SRGN) post-differentiation induction, visualized in VP-specific contact matrices that show the frequencies at which two regions interact simultaneously with the VPs. VPs are indicated with black arrows directly below the matrices and proximity signals around the VPs are excluded. Note that the Tri-C VPs are determined by the location of restriction sites and are not identical to the mwMCC VPs. Black squares and blue shadings indicate regions shown at higher resolution in **c**. Profiles below show ATAC-seq, CTCF ChIP-seq, and MED26 ChIPmentation. Axes scales: ARL4C locus, 0-3204 (ATAC-seq), 0-7030 (CTCF), 0-5636 (MED26); BLK locus, 0-3953 (ATAC-seq), 0-13639 (CTCF), 0-4617 (MED26); SRGN locus, 0-5464 (ATAC-seq), 0-4712 (CTCF), 0-4854 (MED26). Coordinates (hg38): ARL4 locus, chr2:233945774– 234545774; BLK locus, chr5:140539932–140668951; SRGN locus, chr10:68935164–69194029. **c**, Comparison of nano-scale mwMCC (top-right; 50-100 bp resolution) and Tri-C (bottom-left; 500 bp resolution) data, as in **b**. Axes scales: ARL4C locus, 0-2144 (ATAC-seq), 0-5047 (CTCF), 0-4090 (MED26); BLK locus, 0-3953 (ATAC-seq), 0-13634 (CTCF), 0-4617 (MED26); SRGN locus, 0-1960 (ATAC-seq), 0-949 (CTCF), 0-5028 (MED26). Coordinates (hg38): ARL4 locus, chr2:234126834-234197909; BLK locus, chr5:140586172-140614136; SRGN locus, chr10:69041665-69076889.

We benchmarked mwMCC against Tri-C, using previously published data in BLaER1 cells^35^. Tri-C is a multi-way 3C method based on restriction enzyme digestion, which also relies on targeted enrichment of multiplexed viewpoints of interest and therefore allows for analysis at relatively high resolution (500-5000 bp, depending on the size of the viewpoint fragment, library complexity, and sequencing depth)^27^. Both mwMCC and Tri-C data can be visualized in viewpoint-specific 3D contact matrices, which show the frequencies with which two chromatin fragments interact simultaneously with the viewpoint (**Figure 2a**). In these matrices, preferential simultaneous interactions between *cis-*regulatory elements are enriched at the intersections between these elements, whereas mutually exclusive interactions are depleted. Direct comparison of mwMCC and Tri-C data shows that mwMCC detects multi-way chromatin interactions with greater sensitivity (**Figure 2b**) and higher resolution (**Figure 2c**) compared to Tri-C (**Supplementary Note 1**). Whereas Tri-C and other current multi-way 3C methods are mostly limited to analysis of multi-way interactions *between* super-enhancers, the improved resolution of mwMCC allows for analysis of multi-way interactions *within* super*-*enhancers (**Figure 2b**,**c**). With a resolution of up to 50 bp, mwMCC uniquely resolves the internal 3D structure of super-enhancers, and identifies extensive multi-way interactions between the individual constituent elements of super-enhancers within nano-scale hubs (**Figure 2c**,**d**).

An additional advantage of mwMCC is that there are no constraints on the selection of viewpoints. In contrast, Tri-C requires restriction fragments at the targeted viewpoints to be relatively small (<300bp) to facilitate efficient detection of multiple ligation junctions. As a result, Tri-C analysis is usually targeted at enhancers instead of promoters, as this provides more flexibility for viewpoint selection^27^. Since MNase digestion results in uniformly small fragments across the genome, mwMCC does not have this limitation, and allows for analysis of multi-way chromatin interactions for promoter viewpoints and any other viewpoint of choice (**Extended Data Figure 2a,b**).

### Systematic identification of cell-type-specific chromatin hubs

To characterize how multi-way interactions between individual *cis-*regulatory elements form during cellular differentiation, we performed mwMCC experiments over a lymphoid-to-myeloid transdifferentiation course in BLaER1 cells, which are B-cell leukemia cells that can be converted into functional macrophages by exogenous expression of the transcription factor CCAAT enhancer-binding protein alpha (CEBPa; **Figure 1c**)^36^. Based on previous characterizations of this differentiation system^35,37,38^, we focused our analysis on timepoints 0h, 24h, and 96h after differentiation induction, which are representative of a B-cell state, a transition state, and an early macrophage state. We focused our analysis on 51 promoters (**Figure 3**) and enhancers (**Extended Data Figure 3)** of B cell-specific and iMac-specific genes. Consistent with previous analyses in this system^35^, we observe that the formation and dissolution of large-scale chromatin hubs, comprised of multi-way interactions between clusters of enhancers, promoters, and CTCF-binding sites, correlate with upregulation of macrophage-specific genes and downregulation of B-cell specific genes, respectively (**Figure 3a,b, Extended Data Figure 3a,b**). In addition, mwMCC uncovers tissue-specific multi-way interactions between individual *cis-*regulatory elements within nano-scale hubs, which also form and dissolve dynamically over the differentiation course in gene loci that are up- and downregulated, respectively (**Figure 3a,b, Extended Data Figure 3a,b**).

**Figure 3:**
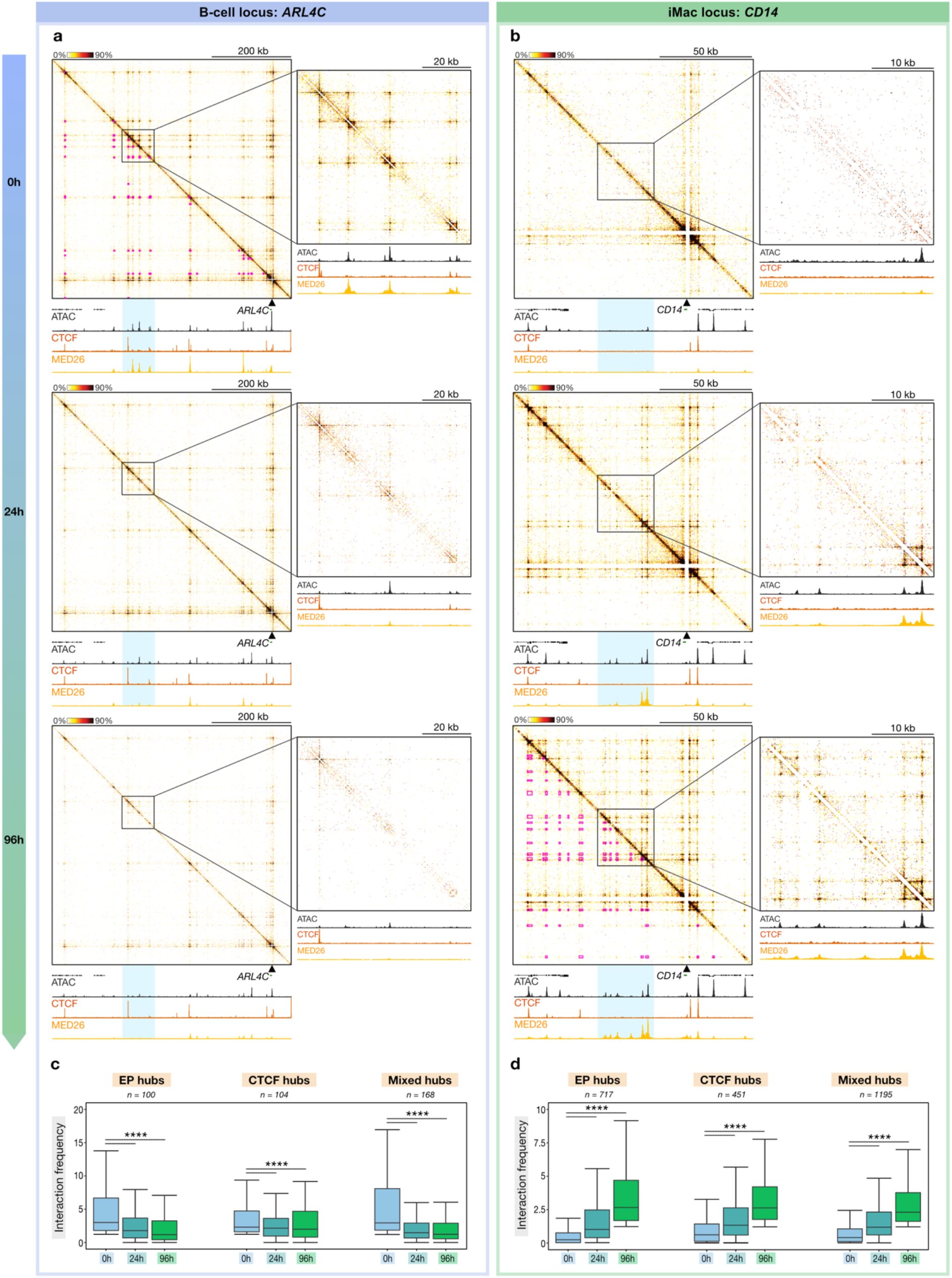
Dynamic formation and dissolution of large-scale and nano-scale chromatin hubs during cellular differentiation. **a**, mwMCC contact matrices of the ARL4C locus (promoter viewpoint; 500/100 bp resolution) through lymphoid-to-myeloid transdifferentiation, as in **2b**. Pink squares indicate loop calls. Axes scales: large-scale, 0-2531 (ATAC-seq), 0-4565 (CTCF), 0-4766 (MED26); nano-scale, 0-1950 (ATAC-seq), 0-4930 (CTCF), 0-4048 (MED26). Coordinates (hg38): large-scale, chr2:233945774-234545774; nano-scale, chr2: 234126834-234197909. **b**, mwMCC contact matrices of the CD14 locus (promoter viewpoint; 200/75 bp resolution) through lymphoid-to-myeloid transdifferentiation, as in **a**. Axes scales: large-scale: 0-3110/(ATAC-seq), 0-6018 (CTCF), 0-7377 (MED26); nano-scale: 0-2182 (ATAC-seq), 0-2000 (CTCF), 0-7557 (MED26). Coordinates (hg38): large-scale, chr5:140539932-140668951; nano-scale, chr5:140586172-140614136. **c**, Multi-way interaction frequencies of promoters of B-cell-specific genes with enhancers and promoters (EP hubs), CTCF-binding sites (CTCF hubs), and a mixture of enhancers, promoters, and CTCF-binding sites (Mixed hubs). Box plots show the interquartile range and median of the data, with whiskers indicating the minima and maxima within 1.5×interquartile ranges. Asterisks indicate statistical significance (two-sided paired Wilcoxon signed-rank test, 96h versus 0h). P-values: EP hubs, P = 2.8 × 10^-8^; CTCF hubs, P = 6.8 × 10^-5^; Mixed hubs, P < 2.2 × 10^-16^. The number of data points (n) in each category is shown above the graph. Data are derived from three biological replicates. **d**, Multi-way interaction frequencies of promoters of iMac-specific genes, as in **c**. P-values: EP hubs, P < 2.2 × 10^-16^; CTCF hubs, P < 2.2 × 10^-16^; Mixed hubs, P < 2.2 × 10^-16^.

The high resolution and sensitivity of mwMCC allow for identification of multi-way interactions between individual *cis-*regulatory elements that are closely spaced together. In addition, the precise nature of the mwMCC data facilitates systematic calling of multi-way interactions (**Methods, Figure 3a,b, Extended Data Figure 3a,b**). Quantification of called multi-way interactions across the 51 loci confirms that the formation and dissolution of chromatin hubs correlate with up- and downregulation of gene expression during lymphoid-to-myeloid transdifferentiation (**Figure 3c,d, Extended Data Figure 3c,d**).

### Synergy and absolute quantification of multi-way interactions

The high resolution and sensitivity of mwMCC offer a unique opportunity to precisely assess to what extent interactions between *cis*-regulatory elements occur independently or synergistically. The detection of multi-way interactions between gene promoters and multiple enhancer elements indicates that these interactions are not mutually exclusive and that a promoter can engage multiple enhancers at the same time. However, it is not immediately apparent whether multi-way interactions between *cis*-regulatory elements occur independently or synergistically. In the example of a gene promoter and two enhancers, the former would indicate that the interactions between the promoter and the two enhancers do not depend on each other but occasionally co-occur by chance. In contrast, the latter would indicate that the interactions of the promoter with the two enhancers are cooperative, for example because the presence of an enhancer (and the proteins it recruits) in the proximity of a promoter stabilizes interactions with an additional enhancer. Such cooperative interactions may be expected in the context of transcriptional condensates, in which the clustering of multiple *cis*-regulatory elements could facilitate the recruitment and concentration of transcription factors, cofactors, and RNAPII, which may in turn stabilize interactions between these elements^39-42^.

If a viewpoint (*V*) interacts independently with two loci, *X* and *Y*, it is expected that the frequency of the multi-way interaction (*V,X,Y*) can be predicted from the frequencies of the pair-wise interactions (*V,X*), (*V,Y*) and (*X,Y*). In contrast, if *V* synergistically interacts with *X* and *Y*, the frequency of the measured multi-way interaction (*V,X,Y*) should be enriched over the multi-way interaction frequency that is predicted from the composite pair-wise interactions^32^. We used high-resolution pair-wise iTMCC data in seven loci to predict the frequency of multi-way interactions with the promoter viewpoint according to an independent interaction model (**Methods, Extended Data Figure 4a,b, Figure 4a**). Next, we calculated the enrichment of the observed multi-way interaction frequencies as measured by mwMCC over the simulated frequencies (**Figure 4b, Extended Data Figure 4c**). Notably, we observe that measured multi-way interactions between *cis-*regulatory elements are strongly enriched, indicating synergistic multi-way interactions between *cis*-regulatory elements, as exemplified by the *SRGN* (**Figure 4b**) and *CCR1* (**Extended Data Figure 4c**) loci. These observations are further confirmed by systematic quantification across the analyzed loci, which shows significant interaction synergy among *cis*-regulatory elements but not among randomly sampled loci in the mwMCC matrix (**Figure 4c**). Categorizing the multi-way interactions between the promoter with (1) enhancers and promoters only; (2) CTCF-binding sites only; (3) an enhancer/promoter and a CTCF-binding site does not reveal clear differences in the extent to which promoter interactions are synergistic across the different classes of *cis-*regulatory elements (**Figure 4c**). Correlating synergy scores with the distance between interacting elements indicates that longer-range multi-way interactions are more synergistic compared to short-range interactions, indicating a bigger dependency on cooperative mechanisms for long-range interactions compared to shorter-range interactions (**Figure 4d**).

**Figure 4:**
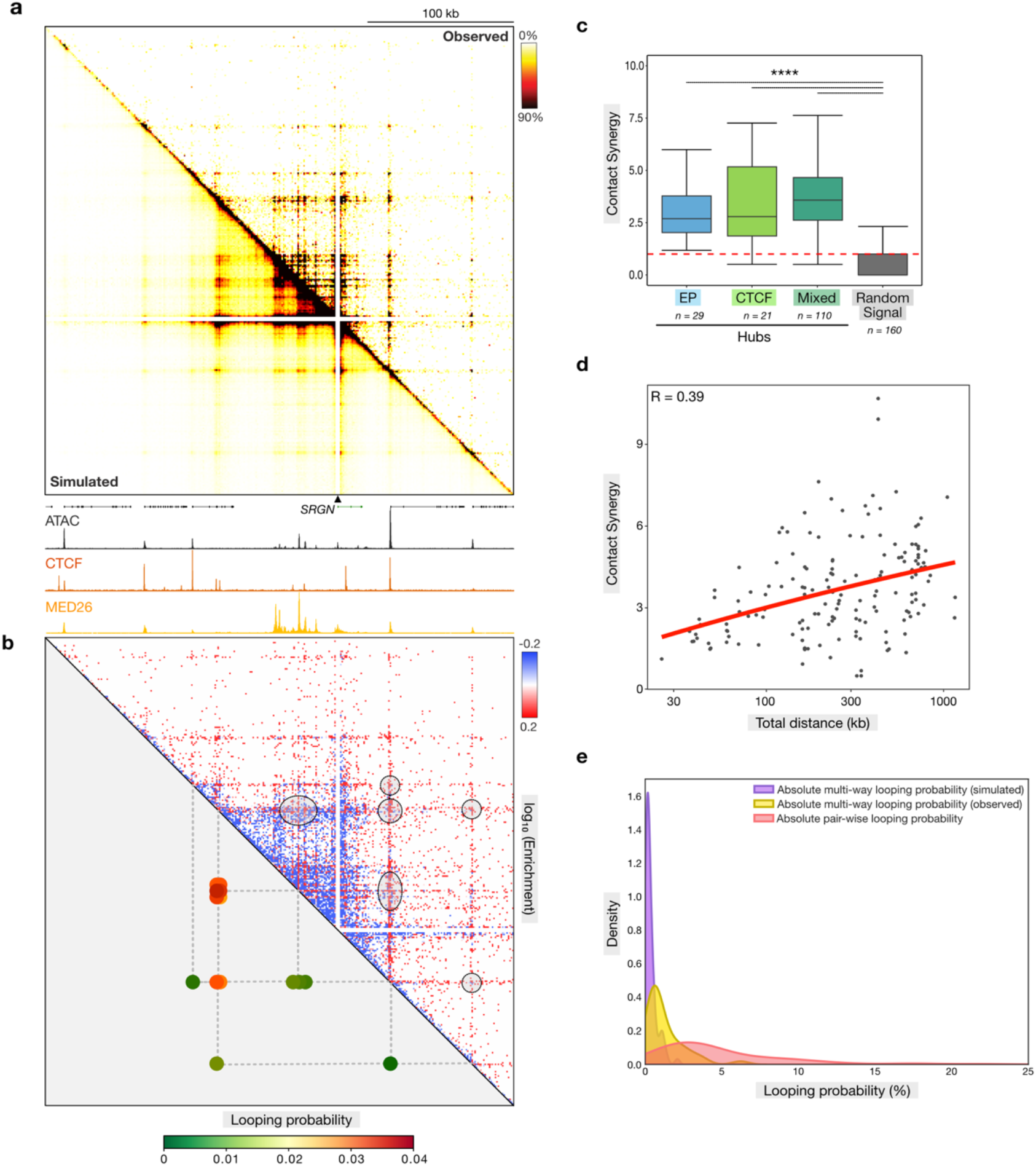
Synergy and absolute quantification of multi-way interactions. **a**, 3D contact matrices of the SRGN locus (promoter viewpoint; 1000 bp resolution) in BLaER1 cells at 96h post-differentiation induction, derived from mwMCC data (top-right) or simulated based on an independent interaction model (bottom-left), as in **2b**. Axes scales: 0-5121 (ATAC-seq), 0-4887 (CTCF), 0-4837 (MED26). Coordinates (hg38): chr10:68888336-69208336. **b**, 3D contact matrices showing enrichment of observed over simulated multi-way contact frequencies (top-right; 1000 bp resolution) and estimated absolute looping probabilities (bottom-left), as in **a. c**, Enrichment of observed over simulated multi-way contact frequencies (contact synergy) of promoters of B-cell- and iMac-specific genes with enhancers and promoters (EP hubs), CTCF-binding sites (CTCF hubs), a mixture of enhancers, promoters, and CTCF-binding sites (Mixed hubs), and randomly sampled loci (Random signal), as in **3c**. Asterisks indicate statistical significance (two-sided Wilcoxon signed-rank test, hubs versus random signal). P-values: EP hubs, P < 2.2 × 10^-16^; CTCF hubs, P = 1.4 × 10^-11^; Mixed hubs, P < 2.2 × 10^-16^. **d**, Correlation between contact synergy and distance between interacting loci, based on Spearman’s correlation. **e**, Distributions of estimated absolute looping probabilities of pair-wise interactions (red), expected multi-way interactions (based on an independent interaction model; purple), and observed multi-way interactions (in mwMCC data; yellow).

Live imaging experiments have shown that interactions between *cis*-regulatory elements in chromatin loops are very dynamic and occur sporadically over time in individual cells^43-49^. A recent study used live imaging data to calibrate relative interactions frequencies as measured by 3C in order to provide estimates of absolute looping probabilities^50^. We used this framework to estimate absolute probabilities of pair-wise chromatin interactions as measured by iTMCC, and obtained mean and maximum probabilities of 4.9% and 24%, respectively, which corresponds well with the reported genome-wide mean and maximum probabilities of 2.3% and 26%, respectively^50^. To get more insight into how absolute probabilities of multi-way interactions compare to pair-wise interactions, we adapted the framework for estimation of absolute interaction probabilities to multi-way interactions, using assumptions that allow us to estimate lower bound probabilities (**Methods, Extended Data Figure 4a**). This results in mean and maximum probabilities of 1.8% and 12%, respectively (**Figure 4b,e, Extended Data Figure 4c**). Notably, these probabilities are significantly higher than would be expected based on an independent interaction model (**Figure 4e**), which is consistent with the strong synergistic effects we observe for multi-way interactions between *cis*-regulatory elements.

### The role of CTCF in large-scale and nano-scale hub formation

We next sought to provide more insight into the molecular mechanisms that underly the formation of chromatin hubs and their potential role in gene regulation. Cohesin and CTCF are known to have an important role in 3D genome organization as they mediate the process of loop extrusion, in which cohesin dynamically extrudes loops that are demarcated by CTCF-bound sites^51,52^. It has been suggested that loop extrusion contributes to (long-range) enhancer-promoter interactions, although its precise role in enhancer-mediated gene activation remains debated^53-65^.

We previously showed that CTCF depletion over the BLaER1 transdifferentiation course does not only reduce multi-way interactions between CTCF-binding sites, but also between enhancers and promoters, leading to a strong impairment of chromatin hub formation, as measured by Tri-C. However, pair-wise interactions between enhancers and promoters, as measured by MCC, were not strongly affected by CTCF depletion, and gene expression changes were minimal^35^. These results indicate that CTCF drives the formation of chromatin hubs and that multi-way interactions in the context of chromatin hubs do not play a major role in the regulation of gene expression. However, a limitation of this study is that it is based on analysis with Tri-C with a resolution of 2-4 kb. Since promoters and enhancers often have CTCF-binding sites in close proximity, it is difficult to separate interactions between enhancers and promoters from interactions between CTCF-binding sites based on Tri-C analysis. Furthermore, based on functional dissections of super-enhancer activity^66-69^, it could be expected that cooperative interactions between enhancer elements *within* a super-enhancer have a more pronounced function in the regulation of gene expression compared to cooperative interactions *between* separate super-enhancers. However, due to the limited resolution of the Tri-C data, the role of CTCF in the formation of cooperative interactions within super-enhancers at nano-scale and their potential function in gene regulation remain unclear.

To revisit the role of CTCF in the formation of chromatin hubs, we performed high-resolution mwMCC experiments in BLaER1 cells after auxin-mediated CTCF depletion throughout the differentiation course (**Figure 5a**)^35,38^. As expected, we observe that CTCF depletion leads to a strong reduction in multi-way interactions that directly involve CTCF-binding sites (**Figure 5b,c**). Notably, we also observe a reduction in long-range multi-way interactions that only involve enhancers and promoters upon CTCF depletion (**Figure 5b,c**). These observations are in line with previous Tri-C data^35^. However, short-range multi-way interactions between enhancers and promoters that form in the context of nano-scale chromatin hubs are not significantly affected by CTCF depletion (**Figure 5b,d**). Together, this shows that large- and nano-scale hubs are mechanistically distinct, as large-scale hubs are dependent on CTCF for their formation, whereas nano-scale hubs form independently of CTCF.

**Figure 5:**
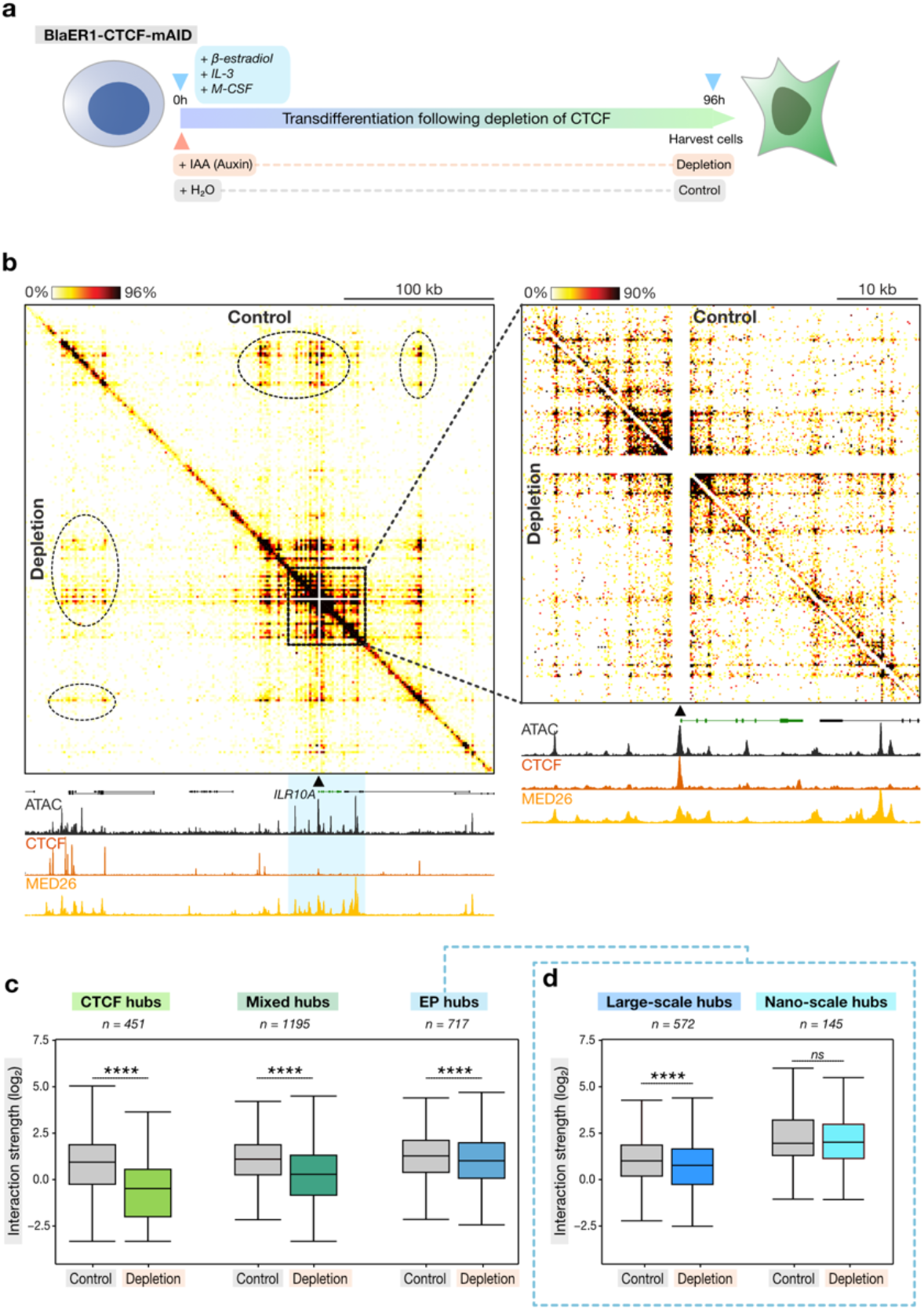
CTCF depletion weakens large-scale chromatin hubs but does not affect nano-scale enhancer-promoter hubs. **a**, Outline of CTCF depletion experiments during lymphoid-to-myeloid transdifferentiation. **b**, mwMCC contact matrices of the IL10RA locus (promoter viewpoint; 1500/200 bp resolution) in control (top-right) and CTCF-depleted (bottom-left) BLaER1 cells at 96h post-differentiation induction, as in **2b**. Axes scales: large-scale: 0-1414/(ATAC-seq), 0-12138/(CTCF), 0-1299/(MED26); nano-scale: 0-1414 (ATAC-seq), 0-2019 (CTCF), 0-1299 (MED26). Coordinates (hg38): large-scale, chr11:117790905-118103188; nano-scale, chr5:117966768-118015968. **c**, Multi-way interaction frequencies of promoters of iMac-specific genes with CTCF-binding sites (CTCF hubs), a mixture of enhancers, promoters, and CTCF-binding sites (Mixed hubs), and enhancers and promoters (EP hubs) in control and CTCF-depleted BLaER1 cells at 96h post-differentiation induction, as in **3c**. Asterisks indicate statistical significance (two-sided paired Wilcoxon signed-rank test, control versus depletion). P-values: CTCF hubs, P < 2.2 × 10^-16^; Mixed hubs, P < 2.2 × 10^-16^; EP hubs, P = 2.7 × 10^-10^. **d**, Multi-way interaction frequencies of promoters of iMac-specific genes with enhancers and promoters separated by more than 10 kb (large-scale hubs) or less than 10 kb (nano-scale hubs) in control and CTCF-depleted BLaER1 cells at 96h post-differentiation induction, as in **3c**. P-values: large-scale hubs, P = 5.0 × 10^-10^; nano-scale hubs, P = 0.058.

### Determinants of nano-scale chromatin hubs

It has been suggested that molecular affinity between the transcription factors and co-activators that are recruited to enhancers and promoters contribute to their interactions^2^. To better understand the role of transcription factors in the formation of nano-scale chromatin hubs, we integrated mwMCC data with the high-resolution iMCCu data of enhancer and promoter regions (**Figure 1h**) to perform “footprinting” analysis of the anchors of multi-way interactions in nano-scale hubs. This analysis leverages the fact that transcription-factor-bound sites are protected from MNase digestion, allowing for bioinformatic contact sequence reconstruction^17^ to identify the DNA sequences involved in observed interactions. In the *IGLL5* locus, which is expressed in B-cells but not in macrophages, we observe nano-scale chromatin hubs involving the gene promoter and several enhancers, that specifically form at 0h, but not at 96h post-differentiation induction (**Figure 6a,b**). One of these enhancers (Enhancer 1) is accessible at both 0h and 96h, whereas the other enhancer (Enhancer 2) and the promoter are inactive at 96h, as is apparent from the regular pattern across the diagonal that indicates the presence of regularly spaced nucleosomes and is typical for inactive regions of chromatin^13,14^ (**Figure 6a-d**). Contact sequence reconstruction identifies specific interactions between binding sites of transcription factors that are active in both B-cells and macrophages in Enhancer 1, such as ETS, FOS, and JUN transcription factors (**Figure 6e**). In contrast, interactions in the promoter (**Figure 6f**) and Enhancer 2 (**Figure 6g**) predominantly involve B-cell-specific transcription factors, such as BCL11A, EBF1, FOXO1, and PAX5. We observe similar patterns in the *MPEG* locus, which is specifically expressed in macrophages and where we observe interactions between macrophage-specific transcription factors in the active state at 96h, which are not detectable at 0h (**Extended Data Figure 6**). Together, this suggests a role for tissue-specific transcription factors in the formation of nano-scale chromatin hubs.

**Figure 6:**
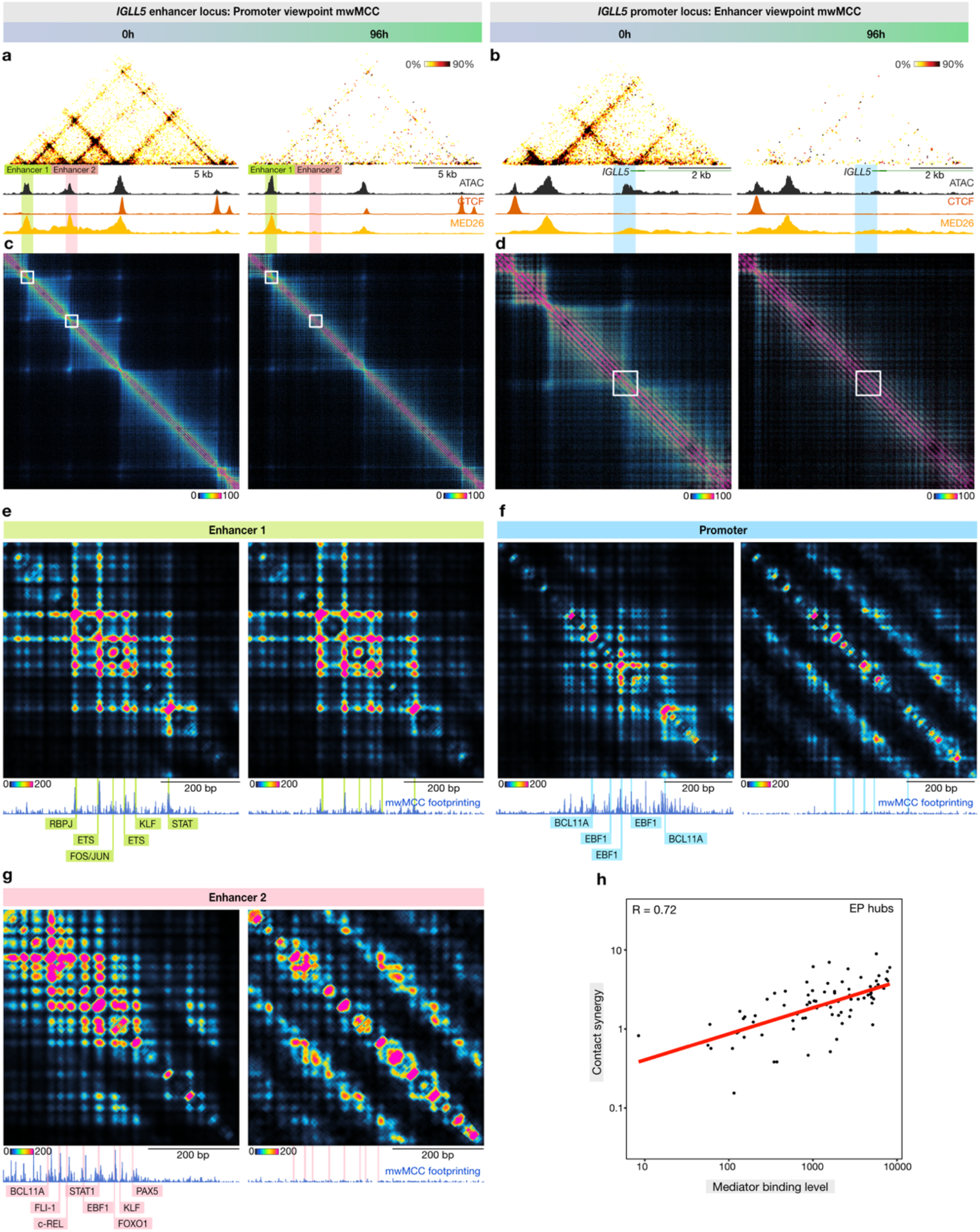
Footprinting analysis of nano-scale chromatin hubs. **a**, mwMCC contact matrices of the IGLL5 enhancers (promoter viewpoint outside matrix, not shown; 100 bp resolution) in BLaER1 cells at 0h (left) and 96h (right) post-differentiation induction, as in **2b**. Axes scales: 0h, 0-1441 (ATAC-seq), 0-7203 (CTCF), 0-7389 (MED26); 960h, 0-1441 (ATAC-seq), 0-7203 (CTCF), 0-7389 (MED26). Coordinates (hg38): chr22:22927192-22944551. **b**, mwMCC contact matrices of the IGLL5 promoter-proximal region (enhancer viewpoint outside matrix, not shown; 100 bp resolution), as in **a**. Axes scales: 0h, 0-982(ATAC-seq), 0-6691 (CTCF), 0-8776 (MED26); 96h, 0-982 (ATAC-seq), 0-6691 (CTCF), 0-8776 (MED26). Coordinates (hg38): chr22:22883520-22891019. **c**, iMCCu contact matrices of the IGLL5 enhancers (1 bp resolution), as in **a. d**, iMCCu contact matrices of the IGLL5 promoter-proximal region (1 bp resolution), as in **b. e-g**, Contact sequence reconstruction matrix, mwMCC footprinting, and transcription factor motif annotation of Enhancer 1 of the IGLL5 locus (**e**), the promoter of the IGLL5 locus (**f**), and Enhancer 2 of the IGLL5 locus (**g**), in BLaER1 cells at 0h (left) and 96h (right) post-differentiation induction. **h**, Correlation between contact synergy and Mediator binding levels, based on Spearman’s correlation.

Transcription factors bound at enhancers recruit the co-activator Mediator, which interacts with the pre-initiation complex at gene promoters and stimulates gene transcription^70^. The Mediator complex has therefore been suggested to function as a bridge between enhancers and promoters^71^. In addition, Mediator has been proposed to contribute to the formation of transcriptional condensates due to its ability to interact with a range of transcription factors and its propensity to form clusters (with RNAPII) that exhibit properties associated with biomolecular condensates^41,72^. Consistent with these roles for Mediator, we previously showed that rapid depletion of the Mediator complex leads to a reduction in enhancer-promoter interactions^73^. To explore the potential role of Mediator in cooperative multi-way interactions, we calculated a synergy score for cooperative interactions involving enhancers and promoters. Notably, this synergy score correlates very strongly with enrichment of Mediator at the corresponding loop anchors, as signified by a correlation coefficient of 0.72 (**Figure 6h**). This is consistent with a potential role for Mediator in the formation of nano-scale regulatory chromatin hubs.

## DISCUSSION

We present an optimized procedure for targeted MNase-based 3C that is more efficient and cost-effective compared to current methods and uniquely allows for targeted analysis of multi-way interactions at sub-nucleosome resolution. In addition, we present a fast and user-friendly computational pipeline for pair-wise and multi-way MNase-based 3C data analysis. Leveraging these developments, we analyze the internal 3D structure of super-enhancers and identify strong and extensive multi-way interactions between the constituent elements of super-enhancers and gene promoters within nano-scale hubs, which are associated with gene activation during cellular differentiation. Integration of mathematical modelling demonstrates that these multi-way interactions are synergistic, providing a structural basis for cooperative behavior of super-enhancer elements in the regulation of gene expression.

Of note, previous analyses of 3way-4C data, generated with restriction enzyme digestion, did not detect strong synergy in regulatory multi-way interactions^32^. It is possible that this discrepancy is explained by differences in resolution, since synergistic effects of focal regulatory interactions may be averaged out by non-synergistic interactions in the immediate surrounding regions, which cannot be distinguished in lower resolution analysis. By adapting a recently published framework for calculation of absolute interaction probabilities^50^, we estimate the mean and maximum of multi-way interaction probabilities to be 1.8% and 12%, respectively. These lower-bound estimates are consistent with recent imaging studies, which show that three-way interactions between enhancers and promoters in the *Nanog* locus in mouse embryonic stem cells (mESCs) occur with a frequency of ∼25%^74^, and that interactions between three super-enhancers in mESCs occur in 5-10% of cells^75^.

By performing mwMCC experiments after auxin-mediated depletion of CTCF, we show that large-scale multi-way interactions between *cis*-regulatory elements are dependent on CTCF. Consistent with previous lower-resolution analysis with Tri-C^35^, we find that the effect of CTCF depletion is most pronounced on multi-way interactions involving CTCF-binding sites, but also significant for multi-way enhancer-promoter interactions, indicating that large-scale chromatin hubs are dependent on cohesin/CTCF-mediated loop extrusion. Interestingly, nano-scale hubs are not weakened upon CTCF depletion. This indicates that cooperative interactions between *cis-*regulatory elements at nano-scale are formed by a different mechanism. Footprinting and correlation analysis suggest a role for tissue-specific transcription factors and the co-activator Mediator in the formation of nano-scale multi-way enhancer-promoter interactions.

Since depletion of CTCF and the associated weakening of large-scale chromatin hubs does not strongly affect gene expression over lymphoid-to-myeloid transdifferentiation^35,38^, we previously concluded that large-scale chromatin hubs do not play a critical role in gene regulation^35^. Importantly, the observation that nano-scale hubs are robust to CTCF depletion indicates that they are not necessarily dispensable for gene regulation. In contrast, it is plausible that nano-scale multi-way interactions between individual elements of super-enhancers have an important regulatory role and form the structural basis for cooperativity between *cis*-regulatory elements that has been reported at this scale^66-69^. We anticipate that the efficiency and usability of mwMCC will facilitate important future investigations to provide more insight into the molecular mechanisms and functions of multi-way interactions in the context of nano-scale chromatin hubs at very high resolution.

## METHODS

### Cell culture

Wild-type^36^ and CTCF-mAID^38^ BlaER1 cells were cultured in RPMI 1640 medium containing 10% FBS (Gibco, 10270106), 25 mM HEPES (Fisher Scientific, 15-630-080) and 2% GlutaMAX Supplement (Thermo Scientific, 35050038). The cells were grown at 37°C and 5% CO_2_, and passaged every 2–3 days, keeping the cell count in the range of 0.2-1×10^6^ cells/mL.

#### BlaER1 differentiation

Wild-type BlaER1 cells were cultured and grown to 70-80% confluency. On the day of transdifferentiation, 16.8×10^6^ cells were seeded in a 150 mm culture dish and treated with transdifferentiation induction medium consisting of fresh culture medium containing 100 nM 17β-estradiol (Calbiochem, 3301), 10 ng/mL human IL-3 (PeproTech, 200-03) and 10 ng/mL human CSF-1 (PeproTech, 300-25). The cells were differentiated for up to 96 h and harvested at the 24 h and 96 h timepoints. Cells cultured in fresh culture medium without transdifferentiation factors were used as the non-differentiated control (0 h timepoint).

#### CTCF depletion

To deplete CTCF, CTCF-mAID BLaER1 cells were subjected to transdifferentiation as described above, using transdifferentiation induction medium supplemented with freshly prepared 500 µM IAA (Sigma-Aldrich, I5148) or H_2_O (control). After 48 h, the cells were replenished with fresh induction medium containing IAA or H_2_O. The cells were harvested at the 96 h timepoint.

### Multi-way Micro-Capture-C

The multi-way Micro-Capture-C (mwMCC) procedure builds on previous protocols^16,76^, which have been extensively optimized and adapted to allow for efficient multi-way analysis. The detailed protocol is included as Supplementary Note 2. mwMCC was performed for 3 biological replicates (with 3-6 technical replicates per biological replicate) per timepoint or condition in BlaER1 WT and CTCF-mAID cells.

#### Crosslinking and permeabilization

Briefly, for each timepoint or condition, 100×10^6^ cells (per biological replicate) were crosslinked with 2% formaldehyde (Thermo Scientific, 28908) for 10 min at room temperature. The reaction was quenched with 0.15 M glycine (Sigma-Aldrich, G7126). The fixed cells were washed once with PBS, split into aliquots of 5×10^6^ cells and permeabilized with 0.005% digitonin (Sigma-Aldrich, D141). After incubation at room temperature for 10 min, the samples were snap-frozen and stored at -80°C.

#### iMCC library generation

Aliquots of fixed and permeabilized cells were thawed on ice. Meanwhile, 100 μL Concanavalin A (ConA) beads (EpiCypher, 21-1411) per sample was activated with cold bead activation buffer (20 mM HEPES pH 7.5 (Jena Bioscience, BU-106-75), 10 mM KCl (Sigma-Aldrich, 1049361000), 1 mM CaCl_2_ (Sigma-Aldrich, C5080), 1 mM MnCl_2_ (Fisher Scientific, 15405419)). The activated beads were added to the samples and the sample-bead mixtures were incubated for 15-30 mins on a rotator at gentle speed. The bead-bound samples were magnetized, cleared and then digested with MNase (NEB, M0247) for 1 h at 37°C and 550 rpm in low Ca^2+^ MNase reaction buffer (10mM Tris-HCl pH 7.5, 10mM CaCl_2_). MNase concentrations (420-560 gel units) were chosen based on batch titration to avoid over- or under-digestion. After incubation, 5mM EGTA (Fisher Scientific, 15415795) was added to quench MNase activity. Following a PBS wash, sample fractions were collected as controls to test digestion efficiency. End preparation and labeling were performed through multiple enzymatic steps to generate ligation-compatible ends in MNase-digested chromatin and incorporate biotinylated dNTPs. The bead-bound digested samples were resuspended in end-repair buffer mix (1X NEBuffer r2.1 (NEB, B6002), 2 mM ATP (Thermo Scientific, R1441), 5 mM DTT (Sigma-Aldrich, 10197777001)) containing 50 U T4 Polynucleotide Kinase (NEB, M0201) and incubated for 30 min at 37°C and 550 rpm. Subsequently, 50 U DNA Polymerase I Large (Klenow) Fragment (NEB, M0210) was added and the samples were incubated for another 30 min at 37°C and 550 rpm. Next, end-label mix (1X T4 DNA Ligase Buffer (NEB, B0202), 1X BSA (Sigma-Aldrich, B8667), 300 μM each of biotin-dATP (Jena Bioscience, BU-835-BIO14), biotin-dCTP (Jena Bioscience, BU-809-BIOX), dGTP (Jena Bioscience, BU-1003), and dTTP (Jena Bioscience, BU-1004)) was added and the samples were incubated for 1-1.5 h at room temperature (20-25°C) and 550 rpm followed by quenching with 30 mM EDTA. Following a PBS wash, the samples were resuspended in ligation mix (1X T4 DNA Ligase Buffer, 1X BSA) containing 10,000 U T4 DNA ligase (NEB, M0202). Ligation reactions were performed overnight at 20°C and 550 rpm. To remove biotinylated dNTPs from unligated ends, the bead-bound ligated samples were cleared, resuspened in 1X NEBuffer 1 (NEB, B7001) and incubated with 1,000 U Exonuclease III (NEB, M0206) for 20 min at 37°C and 550 rpm. Next, samples were reverse crosslinked for at least 6-8 h at 65°C and 1000 rpm by adding 2 mg/mL Proteinase K (Thermo Scientific, EO0491), 0.1 mg/mL RNase A (Thermo Scientific, EN0531) and Buffer AL (Qiagen, 69504). DNA was extracted using DNeasy Blood and Tissue Kit (Qiagen, 69504) and eluted in pre-warmed 150 μL AE Buffer. The resulting iMCC libraries were quantified using Qubit dsDNA BR Kit and ligation efficiencies were assessed on TapeStation and Fragment Analyzer.

#### Gel extraction

DNA fragments of 500-1000 bp were resolved on a 1% agarose gel and were isolated by gel extraction. Briefly, 15-20 μL 6X Gel Loading Dye (NEB, B7024) was added to iMCC library samples. The samples were loaded into combined multi-wells to accommodate the full sample volume, and the gels were run at 120 V for 45-50 min. Gel extraction was performed using GeneJET Gel Extraction Kit (Thermo Scientific, K0692) following manufacturer’s instructions with slight modifications. Gel slices containing the desired DNA fragments were carefully excised, transferred to multiple pre-weighed 2 mL microcentrifuge tubes per sample and weighed (< 1g in each tube). An equal volume of Binding Buffer to gel slice (volume:weight) was added and the mixtures were incubated for 20-30 min until gel slices were completely dissolved. The solubilized gel solutions were transferred to columns and centrifuged at 14,000 x g for 1 min, and this was repeated until the total volume from multiple tubes per sample had been transferred. This was followed by a wash with Wash Buffer and a subsequent dry spin to remove residual ethanol. At the final step, DNA fragments were eluted in pre-warmed 150 μL Elution Buffer. The gel-extracted DNA was quantified using Qubit dsDNA BR Kit and the profiles were assessed on TapeStation and Fragment Analyzer.

#### Biotin enrichment and library preparation

Following gel extraction, equal volumes of gel-extracted DNA and pre-washed Dynabeads MyOne Streptavidin T1 (Invitrogen, 65601) in 2X Binding & Washing (BW) buffer (10 mM Tris-HCl pH 7.5 (Invitrogen, 15567027), 1 mM EDTA pH 8, 2M NaCl (Invitrogen, AM9760G)) were mixed to isolate biotin-enriched fragments. The bead–sample mixtures were incubated for 0.5–1 h at room temperature on a rotator at gentle speed. The bead-bound samples were washed twice with 1X TBW buffer (1X BW buffer, 0.1% Tween-20 (Sigma-Aldrich, P1379)) and resuspended in 100 µL water.

Library preparation was performed using NEBNext Ultra II DNA Library Preparation Kit for Illumina (NEB, 7645) according to manufacturer’s instructions with some modifications. Each bead-bound sample was split into 2 reactions before proceeding with end preparation and adaptor ligation. Briefly, 7 μL End Prep reaction buffer and 3 μL End Prep enzyme were added to each reaction, and the reactions were incubated for 45 min at 20°C and 600 rpm, followed by heat inactivation for 30 min at 65°C. Next, 2.5 μL NEB Illumina Adaptor, 30 μL UltraII ligation Master Mix and 1 μL Ligation Enhancer were added, and the reactions were incubated for 30 min at 20°C and 600 rpm. Subsequently, 3 μL USER enzyme was added and the reactions were further incubated for 30 min at 37°C and 600 rpm. The reactions were washed twice with 1X TBW buffer and resuspended in 57 μL water. Each reaction was further split into 2 reactions before proceeding with the indexing PCR for 10 cycles using Herculase II Fusion DNA Polymerase (Agilent Technologies, 600677). All reactions from the same original sample were pooled and purified using 0.9X MagBind TotalPure NGS beads (Omega Bio-Tek, M1378-01).

#### Pre-capture biotin clearing

Following indexing, the samples were subjected to an additional streptavidin binding step to remove residual biotin-containing fragments from previous steps. Briefly, equal volumes of indexed samples and pre-washed Dynabeads MyOne Streptavidin T1 in 2X BW buffer were mixed and the bead-sample mixtures were incubated at room temperature on a rotator at gentle speed. Next, the bead-sample mixtures were magnetized, cleared and the supernatants were transferred to fresh microcentrifuge tubes, followed by another round of clearing. The samples were purified using 0.9X MagBind TotalPure NGS beads. The final libraries were quantified using Qubit dsDNA BR Kit and the profiles were assessed on TapeStation.

#### Double capture enrichment

Biotinylated capture oligonucleotides (120 bp) were designed using a python-based oligonucleotide tool^77^ (https://oligo.readthedocs.io/en/latest/). Double capture enrichment was independently performed twice as described previously^78^ using KAPA HyperCapture Reagent Kit (Roche, 09075828001) with a few modifications. Briefly, 2 μg per replicate for each timepoint or condition was used as input. Samples were pooled and distributed across multiple hybridization tubes (maximum 12 μg per hybridization tube). Next, COT DNA was added and samples were dried by vaccum centrifugation at 45°C. Subsequently, Universal Enhancing Oligonucleotides, Hybridization buffer, Component H, and diluted capture oligonucleotides were added and mixed well. Following initial denaturation for 5 min at 47 °C, hybridization was performed at 47 °C for 65-72 h. Hybridized reactions were then incubated with pre-washed Dynabeads M-270 Streptavidin beads (Invitrogen, 65306) for 45 min at 47 °C and 600 rpm. The bead-bound samples were subsequently washed with a series of wash buffers and finally resuspended in water. Next, PCR amplification was performed for 8 cycles, followed by DNA purification using 1.8X MagBind TotalPure NGS beads. A second capture was carried out for 24 h using the same approach but with 7 PCR cycles.

#### Sequencing

The final libraries were quantified using Qubit dsDNA BR Kit and profiles were assessed on Fragment Analyzer. Sequencing was performed on the NextSeq 2000 Illumina platform using the long-read 300-bp paired end (600 cycles) XLEAP-SBS Kit (Illumina, 20100984).

### Improved Micro-Capture-C, Improved Tiled Micro-Capture-C and Improved Micro-Capture-C ultra

Improved Micro-Capture-C (iMCC) and Improved Tiled Micro-Capture-C (iTMCC) were performed for 3 biological replicates per timepoint. Improved Micro-Capture-C ultra (iMCCu) was performed for 3 biological replicates (with 2-4 technical replicates) per timepoint. For iMCC, iTMCC and iMCCu experiments, the same protocol was followed as described in the previous ‘mwMCC’ section with a few exceptions. Post-ligation iMCC libraries were size selected using 0.9X MagBind TotalPure NGS beads, followed by sonication for 250-280 sec on Covaris S220 Focused-ultrasonicator (10% Duty Factor, 200 Cycles per Burst, 175 Peak Incident Power). The samples were then subjected to biotin enrichment using Dynabeads MyOne Streptavidin T1, followed by sequencing library preparation using the NEBNext Ultra II DNA Library Preparation Kit for Illumina. Double capture enrichment for iMCC and iMCCultra, using 1.75 μg and 3 μg per replicate, respectively, was performed as described in the previous ‘mwMCC’ section. For iTMCC, tiled enrichment was performed as described previously^14^ using Twist Hybridization and Wash Kit (Twist Bioscience, 101025). Sequencing was performed on the NextSeq 1000/2000 Illumina platform using 150-bp paired end (300 cycles) XLEAP-SBS Kit (Illumina, 20100985).

### Analysis of MNase-based 3C data with the Capture MNase-based 3C Analysis Pipeline

A Snakemake pipeline named Capture MNase-based 3C Analysis Pipeline (C-MAP) was developed to process MNase-based 3C data. The pipeline starts with removing PCR duplicates with 100% sequence identity from paired-end FASTQ files using the BBMap dedupe.sh function (version 38.18). Adaptor sequences and low-quality end sequences are then trimmed using Trim Galore (version 0.6.10). Overlapping paired-end reads are merged with FLASH (version 1.2.11) to reconstruct the original DNA fragments. Both successfully merged reads (flashed reads) and unmerged paired-end reads (unflashed reads) are aligned to the reference genome (hg38) using BWA-MEM (version 0.7.12). The BWA-MEM algorithm, which performs local alignment and can generate separate alignments for distinct segments of a query sequence, is used to resolve ligation junctions in chimeric reads. Parameters are adjusted to increase sensitivity (-k 18 -w 1 -L 0 -r 1 -T 18), enabling the detection of multiple shorter alignments. After alignment, unflashed reads are merged if the genomic distance between their paired ends is shorter than the allowed gap threshold (default: 200 bp). Segments overlapping with viewpoints are designated as capture sites, while the remaining segments are designated as reporters. Chimeric reads are separated based on their capture sites. Reads lacking capture sites or ligation junctions are filtered out. PCR duplicates, defined as reads with identical or near-identical alignments (allowing a 2 bp wobble at segment ends), are removed.

To generate pair-wise interaction profiles (for e.g., (i)MCC data), reads with a unique capture site and at least one reporter are retained. Viewpoint-proximal reporters are excluded, and coverage tracks for the remaining reporters are generated in bigWig format using the bedtools genomecov function (version 2.31.1). The interaction profiles are normalized for an equal number of cis reporters (reporters from the same chromosome as the viewpoint). Plots of ligation junctions enable footprinting of DNA-binding proteins. (i)MCC footprinting is visualized in coverage tracks for the ligation junctions, which are generated using bedtools genomecov function and normalized for an equal number of ligation junctions in cis.

To generate contact matrices (for e.g., (i)TMCC, RCMC, or (i)MCCu data), reads with at least one ligation junction located within the tiled regions are retained. For each tiled region, the ligation junctions are binned, and a raw contact matrix is computed with cooler cload function (version 0.10.2), followed by balancing with the cooler balance function. Contact matrices are plotted in Python (version 3.9.18) using cooler (version 0.10.2), matplotlib (version 3.8.2) and cooltools (version 0.7.1) packages.

To generate viewpoint-specific 3D contact matrices (for e.g., mwMCC data), reads with a unique capture site and at least two cis reporters are retained. Multi-way interactions are visualized by fixing the viewpoint and binning the remaining pairs of cis reporters from each read into a contact matrix with a custom Python script. Viewpoint-proximal interactions are excluded and the matrices are further normalized for the total multi-way interaction counts within a 2 Mb region surrounding the viewpoint.

### Benchmarking of pair-wise MNase-based methods

Both conventional MCC and iMCC data were analyzed using C-MAP in ‘capture’ mode with the same settings. The proportion of valid ligation junctions was defined as the ratio of chimeric reads with at least one ligation junction over all high-confidence mapped reads (MAPQ ≥ 20). The capture efficiency was defined as the ratio of reads with at least one capture site over all high-confidence mapped reads. To allow for a direct comparison of valid cis interactions, conventional MCC and iMCC data were downsampled to equal sequencing depth, equivalent to an average of 10k reads per viewpoint.

TMCC, RCMC^16^, iTMCC, MCCu^17^, and iMCCu data were analyzed using C-MAP in ‘tiled’ mode with comparable settings. TMCC, RCMC and iTMCC were first downsampled to an average of 4M reads per targeted Mb. For every targeted region across every sample, the number of valid cis interaction pairs per targeted Mb was calculated and compared. Similarly, MCCu and iMCCu were downsampled to 250k reads per targeted kb and the number of valid cis interaction pairs per targeted kb was calculated and compared.

### Analysis of Tri-C data

Tri-C data^35^ were analyzed using the CapCruncher pipeline (version 0.2.3) in capture mode, as reported previously^35^. A custom script was used to extract reads with two or more cis reporters. These reads were then used to calculate multi-way interaction counts between restriction fragments for each viewpoint. The interaction counts were binned and corrected for the number of restriction fragments present in each bin. The matrices were further normalized for the total multi-way interaction counts within a 2 Mb region (which was corrected for the number of restriction fragments present in the region) surrounding the viewpoint.

### Quantification of multi-way chromatin interactions

To systematically quantify multi-way interactions detected by mwMCC, mwMCC data were first analyzed as pair-wise MCC data in ‘capture’ mode with C-MAP. MCC peaks were called and annotated as described previously^35^. Next, candidate multi-way interactions were identified by examining all combinations of MCC peaks interacting with the viewpoint. To ensure robust quantification, all MCC peaks were extended by ± 250 bp and merged if their distance was less than 500 bp. Peaks within ± 2 kb of the viewpoint were excluded. Candidate multi-way interactions were identified from the remaining peaks. The interaction frequency of each candidate multi-way interaction was quantified at 100 bp resolution by extracting the corresponding contact submatrix and calculating the mean of the upper 50% of non-zero bins, which was corrected by a scaling factor. Candidate hubs were filtered to retain only those with interaction frequencies above a threshold of 1.2–1.5 (depending on the viewpoint) at a timepoint at which the target locus was actively expressed.

Hubs were grouped into enhancer-promoter (EP) hubs, CTCF hubs, and mixed hubs based on the cis-regulatory elements involved in the multi-way interactions. The interaction frequencies of hubs across different groups were measured during cellular differentiation and following CTCF depletion. A two-sided paired Wilcoxon signed-rank test was used to compare the interaction frequencies between the two conditions.

### Assessment of synergy of multi-way interactions

To assess the synergistic nature of multi-way interactions between a viewpoint (*V*) and two genomic loci (*X* and *Y*), we studied the deviation of the observed multi-way interaction frequency from the expectation under an independent pair-wise interaction model. Under the independent interaction model, the expected multi-way interaction is fully explained by the pair-wise contributions:

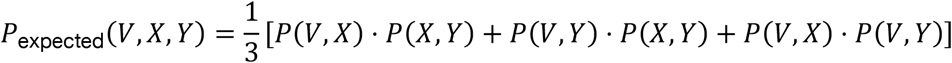

where *P*_expected_(*V, X, Y*) represents the expected absolute multi-way looping probability assuming no high-order synergy of a multi-way interaction between the viewpoint (*V)* and the two genomic loci (*X* and *Y*), while *P*(*V, X*), *P*(*V, Y*) and *P*(*X, Y*) represent the absolute pair-wise looping probabilities.

In practice, for a given genomic region and viewpoint, pair-wise 3C interaction frequencies were extracted from the balanced iTMCC contact matrix and the expected multi-way interaction frequency was simulated based on the probability distribution of pair-wise interaction frequencies. Both the observed and expected matrices of multi-way interaction frequencies were normalized to have equal total sums to enable direct comparison. Contact synergies were quantified as the enrichment of observed over expected interaction frequencies. Synergies among multi-way interactions involving cis-regulatory elements or randomly sampled loci were measured at 500 bp resolution. To ensure robust quantification, cis-regulatory elements, identified by MCC peaks, were extended by ± 500 bp and merged if separated by less than 1000 bp. Randomly sampled loci were defined to match the size of the cis-regulatory elements.

### Estimation of absolute multi-way looping probabilities

To estimate the absolute looping probability of a multi-way interaction (*V,X,Y*), we first computed absolute looping probabilities of pair-wise interactions (*V,X*), (*V,Y*), and (*X,Y*) using the AbLE (Absolute Looping Estimator) framework^50^. AbLE scores were derived from balanced iTMCC contact matrices via the looptools.py module^50^. These scores were converted to absolute pair-wise interaction probabilities using a previously determined calibration factor of 0.186^50^. Absolute multi-way looping probabilities were then estimated by incorporating derived contact synergies into absolute pair-wise looping probabilities.

The observed multi-way interaction frequency detected at genomic bin *i*, denoted *O*_*i*_(*VXY*), is proportional to the absolute multi-way looping probability *P*_*i*_(*V, X, Y*):

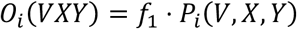

where *f*_1_ is an unknown scaling factor.

According to how we quantified contact synergies in the last section, the expected multi-way interaction frequency simulated at genomic bin *i*, denoted *S*_*i*_(*VXY*), is proportional to the average of products of absolute pair-wise looping probabilities *P*_*i*_(*V, X*), *P*_*i*_(*V, Y*) and *P*_*i*_(*X, Y*):

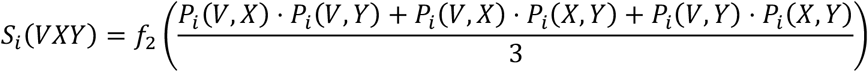

where *f*_2_ is an unknown scaling factor.

*O*_*i*_(*VXY*) and *S*_*i*_(*VXY*) were normalized to have equal total sums:

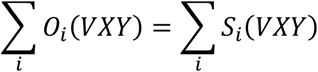

The contact synergy at genomic bin *i*, denoted *Synergy*_*i*_ was defined as:

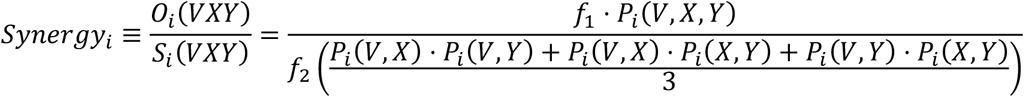

Therefore, the absolute multi-way looping probability can be computed as:

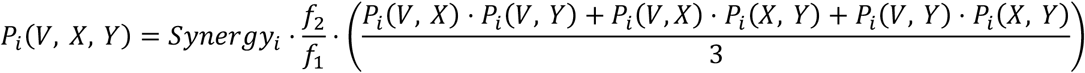

Here, 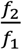 is derived by summing both *O*_*i*_(*VXY*) and *S*_*i*_(*VXY*) over *i*, applying the normalization equation, and solving the equation for 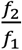, giving

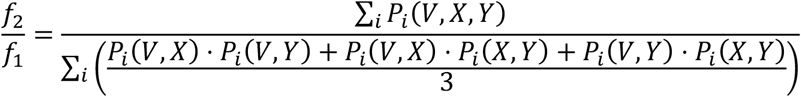

The ratio 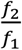 equals 1 when all pair-wise interactions across a large genomic distance are independent; it is greater than 1 when interactions are positively correlated; it is less than 1 when they are negatively correlated.

We assume 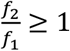 because, across the large genomic scale that we normalize *O* (*VXY*) and *S*_*i*_(*VXY*) over, we do not expect a general repulsive effect between pair-wise interactions from known chromatin organization principles. Therefore, we estimated the lower bound of absolute multi-way looping probability with

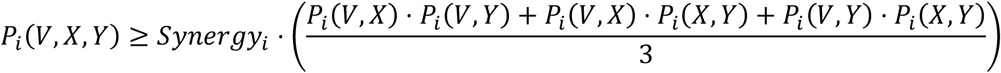

### Plotting of contact sequence reconstruction maps

Contact sequence reconstruction maps were generated following the same approach as reported previously^17^. iMCCu data were analyzed with the pipeline in ‘tiled’ mode to resolve the position and directionality of ligation junctions. Ligation junctions from all replicates were merged. To construct a matrix presenting potential protein binding sites, each ligation junction was represented as a rotated 18 x 12 bp rectangle, with the rectangle’s orientation reflecting the direction in which the DNA fragments were ligated. The matrix was normalized by the total number of ligation junctions detected. Transcription factor binding sites were annotated with JASPAR (2026 release).

### Quantification of protein occupancy at hubs

The occupancy of MED26 at iMCC peaks was quantified by extracting the coverage information from the corresponding bigWig files and calculating the average of the highest 70% of base pair coverage values across the iMCC peaks as described previously^35^. Chromatin hubs were defined as pairs of interacting regions (anchors) that contact the viewpoint. Protein occupancy for each anchor was calculated as the width-weighted mean of protein signal across overlapping MCC peaks. Hub-level occupancy was then defined as the sum of the occupancy values of the two anchors, representing the total protein enrichment across both interacting regions. Spearman rank correlation coefficient was computed to investigate the relationship between the occupancy of MED26 and contact synergy at hubs during lymphoid-to-myeloid differentiation.

### Public data analysis

ATAC-seq^38^ and ChIP-seq data for CTCF^38^ and MED26^35^ were analyzed using the NGseqBasic pipeline^79^. Published Region Capture Micro-C (RCMC)^16^ and Micro Capture-C ultra (MCCultra)^17^ datasets were re-analyzed using our custom C-MAP pipeline.

## ACKNOWLEDGEMENTS

We would like to thank Patrick Cramer for infrastructure support; Luca Giorgetti and Mattia Ubertini for advice on the calculation of absolute interaction probabilities; Kerstin Maier, Myriam Rohm, Petra Rus, Lea Siegmund, and Robin Walz for experimental support; and all members of the Oudelaar group for discussions and feedback. This work was supported by the Max Planck Society (AMO); the European Research Council (Starter Grant 3D-REG 101115401, AMO); the Deutsche Forschungsgemeinschaft (DFG) via SFB 1565 (project 469281184/P02, AMO); the PhD program “Genome Science”—International Max Planck Research School at the Georg August University Göttingen (SR and MK); and the MSc/PhD program “Molecular Biology”— International Max Planck Research School at the Georg August University Göttingen (YZ).

## AUTHOR CONTRIBUTIONS STATEMENT

S.R. and Y.Z. contributed equally and are co-first authors of this article. The order of the co-first authors is based on alphabetic order and the co-first authors are free to alter this on their CVs. S.R. developed mwMCC, carried out most of the experiments, performed basic data analyses, prepared the majority of the figures, and wrote the manuscript. Y.Z. developed C-MAP, performed the majority of the bioinformatic analyses and mathematical modelling, contributed to figure preparation, and wrote the manuscript. M.A.K. performed Tri-C experiments. I.M.F. provided important conceptual advice for the development of mwMCC. J.S. supervised the development of C-MAP, bioinformatic analyses, and mathematical modelling. A.M.O. conceived and supervised the project, acquired funding, and wrote the manuscript. All authors provided feedback on the manuscript.

## EXTENDED DATA FIGURES

**Extended Data Figure 1:**
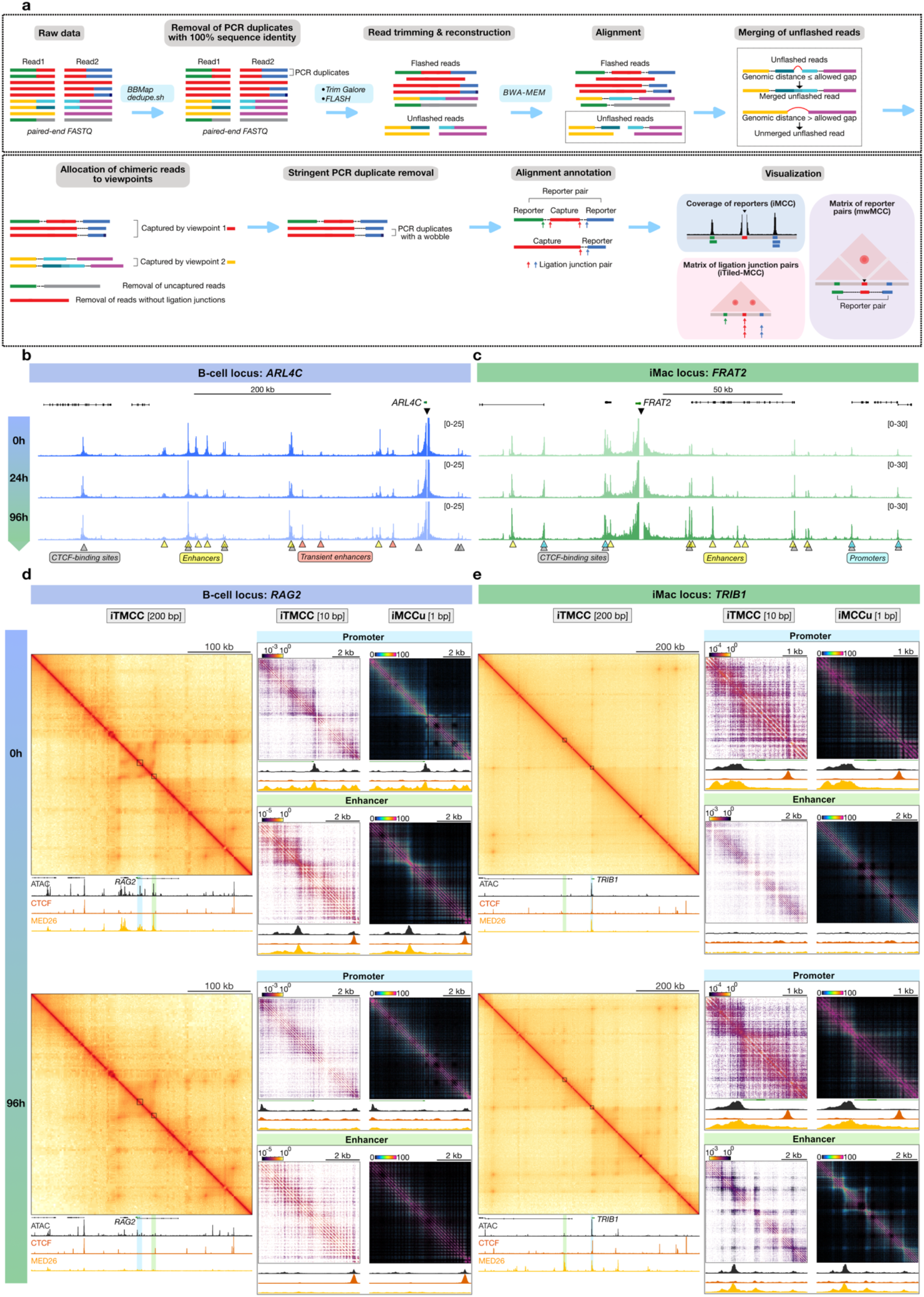
Improved targeted MNase-based 3C experimental procedure and analysis pipeline. **a**, Outline of C-MAP, an analysis pipeline for targeted MNase-based 3C data. **b**, iMCC interaction profiles of the ARL4C and FRAT2 promoters through lymphoid-to-myeloid transdifferentiation. Black triangles at the top indicate the viewpoint; yellow, red, blue, and grey triangles indicate enhancers, transient enhancers, promoters, and CTCF-binding sites, respectively. Coordinates (hg38): ARL4C locus, chr2:233908830-234563849; FRAT2 locus, chr2:234131765-234169572. **d**, iTMCC large-scale contact matrices (200 bp resolution), iTMCC nano-scale contact matrices (10 bp resolution), and iMCCu contact matrices (1 bp resolution) of the RAG2 locus through lymphoid-to-myeloid transdifferentiation. Small black squares and shadings in blue and green in the large-scale iTMCC contact matrices on the left indicate regions shown at higher resolution on the right. Profiles below show ATAC-seq, CTCF ChIP-seq, and MED26 ChIPmentation. Axes scales: large-scale, 0-2130 (ATAC-seq), 0-10994 (CTCF), 0-2527 (MED26); promoter nano-scale, 0-1718 (ATAC-seq), 0-1000 (CTCF), 0-1233 (MED26); enhancer nano-scale, 0-1826 (ATAC-seq), 0-2360 (CTCF), 0-2831 (MED26). Coordinates (hg38) as in **1f** and **1g. e**, iTMCC large-scale contact matrices (200 bp resolution), iTMCC nano-scale contact matrices (10 bp resolution), and iMCCu contact matrices (1 bp resolution) of the TRIB1 locus through lymphoid-to-myeloid transdifferentiation, as in **d**. Axes scales: large-scale, 0-5963 (ATAC-seq), 0-6348 (CTCF), 0-4182 (MED26); promoter nano-scale, 0-8268 (ATAC-seq), 0-9066 (CTCF), 0-3320 (MED26); enhancer nano-scale, 0-1437 (ATAC-seq), 0-1500 (CTCF), 0-5546 (MED26). Coordinates (hg38) as in **1f** and **1g**.

**Extended Data Figure 2:**
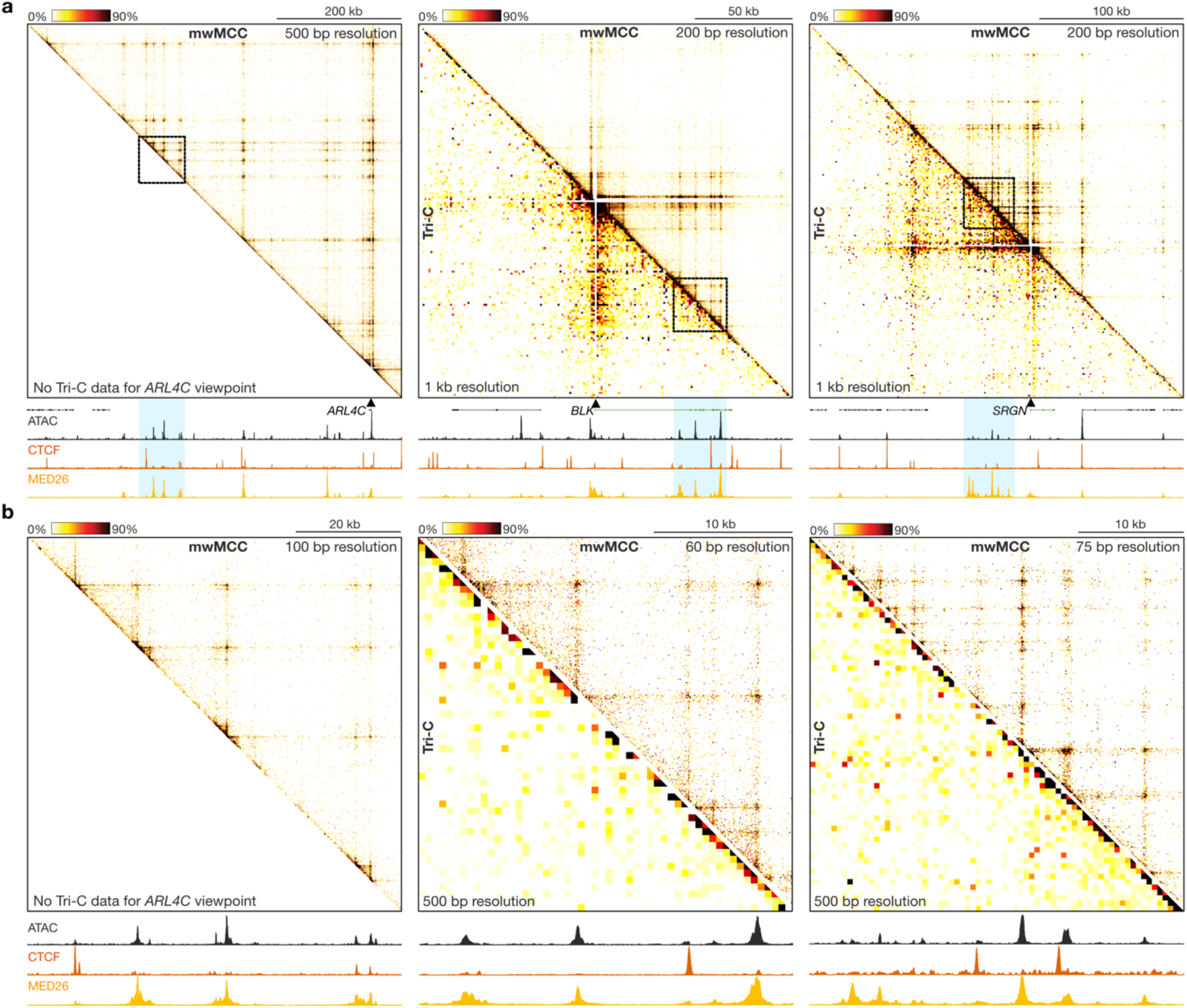
Benchmarking of multi-way Micro-Capture-C. **a**, Comparison of large-scale mwMCC (top-right; 200-500 bp resolution) and Tri-C (bottom-left; 1 kb resolution) data from promoter viewpoints in BlaER1 cells at 0h (ARL4C and BLK) and 96h (SRGN) post-differentiation induction, as in **2b**. Tri-C data for the ARL4C promoter is missing, because it is not possible to generate due to the large size of the corresponding NlaIII restriction fragment (>1.5 kb). Black squares and blue shadings indicate regions shown at higher resolution in **b**. Profiles below show ATAC-seq, CTCF ChIP-seq, and MED26 ChIPmentation. Axes scales and coordinates (hg38 as in **2b. b**, Comparison of nano-scale mwMCC (top-right; 60-100 bp resolution) and Tri-C (bottom-left; 500 bp resolution) data, as in **a**. Axes scales and coordinates (hg38 as in **2b**.

**Extended Data Figure 3:**
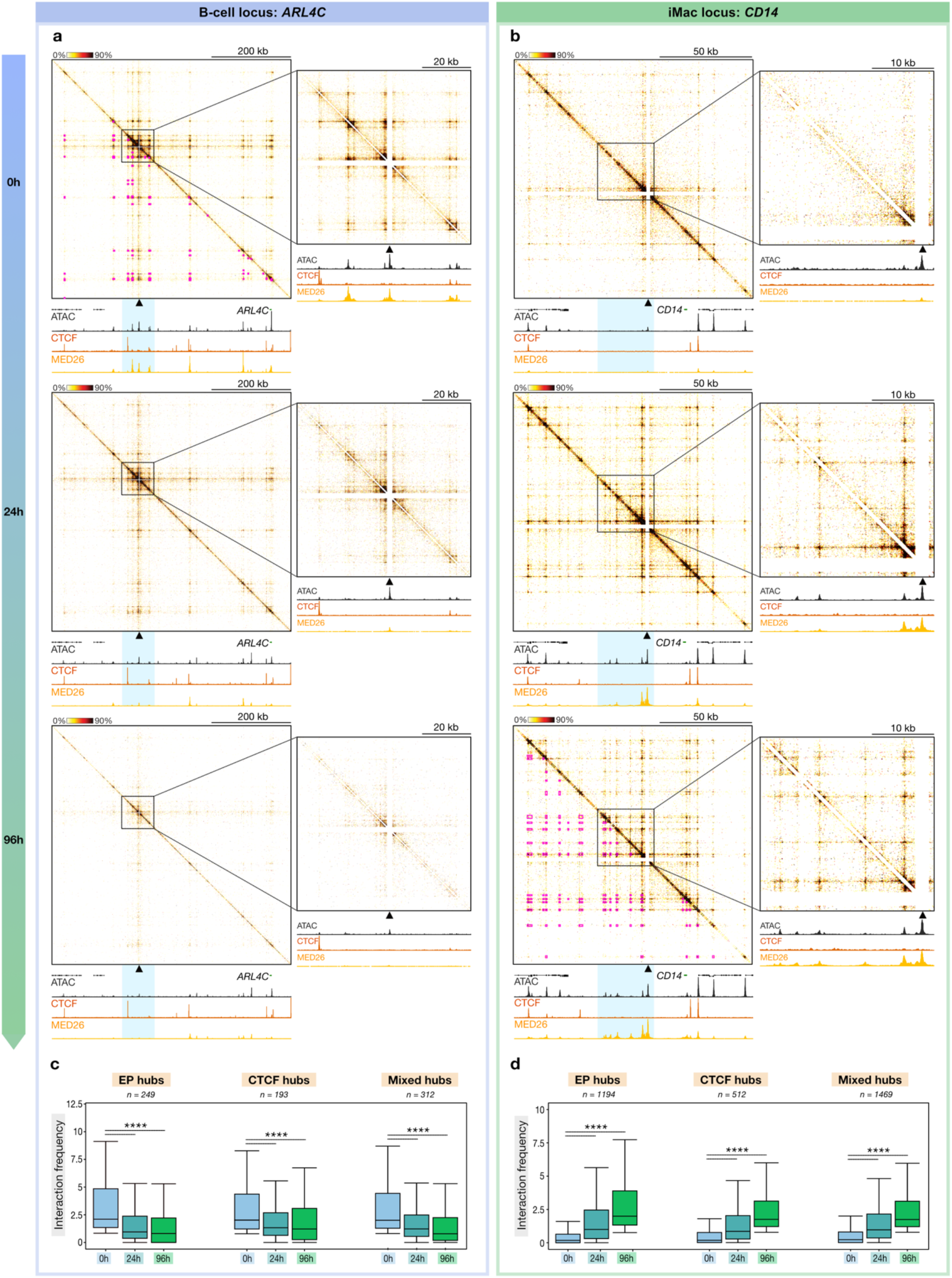
Dynamic formation and dissolution of large-scale and nano-scale chromatin hubs during cellular differentiation. **a**, mwMCC contact matrices of the ARL4C locus (enhancer viewpoint; 500/100 bp resolution) through lymphoid-to-myeloid transdifferentiation, as in **2b**. Pink squares indicate loop calls. Axes scales and coordinates (hg38) as in **3a. b**, mwMCC contact matrices of the CD14 locus (enhancer viewpoint; 200/75 bp resolution) through lymphoid-to-myeloid transdifferentiation, as in **a**. Axes scales and coordinates (hg38) as in **3b. c**, Multi-way interaction frequencies of enhancers of B-cell-specific genes with enhancers and promoters (EP hubs), CTCF-binding sites (CTCF hubs), and a mixture of enhancers, promoters, and CTCF-binding sites (Mixed hubs), as in **3c**. Asterisks indicate statistical significance (two-sided paired Wilcoxon signed-rank test, 96h versus 0h). P-values: EP hubs, P < 2.2 × 10^-16^; CTCF hubs, P = 5.9 × 10^-1^; Mixed hubs, P < 2.2 × 10^-16^. The number of data points (n) in each category is shown above the graph. Data are derived from three biological replicates. **d**, Multi-way interaction frequencies of enhancers of iMac-specific genes, as in **c**. P-values: EP hubs, P < 2.2 × 10^-16^; CTCF hubs, P < 2.2 × 10^-16^; Mixed hubs, P < 2.2 × 10^-16^.

**Extended Data Figure 4:**
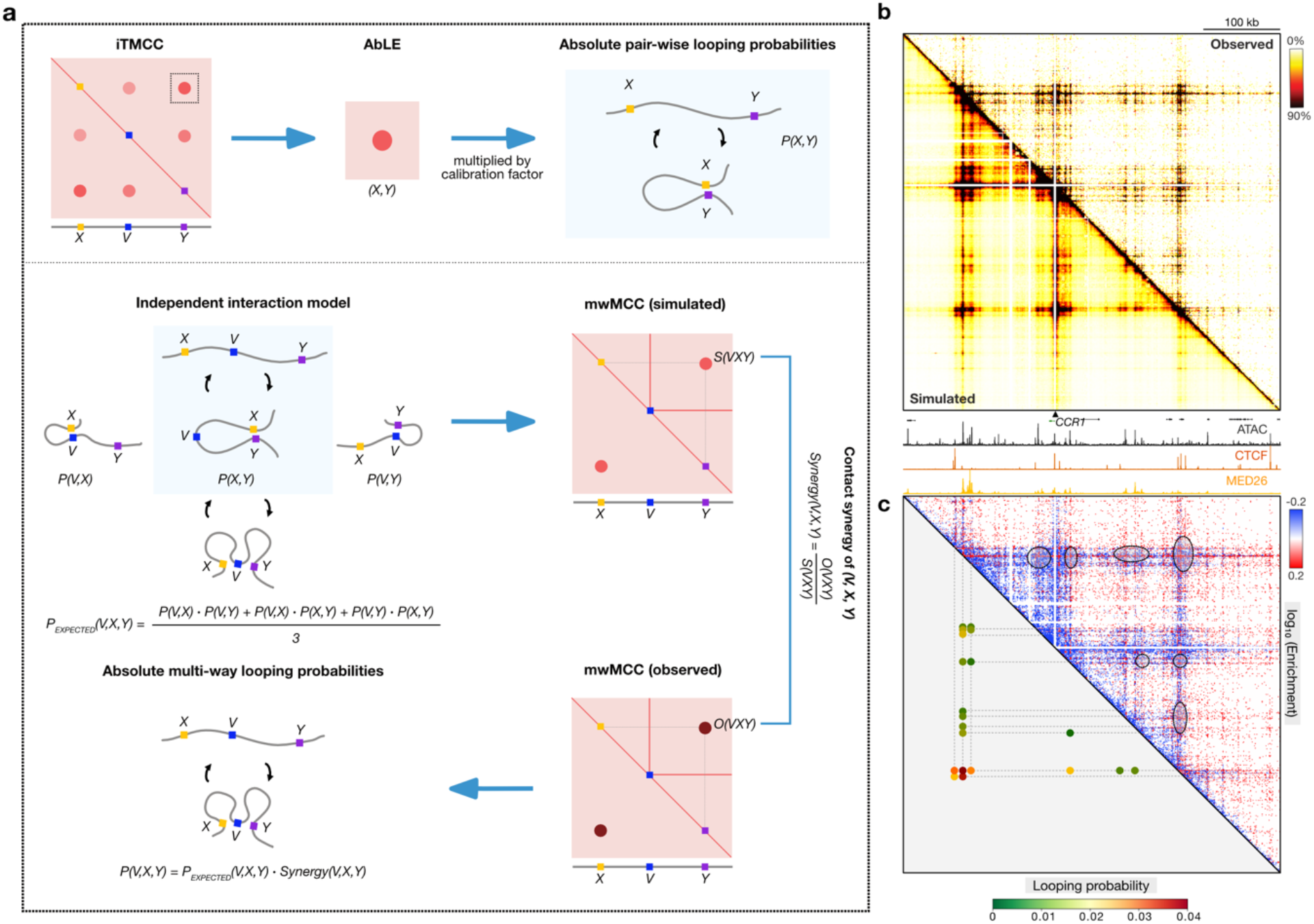
Synergy and absolute quantification of multi-way interactions. **a**, Outline of workflow for calculations of contact synergies and absolute multi-way looping probabilities. iTMCC loops are quantified using the Absolute Looping Estimator (AbLE)^50^. AbLE scores are converted to absolute pair-wise looping probabilities using the calibration factor 0.186^50^. An independent interaction model assumes independent co-occurrence of pair-wise interactions and no synergy between pair-wise interactions. Based on this model, expected multi-way looping probabilities are calculated and mwMCC contact matrices are simulated. Contact synergies are assessed by the deviation of the observed multi-way contact frequencies from the expectation under the independent interaction model, defined as the enrichment of observed over simulated multi-way contact frequencies. Absolute multi-way looping probabilities are estimated by incorporating contact synergies into expected multi-way looping probabilities. **b**, 3D contact matrices of the CCR1 locus (promoter viewpoint; 1000 bp resolution) in BLaER1 cells at 96h post-differentiation induction, derived from mwMCC data (top-right) or simulated based on an independent interaction model (bottom-left), as in **2b**. Axes scales: 0-875 (ATAC-seq), 0-7000 (CTCF), 0-5244 (MED26). Coordinates (hg38): chr3:46011998-46501998. **c**, 3D contact matrices showing enrichment of observed over simulated multi-way contact frequencies (top-right; 1000 bp resolution) and estimated absolute looping probabilities (bottom-left), as in **b**.

**Extended Data Figure 5:**
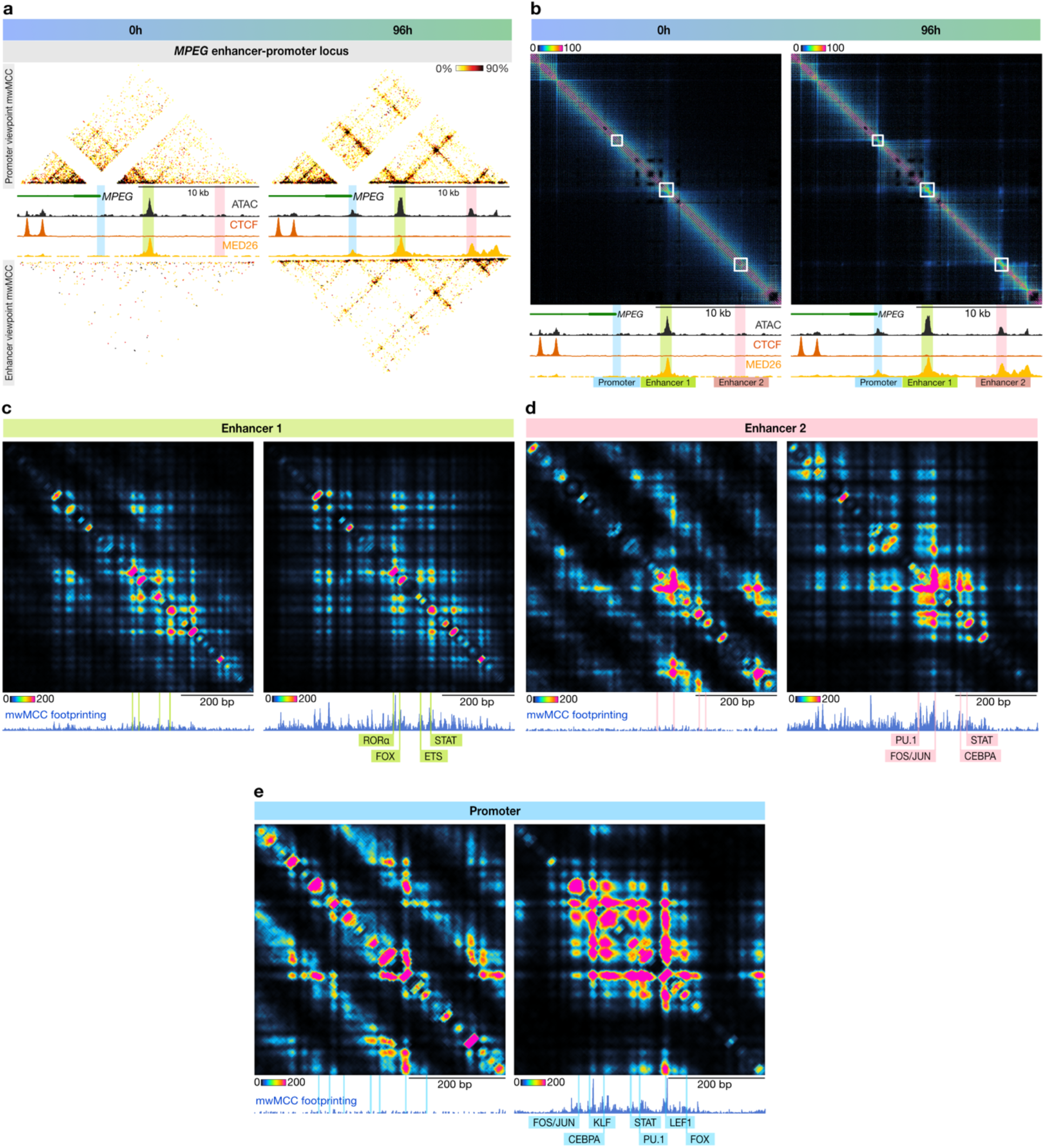
Footprinting analysis of nano-scale chromatin hubs. **a**, mwMCC contact matrices of the MPEG locus (100 bp resolution) in BLaER1 cells at 0h (left) and 96h (right) post-differentiation induction, as in **2b**. Axes scales: 0h, 0-1433 (ATAC-seq), 0-5142 (CTCF), 0-2832 (MED26); 96h, 0-1433 (ATAC-seq), 0-5142 (CTCF), 0-2832 (MED26). Coordinates (hg38): chr11:59206032-59226040. **b**, iMCCu contact matrices of the MPEG locus (1 bp resolution), as in **a. c-e**, Contact sequence reconstruction matrix, mwMCC footprinting, and transcription factor motif annotation of Enhancer 1 of the MPEG locus (**c**), Enhancer 2 of the MPEG locus (**d**), and the promoter of the MPEG locus (**e**), in BLaER1 cells at 0h (left) and 96h (right) post-differentiation induction.

## SUPPLEMENTARY NOTE 1

In this supplementary note, we discuss specific steps of the i(T)MCC(u)/mwMCC procedures that we systematically optimized to provide transparency on the aspects that improve the efficiency of the methods.

Note: Most values reported here were obtained during early optimization stages and therefore represent underestimates of the quality control parameters. The parameters reported in the main article represent the final optimized versions of the protocols.

### Crosslinking and permeabilization

#### Input material

In this study, we used 5 million fixed cells per replicate for all experiments. However, this protocol has also been successfully used with lower cell inputs of 2-3 million fixed cells per replicate.

#### Single vs. dual crosslinking

We did not notice specific improvements in key quality control parameters when performing dual crosslinking (30 min disuccinimidyl glutarate (DSG) + 10 min formaldehyde (FA)) compared to single crosslinking (10 min FA). We did, however, consistently notice a reduction in *cis*-to-*trans* ratios with dual crosslinking (**SN_1A-B**).

**SN_1.**
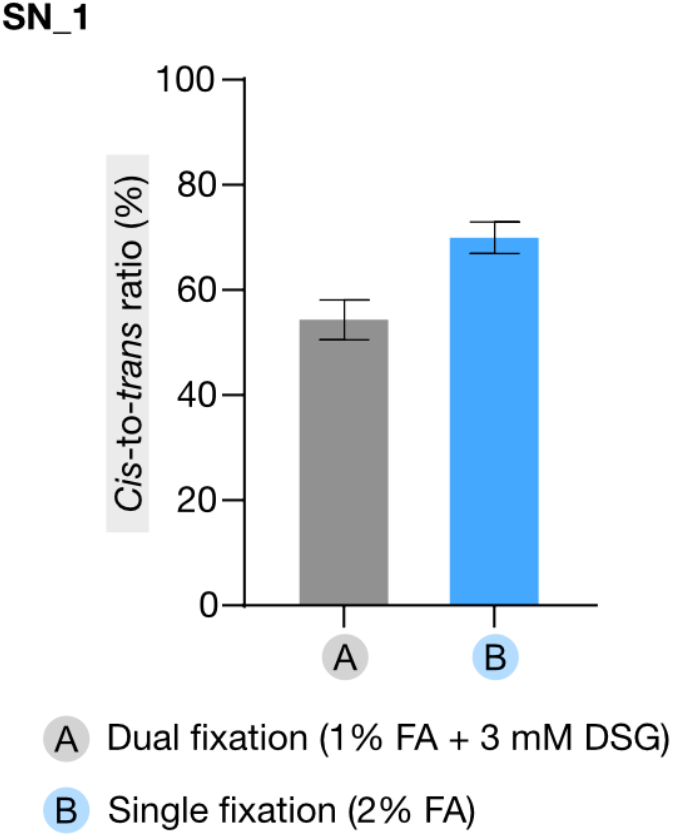

### iMCC library generation

#### Concanavalin A beads

We found that the use of Concanavalin A (ConA) beads to bind crosslinked and permeabilized cells prior to MNase digestion, and throughout all intermediate steps up to ligation, increases library yield post-ligation 2-3 fold (**SN_2A-C**). This increases overall library complexity, which is especially critical for the multi-way module. Additionally, we observed that implementation of ConA beads improves *cis*-to-*trans* ratios. In our hands, when ConA beads are used, a higher amount of MNase is required to digest the chromatin compared to standard conditions. We tested whether binding the samples post-digestion would work equally well. However, this led to less optimal digestion and ligation profiles, as well as lower post-ligation yields, although the yield was still higher compared to samples that were not bound to ConA beads at all (**SN_2B**).

**SN_2.**
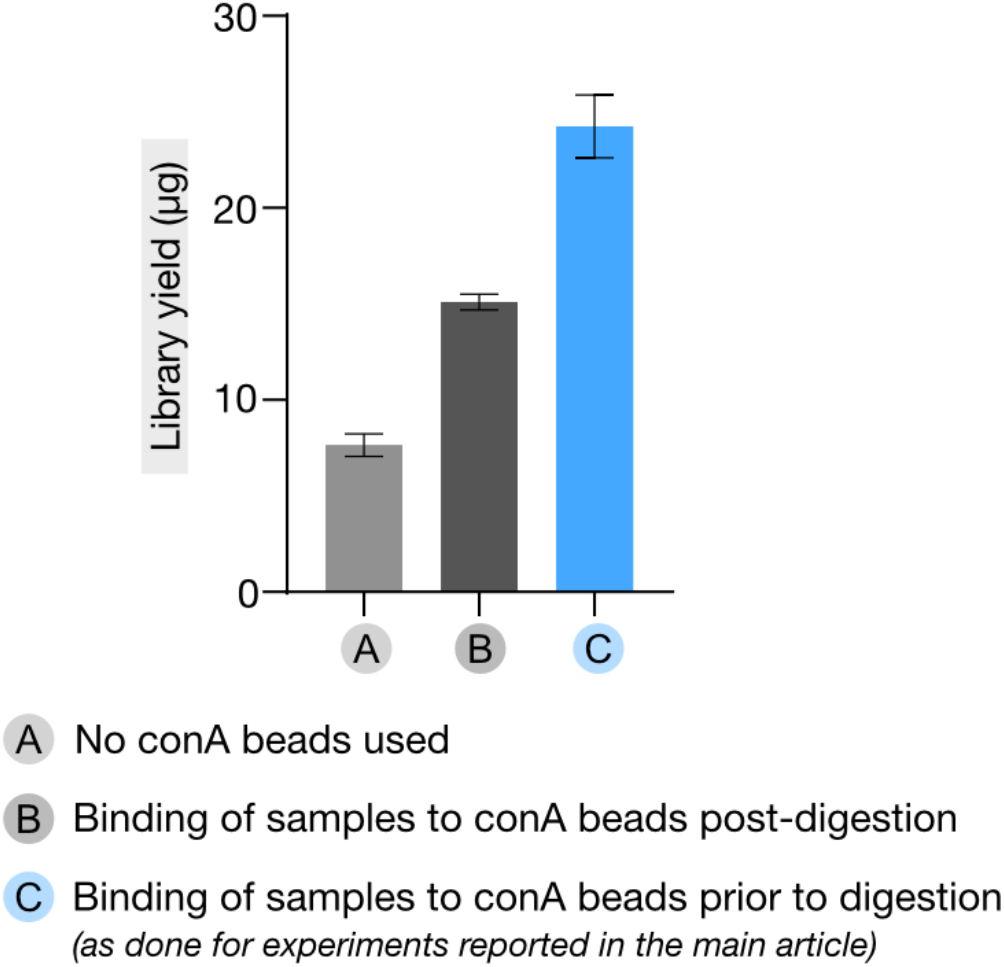

#### Digestion

Although biotin labelling allows for enrichment of valid ligation junctions, it is not possible to specifically enrich for fragments with two valid junctions for mwMCC. We found that a high degree of digestion (**Supplementary Note 2**) results in higher proportions of valid pair-wise and multi-way ligation junctions (even when ligation appears relatively inefficient in DNA profiles).

#### End-prep/labeling/ligation

We found that extending the duration of the reactions for end preparation with T4 PNK and Klenow, end labeling with dNTPs, and ligation with T4 ligase improves library quality. In our hands, increasing the concentrations of T4 PNK and Klenow did not have any effect.

#### DNA extraction

We found that extraction of DNA post-ligation and reverse cross-linking with the DNeasy Blood and Tissue Kit (Qiagen, 69504) results in higher yields compared to conventional phenol-chloroform extraction (**SN_3A-B**).

**SN_3.**
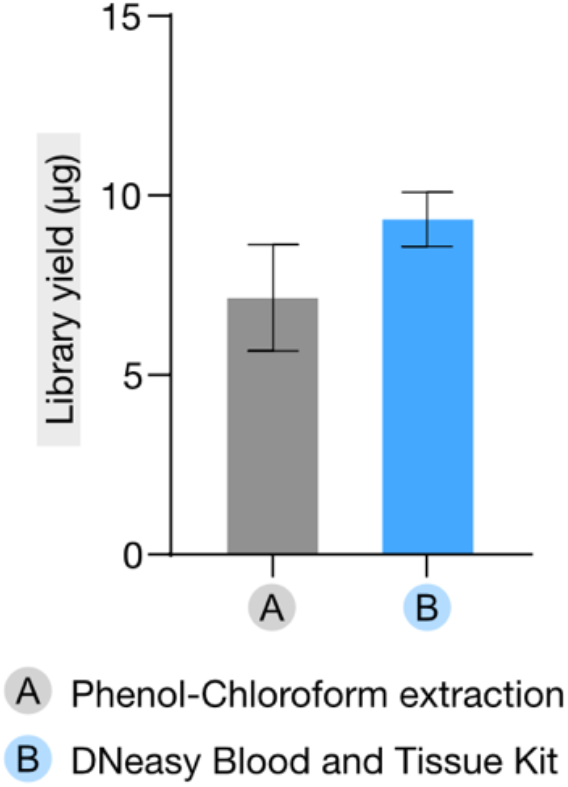

### mwMCC Size selection

We tested several sonication conditions (20-70 s) and strategies for double-sided size selection with Solid-Phase Reversible Immobilization (SPRI) beads to enrich libraries for relatively large fragments (>500) that allow for multi-way analysis. However, unlike for the pair-wise module, sonication and double-sided size selection with SPRI beads were not sufficiently precise and therefore did not allow for the generation of libraries with the desired fragment size distribution without compromising library yield and complexity. We found that gel extraction allows for more efficient size selection and results in the highest proportion of reads with two ligation junctions or more (**SN_4A-C**). We further refined the gel extraction procedure by testing different gel percentages (0.8-1.5%), input loading, fragment distribution ranges, and extraction methods to maximize the output post-gel extraction and retain optimal library complexity. With these optimizations, mwMCC generates data with similar proportions of multiple ligation junctions as the Tri-C method, even though Tri-C is based on the inherently efficient ligation of sticky ends after digestion with the restriction enzyme NlaIII (**SN_4D-E**).

**SN_4.**
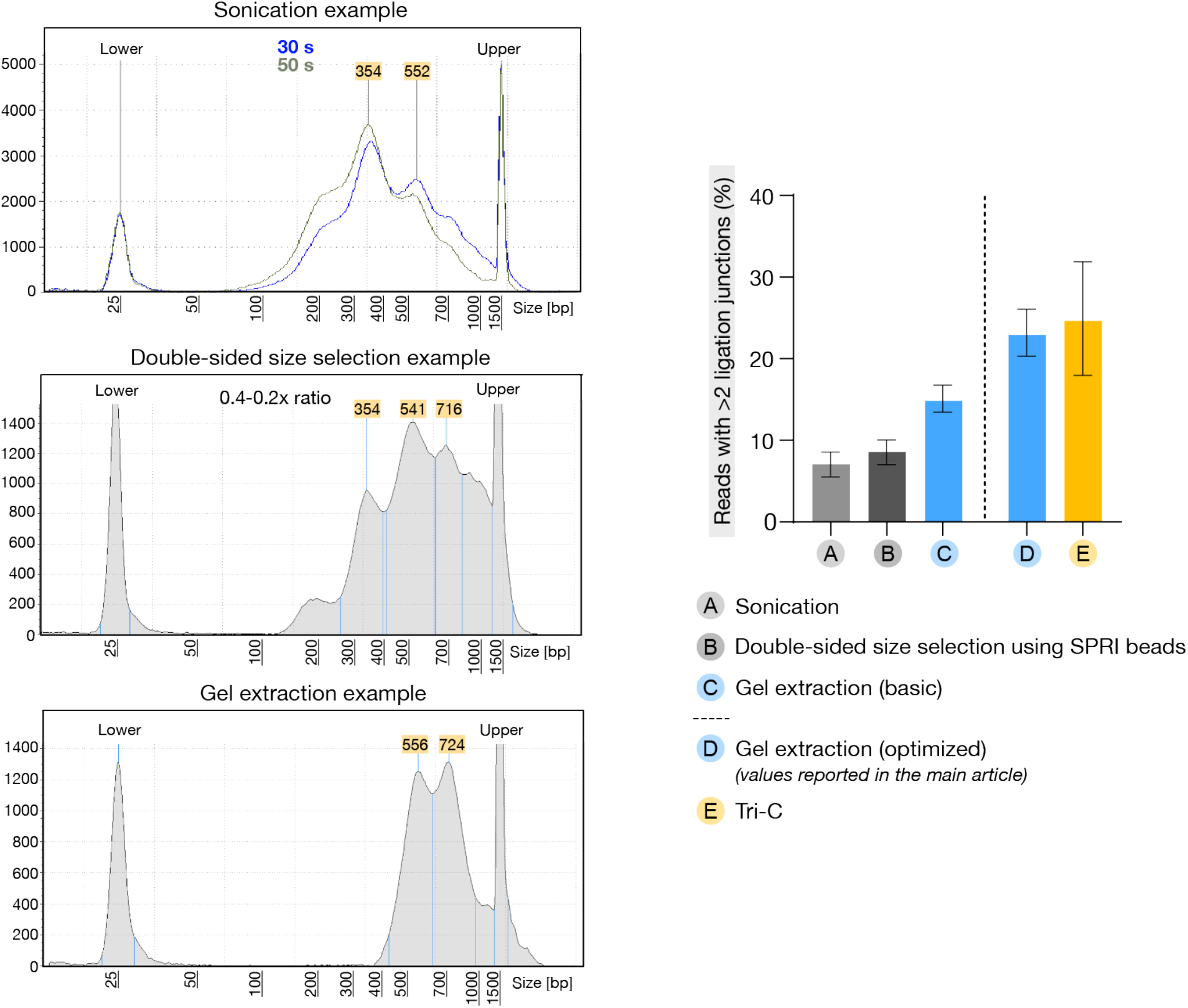

### Adaptor ligation and indexing PCR

It is often useful to obtain relatively high post-indexing PCR library yields to allow for re-use of libraries with multiple capture-oligonucleotide pools (to boost complexity or allow for enrichment for additional regions of interest). In the optimized procedure, biotin-containing fragments are enriched using streptavidin T1 beads prior to library preparation. However, streptavidin beads can interfere with the downstream enzymatic adaptor ligation and indexing PCR and thereby reduce library yield **(SN_5A)**. By splitting the reactions for both the adaptor ligation and indexing PCR reactions, we dilute the beads at each stage and thus minimize their interference. This results in approximately 6-fold higher concentrations of indexed libraries from the same pre-pulldown input (**SN_5C**), and outperformed splitting only at the indexing PCR step (**SN_5B**). Additionally, consistent with previous reports^16^, we find that streptavidin T1 dynabeads beads outperform C1 dynabeads beads, yielding approximately 3-fold more indexed library (**SN_5D-E**).

**SN_5.**
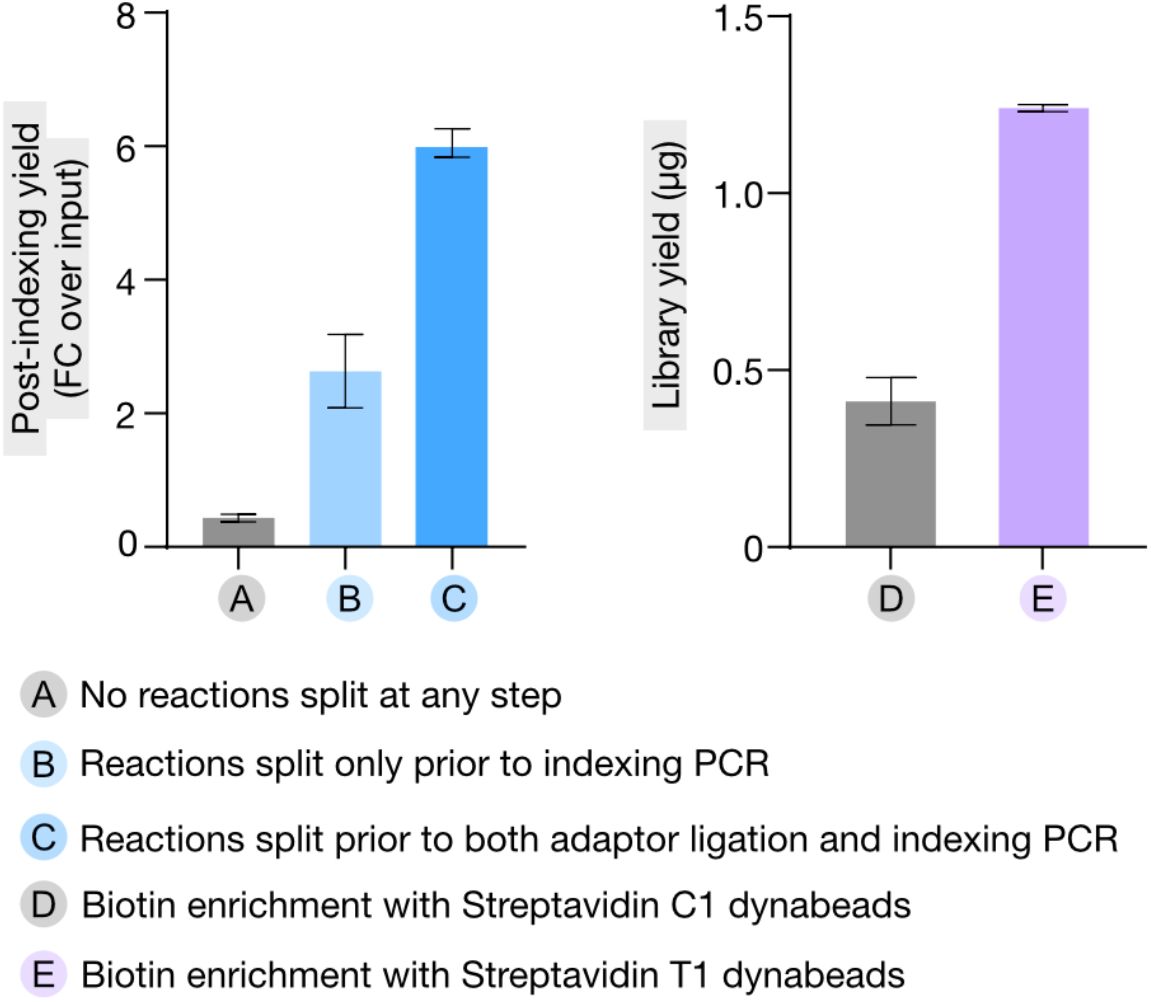

### Oligonucleotide capture

For both the pair-wise and multi-way modules, we noticed that complexity is never saturated with just one round of oligonucleotide capture. Setting up multiple captures with the same indexed libraries boost complexity by 1.6-fold on average (**SN_6A-B**). We also tested using 2x more probes in a single oligonucleotide capture but this did not significantly improve complexity.

**SN_6.**
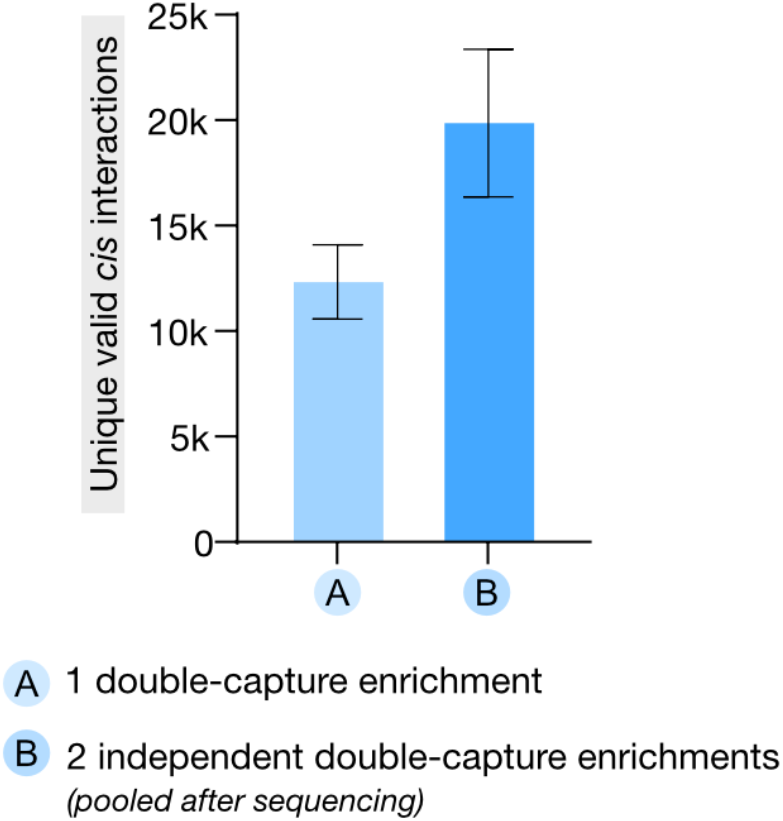

### Capture efficiency

We define capture efficiency as the percentage of reads that contain chimeric fragments that overlap a targeted viewpoint. In our hands, the incorporation of biotin labeling of digested ends results in lower capture efficiency, likely due to carry-over of biotinylated contaminants that interfere with the biotinylated capture oligonucleotide enrichment. The initial average capture efficiencies we obtained with the improved pair-wise and multi-way procedures were 42.8% (**SN_7G**) and 11.6% (**SN_7A**), respectively. We tried different strategies to improve the capture efficiency, focusing on the multi-way module: 1) an extra streptavidin-binding step to remove biotinylated contaminants from the indexed library pools prior to oligonucleotide capture; 2) 5x higher concentration of capture probes; 3) a third round of capture enrichment, as it has been shown that additional rounds of capture enrichment improve capture efficiency^80^. All conditions improved the capture efficiency. An extra streptavidin-binding step and 5x higher concentration of probes improved the efficiency by an average of 4.2-fold (**SN_7B**) and 2.9-fold (**SN_7C**), respectively. In combination, these effects were not additive (**SN_7D**). We observed the highest efficiency with including a third round of capture enrihcment (5.8 fold on average) (**SN_7E**). However, this is time-consuming, not cost-effective (as commercial probes are expensive), and comes at the expense of a higher duplication rate. We therefore focused on further optimizing the biotin pre-clearing (**Supplemental Note 2**). In combination with a 1.5-2x higher probe concentration, this leads to equally good capture effciencies, resulting in 68% on average for the multi-way module (**SN_7H**) and 77.8% on average for the pair-wise module (**SN_7H**). The optimizations of the capture efficiency for mwMCC and iMCC extend to iTMCC and iMCCu.

**SN_7.**
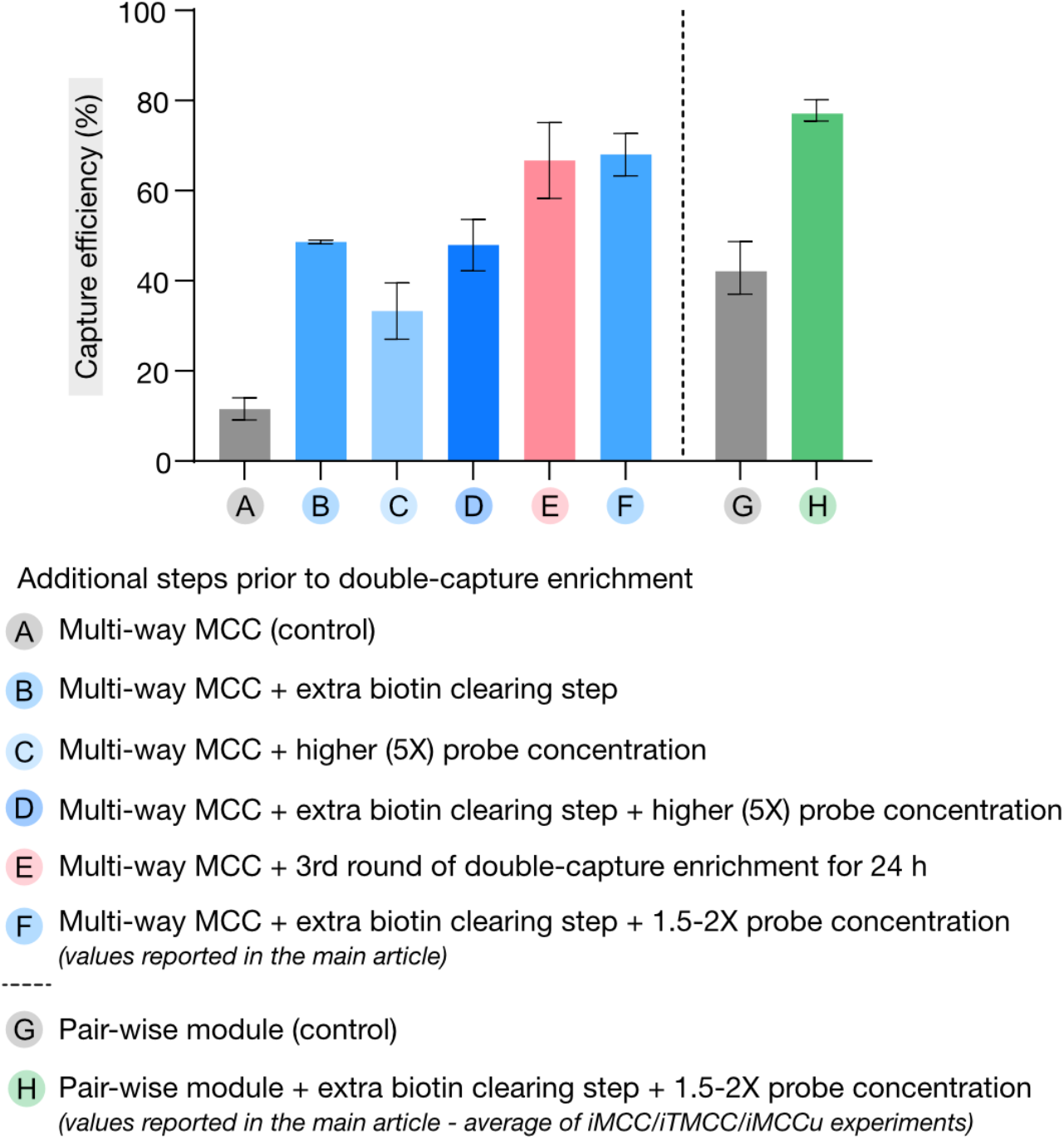

### Caspase-activated DNase

As we were preparing this manuscript, two pre-prints were published that report on the advantages of replacing MNase by caspase-activated DNase (CAD; also known as DNA fragmentation factor 40, DFF40)^81,82^. Compared to MNase, CAD allows for higher ligation efficiency, which is particularly useful for analysis of long “chromosome walks” that involve many ligation junctions. mwMCC predominantly relies on the detection of triplets. Extrapolating from the analyses presented in the pre-prints, integration of CAD could improve the efficiency with which triplets are detected in mwMCC by about 2-fold^81,82^. Replacing MNase with CAD is therefore likely a useful future mwMCC adaptation. However, since CAD is not commercially available, the usability of this adaptation would be limited to researchers with expertise in and equipment for protein purification.

## SUPPLEMENTARY NOTE 2

This supplementary note provides a step-by-step protocol for the improved pair-wise (iMCC/iTMCC/iMCCu) and multi-way (mwMCC) MNase-based 3C methods. This protocol builds on previously published protocols^14,16,76,77,83^ but is heavily optimized to allow for increased efficiency and throughput.

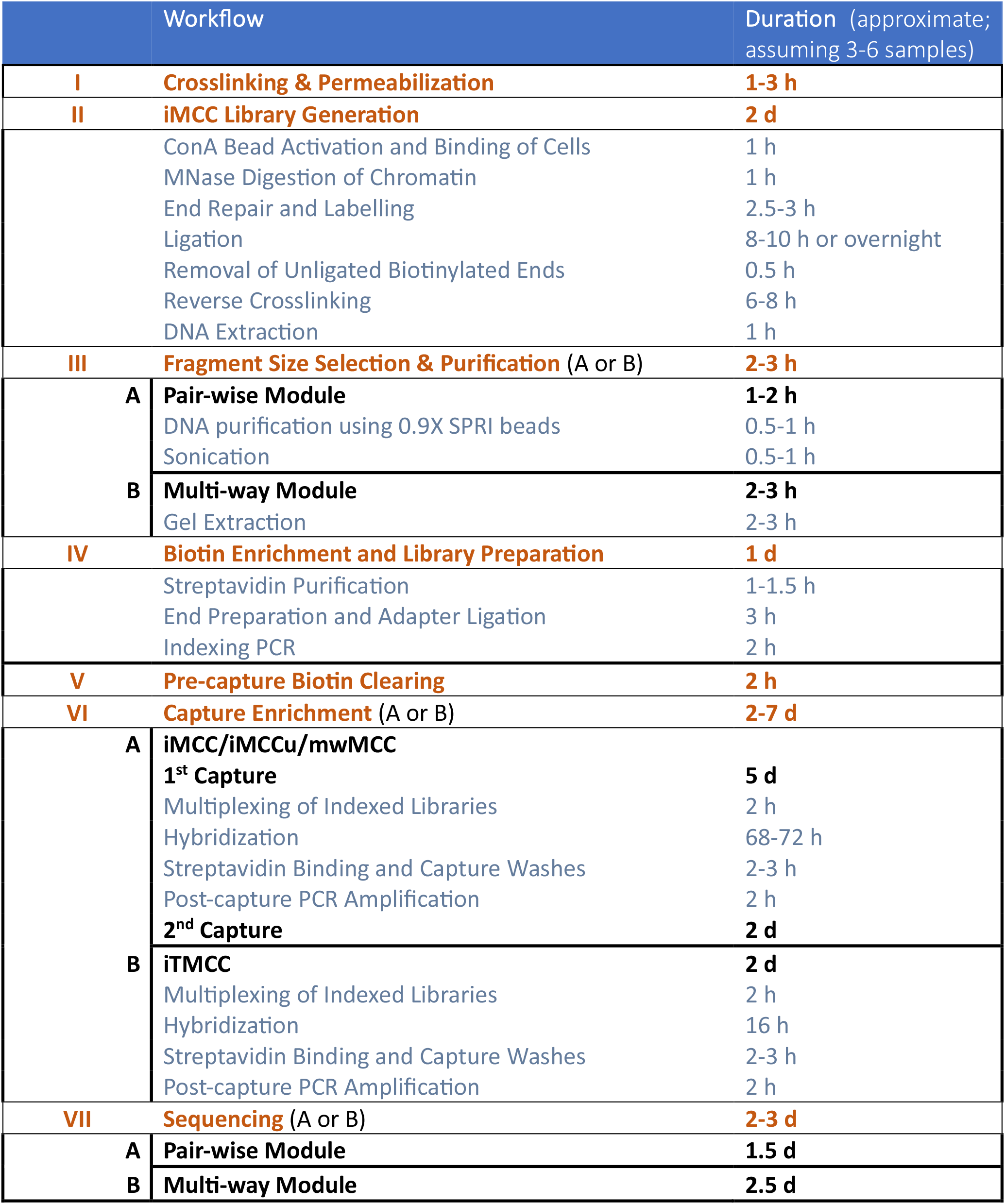

### Materials & Reagents

▪ 16% Formaldehyde (Thermo Scientific, 28908)
▪ Glycine (Sigma-Aldrich, G7126) Prepare 1.5 M stock solution, store at 4°C, and use within 1 week.
▪ Digitonin (Sigma-Aldrich, D141) Prepare 1% (w/v) digitonin by weighing 1-5 mg in an empty microcentrifuge tube and adding appropriate amount of DMSO (100-500 µL).
▪ CUTANA Concanavalin A-Conjugated Paramagnetic Beads (EpiCypher, 21-1411)
▪ HEPES pH-7.5 (Jena Bioscience, BU-106-75)
▪ 1 M KCl (Sigma-Aldrich, 1049361000) Prepare 1 M stock solution, filter-sterilize, and store at RT.
▪ 1 M MgCl_2_ (Invitrogen, AM9530G)
▪ 1 M MnCl_2_ (Fisher Scientific, 15405419)
▪ Micrococcal Nuclease (MNase) (20 gelUnits/µL) (NEB, M0247)
▪ 0.5 M CaCl_2_ (Sigma-Aldrich, C5080) Prepare 0.5 M stock solution, filter-sterilize, aliquot, and store at 4°C.
▪ 0.5 M EGTA pH-8 (Fisher Scientific, 15415795)
▪ 0.5 M EDTA pH-8 (Invitrogen, AM9260G)
▪ 10X NEBuffe r2.1 (NEB, B6002)
▪ 100 mM ATP (Thermo Scientific, R1441)
▪ 1 M Dithiothreitol (DTT) (Sigma-Aldrich, 646563)
▪ T4 PNK Kinase (NEB, M0201)
▪ DNA Polymerase I, Large (Klenow) Fragment (NEB, M0210)
▪ 1 mM Biotin-14-dATP (Jena Bioscience, BU-835-BIO14)
▪ 1 mM Biotin-11-dCTP (Jena Bioscience, BU-809-BIOX)
▪ 100 mM dGTP (Jena Bioscience, BU-1003)
▪ 100 mM dTTP (Jena Bioscience, BU-1004)
▪ 200X BSA (Sigma-Aldrich, B8667)
▪ T4 DNA Ligase Reaction Buffer (NEB, B0202)
▪ T4 DNA Ligase (NEB, M0202)
▪ NEBuffer 1 (NEB, B7001)
▪ Exonuclease III (E. coli) (NEB, M0206)
▪ Proteinase K (Thermo Scientific, EO0491)
▪ RNase A (Thermo Scientific, EN0531)
▪ DNeasy Blood & Tissue DNA Extraction Kit (Qiagen, 69506)
▪ microTUBE AFA Fiber Pre-Slit Snap-Cap (Covaris, 520045)
▪ 6X Gel Loading Dye (NEB, B7024)
▪ GeneJET Gel Extraction Kit (Thermo Scientific, K0692)
▪ 1 M Tris-HCl pH-7.5 (Invitrogen, 15567027)
▪ 1 M Tris-HCl pH-8 (Invitrogen, AM9855G)
▪ 5 M NaCl (Invitrogen, AM9760G)
▪ Tween 20 (Sigma-Aldrich, P1379)
▪ Dynabeads MyOne Streptavidin T1 (Invitrogen, 65601)
▪ Dynabeads M-270 Streptavidin (Invitrogen, 65305)
▪ NEBNext Ultra II DNA Library Prep Kit for Illumina (NEB, 7645)
▪ Herculase II Fusion Polymerase kit (Agilent, 600677)
▪ Mag-bind TotalPure NGS Beads (Omega Bio-Tek, M1378-01) (or alternatives)
▪ Biotinylated capture oligonucleotides (xGen MRD Hybridization Pane/Twist Custom Panel)
▪ KAPA HyperCapture Reagent kit (Roche, 9075828001)
▪ KAPA Library Amplification kit + Primer Premix (Roche, 07958978001)
▪ Twist Standard Hyb and Wash Kit (for iTiled-MCC) (Twist Bioscience, 116614)
▪ Twist Universal Blockers (for iTiled-MCC) (Twist Bioscience, 100578)
▪ Twist Binding and Purification Beads (for iTiled-MCC) (Twist Bioscience, 100983)
▪ Mouse Cot-1 DNA (for mouse-origin samples) (Invitrogen, 18440-016)
▪ Qubit 1X dsDNA High Sensitivity (HS) and Broad Range (BR) Assay Kits (Invitrogen, Q33231)
▪ D1000 ScreenTape and Reagents (Agilent, 50675582 & 50675583)
▪ Agilent High Sensitivity NGS Fragment Kit (Agilent, DNF-474-0500)

### Stock Solutions

#### 10X MNase reaction buffer

Prepare 10X buffer in bulk and aliquot into 1.5-mL microcentrifuge tubes. Store at -20°C.

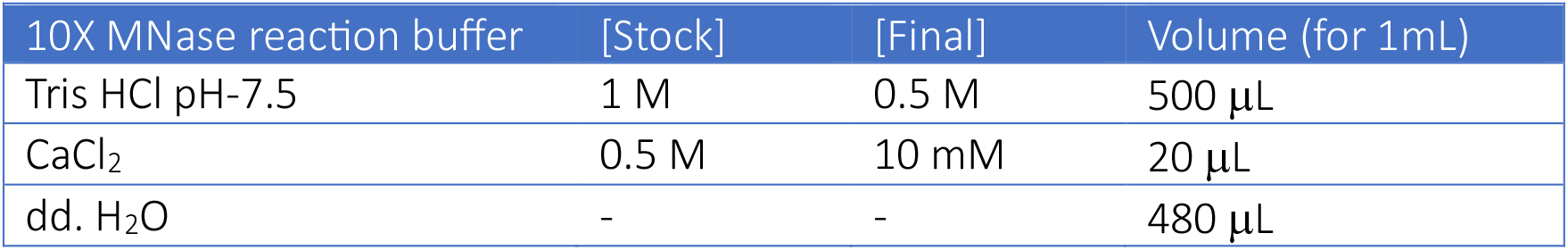

#### MNase storage buffer

Prepare 1:10 or 1:100 MNase dilution in MNase storage buffer and aliquot into small volumes. Store at -20°C or -80°C and avoid repeated freeze-thaw cycles.

| MNase storage buffer | [Stock] | [Final] | Volume (for 1mL) |
| --- | --- | --- | --- |
| Tris HCl pH-7.5 | 1 M | 10 mM | 10 µL |
| EDTA pH-8 | 0.5 M | 2 mM | 4 µL |
| NaCl | 5 M | 50 mM | 10 µL |
| Glycerol | 100% | 50% | 500 µL |
| dd. H<sub>2</sub>O | - | - | 476 µL |

#### 2X Binding and Washing buffer (referred to as BW buffer)

Prepare 2X BW buffer in bulk, filter-sterilize, and store at RT.

2X Binding and Washing buffer (referred to as BW buffer) Prepare 2X BW buffer in bulk, filter-sterilize, and store at RT.
| 2X BW buffer | [Stock] | [Final] | Volume (for 500 mL) |
| --- | --- | --- | --- |
| Tris HCl pH-7.5 | 1 M | 10 mM | 5 mL |
| EDTA pH-8 | 0.5 | 1 mM | 1 mL |
| NaCl | 5 M | 2 M | 200 mL |
| dd. H<sub>2</sub>O | - | - | 294 mL |

#### 1X BW buffer containing 0.1% [v/v] Tween-20 (referred to as TBW buffer)

Prepare 1X TBW buffer in bulk, stir well, filter-sterilize, and store at RT.

1X BW buffer containing 0.1% [v/v] Tween-20 (referred to as TBW buffer) Prepare 1X TBW buffer in bulk, stir well, filter-sterilize, and store at RT.
| 1X TBW buffer | [Stock] | [Final] | Volume (for 500 mL) |
| --- | --- | --- | --- |
| 2X BW | 2 M | 1 M | 250 mL |
| Tween-20 | 100% | 0.1% | 500 μL |
| dd. H<sub>2</sub>O | - | - | 250 mL |

### Protocol

#### I. Crosslinking & Permeabilization

*Note: If possible, fix 40-100 x 10*^*6*^ *cells per biological replicate to ensure enough material for MNase titration, main experiments, and backup aliquots*.

--- for each biological replicate ---

1. Trypsinize (if adherent) and centrifuge cells for 5 min at 300 x g and room temperature (RT). Resuspend the cell pellet in media, count cells, transfer 10 x 10^6^ cells per 15-mL Falcon tube and adjust the final volume to 8.75 mL with media. *If processing more than 10 x 10*^*6*^*cells, it may be convenient to scale up and use 50 mL Falcon tubes (up to 40 x 10*^*6*^ *cells in 35 mL of media per tube)*.
2. Add 1.25 mL of 16% formaldehyde (2%, final) to the cell suspension.
3. Incubate for 10 min at RT while gently tumbling or rotating.
4. Quench formaldehyde by adding 1.5 mL of 1.5 M glycine (∼0.15 M, final).
5. Centrifuge for 5 min at 300 x g and 4°C. Discard the supernatant.
6. Resuspend the cell pellet in 10 mL of ice-cold 1X PBS. *At this step, cell pellets can be pooled into a single 15 mL Falcon tube (if more than 1 tube)*.
7. Centrifuge for 5 min at 300 x g and 4°C. Discard the supernatant.
8. Resuspend the cell pellet in ice-cold 1X PBS to a concentration of 5 x 10^6^ cells/mL (based on the initial cell count) and aliquot 1 mL per tube into 1.5-mL microcentrifuge tubes. *Cell loss during washes is expected, so the final concentration may not be exactly 5 x 10*^6^ *cells/mL. This is fine as the goal is to ensure equal cell numbers across all aliquots to maintain consistency*.
9. Add 5 µL of 1% digitonin (0.005% [w/v], final) to each tube, mix by inversion, and incubate for 10 min at RT. *Use the optimized digitonin concentration for your specific cell type*.
10. Snap-freeze the aliquots using liquid nitrogen and store at -80°C.

#### II. iMCC Library Generation

--- for each aliquot ---

##### ConA Bead Activation and Binding of Cells

11. Thaw a fixed cell aliquot (on ice). Prepare fresh concanavalin A (conA) bead activation buffer and store at 4°C or on ice until ready to use. 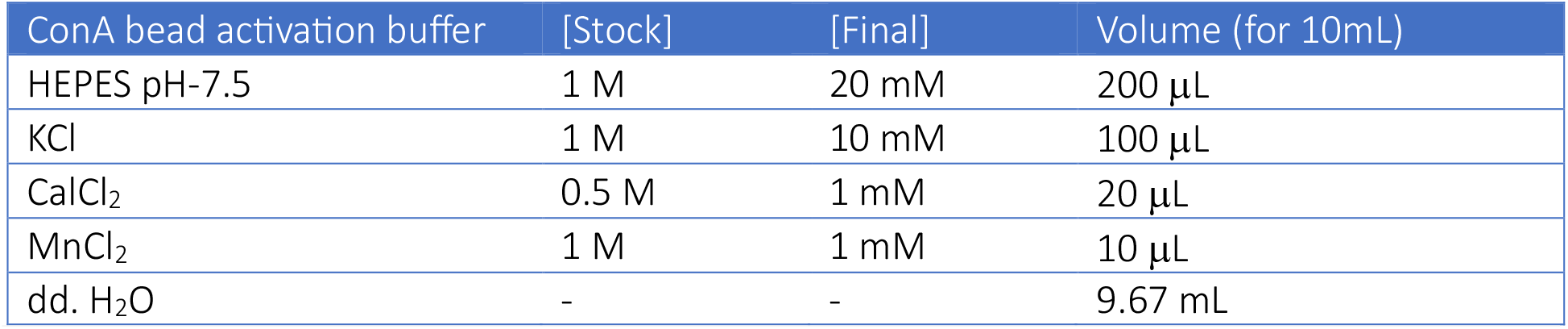 *Scale to the number of samples.*
12. Transfer 100 µL of conA beads to a 1.5-mL microcentrifuge tube. Spin down briefly, magnetize, and discard the supernatant.
13. Resuspend the beads in 1 mL of ice-cold bead activation buffer and mix well by pipetting. Spin down briefly, magnetize, and discard the supernatant.
14. Repeat Step 13 once for a total of two washes.
15. Resuspend the beads in 100 µL of ice-cold bead activation buffer and keep on ice until the aliquot is fully thawed. Then, add the activated beads to cells.
16. Incubate the mixture for ∼15-30 min at RT on a rotator (4-5 rpm). Spin down briefly, magnetize, and discard the supernatant.

##### MNase Digestion of Chromatin

17. Resuspend the beads with bound cells in ∼450 µL of dd. H_2_O and add 50 µL of 10X MNase reaction buffer. *Note that some extra volume may remain from the previous step. Adjust the dd*.*H*_2_*O volume accordingly to reach a final volume of 500 µL*.
18. Add MNase to the sample. Mix gently by inversion. *The required concentration should be determined by titration for each new MNase lot*.
19. Incubate for 1 h at 37°C and 550 rpm on a thermomixer. *As beads settle quickly, mix gently by inversion every ∼15 min*.
20. Quench MNase by adding 5 µL of 0.5M EGTA (5 mM, final) and mix gently by inversion. Spin down briefly, magnetize, and discard the supernatant.
21. Resuspend the beads in 1 mL of ice-cold 1X PBS and add 10 µL of 0.5 M EGTA. Mix gently by inversion and remove 50 µL as a digestion control. Spin down the remainder briefly, magnetize, and discard the supernatant.

###### Digestion QC *(perform the QC in parallel to the next steps)*

*DNA extraction using the DNeasy Blood and Tissue Kit (Qiagen, 69506)*.

1. Use the 50 µL digestion control aliquot from Step 21 and adjust to 200 µL with 1X PBS.
2. Add 20 µL of Proteinase K and 200 µL of buffer AL. Vortex thoroughly.
3. Incubate for 2 h on a thermomixer at 65°C and 1000 rpm (for reverse crosslinking).
4. Add 200 µL of 100% EtOH, vortex, and transfer the mixture to a spin column.
5. Centrifuge for 1 min at 6000 x g and RT. Discard the flowthrough.
6. Add 500 µL of Buffer AW1 and centrifuge for 1 min at 6000 x g and RT. Discard the flowthrough.
7. Add 500 µL of Buffer AW2 and centrifuge for 3 min at 20.000 x g and RT.
8. Place the spin column onto a 1.5-mL microcentrifuge tube and add 30 µL of Buffer AE. Incubate for 1 min at RT.
9. Centrifuge for 1 min at 6000 x g and RT. Discard the spin column. Samples can be stored at -20°C.
10. Run the sample on TapeStation or Fragment Analyzer to assess the digestion efficiency.

##### End Repair and Labelling

22. Resuspend the beads in 85 µL of end-repair buffer mix. 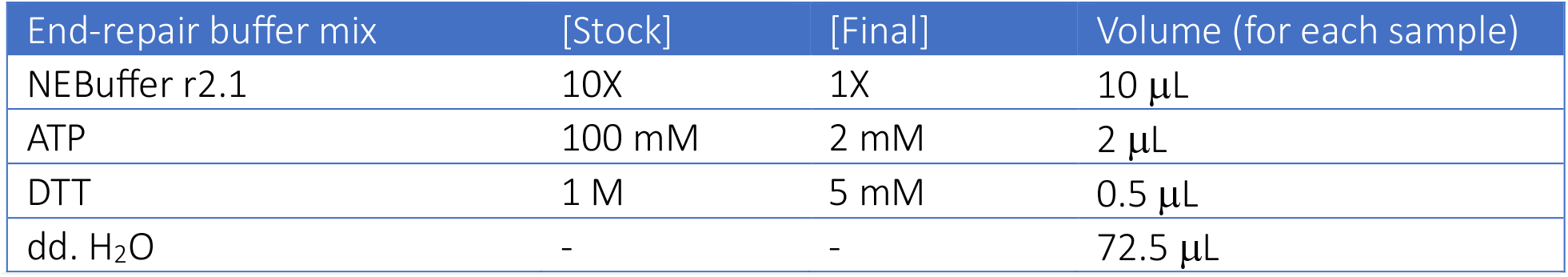 *Prepare a master mix scaled to the number of samples.*
23. Add 5 µL of T4 PNK (50 U, final), mix gently by pipetting, and incubate for 30 min at 37°C and 550 rpm on a thermomixer. *Gently flick the tube halfway through incubation*.
24. Add 10 µL of Klenow (50 U, final), mix gently by pipetting, and incubate for 30 min at 37°C and 550 rpm on a thermomixer. *Gently flick the tube halfway through incubation*.
25. Add 50 µL of end-label mix and mix gently by pipetting. 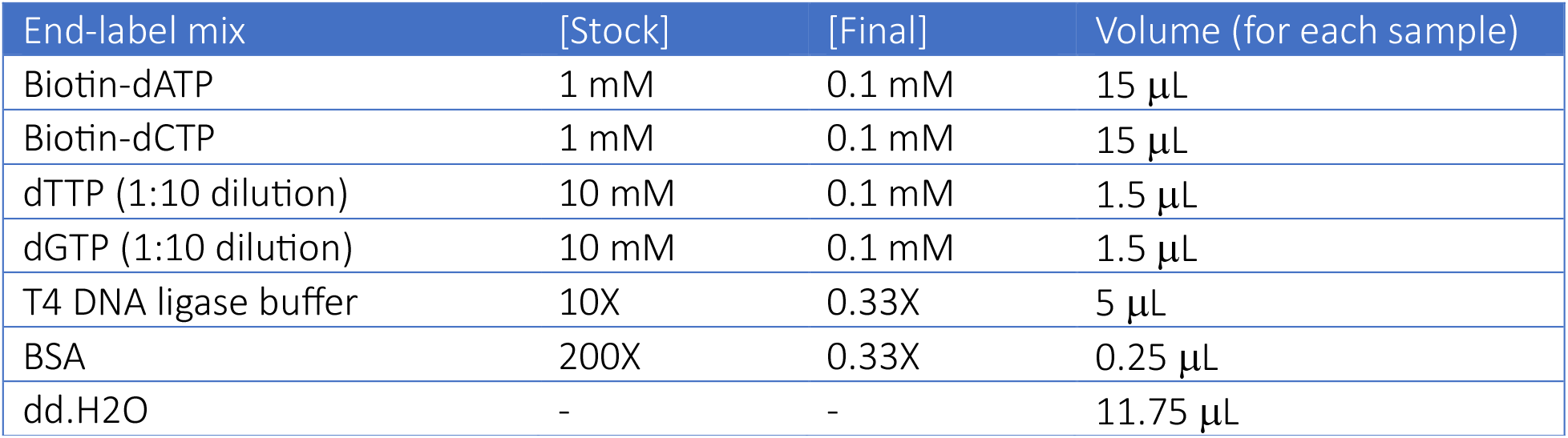 *Prepare a master mix scaled to the number of samples*.
26. Incubate for 1-1.5 h at 25°C and 550 rpm on a thermomixer. *Gently flick the tube halfway through incubation*.
27. Quench reaction by adding 9 µL of 0.5 M EDTA (30 mM, final). Spin down briefly, magnetize, and discard the supernatant.
28. Resuspend the beads in 1 mL of ice-cold 1X PBS. Spin down briefly, magnetize, and discard the supernatant.

##### Ligation

29. Resuspend the beads in 500 µL of ligation mix. Mix gently by pipetting. 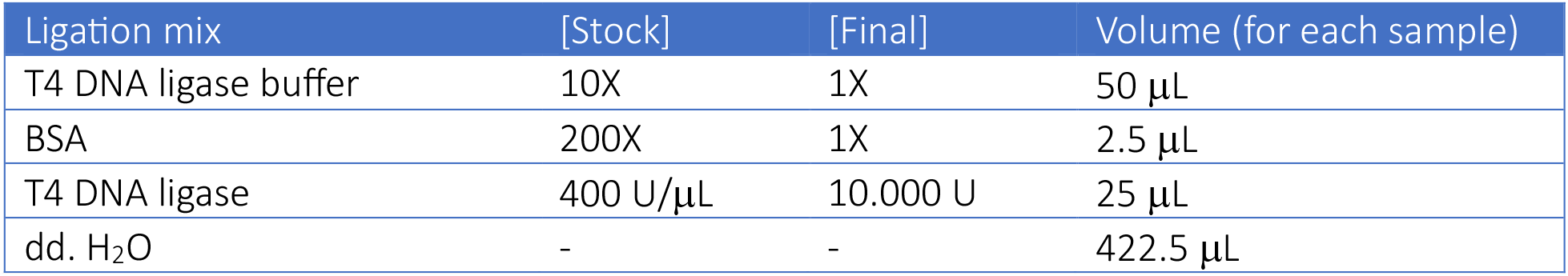 *Prepare a master mix scaled to the number of samples.* *Add T4 DNA ligase to the master mix just before use.*
31. Incubate for at least 8 h or overnight at 20°C and 550 rpm on a thermomixer. Spin down briefly, magnetize and discard the supernatant.

##### Removal of Unligated Biotinylated Ends

32. Resuspend the beads in of 170 µL of dd.H_2_O. Then, add 20 µL of 10X NEBuffer 1 and 10 µL of Exonuclease III (1000U). Mix gently by pipetting.
33. Incubate for 15 min at 37°C and 550 rpm on a thermomixer.

##### Reverse Crosslinking

34. Add 2 µL of 10 mg/mL RNase A (0.1 mg/mL, final), 20 µL of Proteinase K (2 mg/μL, final) and 200 µL of buffer AL. Mix by vortexing.
35. Incubate for 6-8 h (minimum 4 h) at 65°C and 1000 rpm on a thermomixer.

##### DNA Extraction

*DNA extraction is performed using the DNeasy Blood and Tissue Kit (Qiagen, 69506) with slight modifications*.

36. Add 200 µL of 100% EtOH, vortex thoroughly and transfer the mixture to a spin column.
37. Centrifuge for 1 min at 6000 x g and RT. Discard the flowthrough.
38. Add 500 of µL buffer AW1 and centrifuge for 1 min at 6000 x g and RT. Discard the flowthrough.
39. Add 500 µL of buffer AW2 and centrifuge for 1 min at 6000 x g and RT. Discard the flowthrough.
40. Centrifuge for 1 min at 20.000 x g and RT (dry spin).
41. Place the spin column onto a 1.5-mL microcentrifuge tube and add 75 µL of buffer AE (pre-heated at 65°C). Incubate for 1-2 min at RT. Then, centrifuge for 2 min at 10.000 x g and RT.
42. Repeat the elution by adding another 75 µL of buffer AE (pre-heated at 65°C). Incubate for 1-2 min at RT and centrifuge for 2 min at 10.000 x g and RT. Discard the spin column.
43. Run the sample on TapeStation or Fragment Analyzer to assess the ligation efficiency and measure the DNA concentration using Qubit BR Kit.

##### SAFE STOPPING POINT

*Store the sample at -20°C for long-term storage or at 4°C if to be used within a few days*.

### III Fragment Size Selection & Purification

*For pair-wise module (iMCC/iTMCC/iMCCu), follow Steps 44-54 and skip Steps 55-70. For multi-way module (mwMCC), skip Steps 44-54 and follow Steps 55-70*.

#### A. Pair-wise Module

##### DNA purification using 0.9X SPRI beads

*This is done to remove mononucleosome fragments prior to sonication. Although biotin-mediated enrichment of chimeric fragments in this protocol in theory makes this step dispensable, we observe a slightly improved chimeric ratio when it is performed and therefore recommend following it*.

44. Bring 135 µL (0.9X) of MagBind TotalPure NGS beads to RT. Add beads to the sample. Mix well by pipetting.
45. Incubate for 10 min at RT. Magnetize and discard the supernatant once clear.
46. Add 800 µL of freshly prepared 80% ethanol while still magnetized. Wait 30 sec and discard ethanol.
47. Repeat Step 46 for a total of two washes. Spin down briefly and discard any remaining traces of ethanol.
48. Air dry beads on magnetic stand at RT until matte in appearance.
49. Resuspend the beads in 135 µL TE buffer. Incubate for 5 min at RT. *Resuspending in TE buffer rather than water is important for optimal sonication efficiency*.
50. Magnetize and once clear, recover 130 µL of eluate and directly transfer to a Covaris microTUBE.

##### Sonication

51. Sonicate the sample using a Covaris S220 Focused-ultrasonicator to achieve an average target fragment size of 180-200 bp. 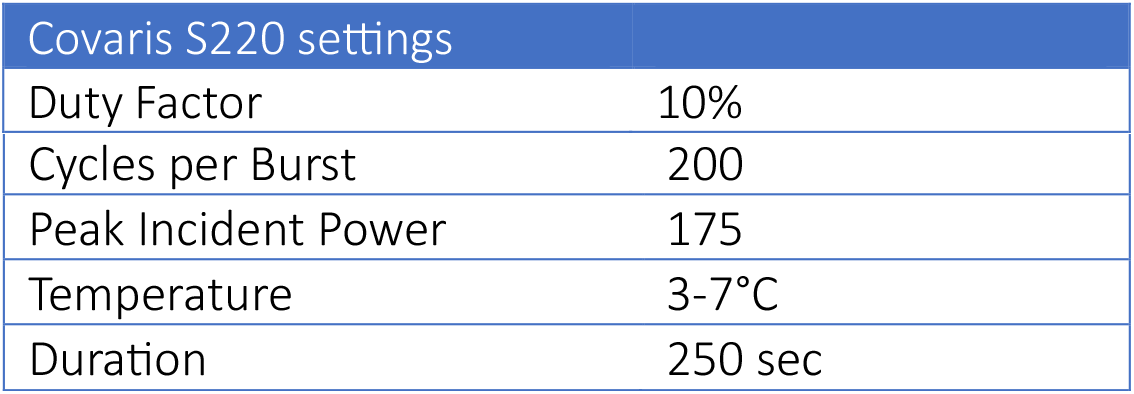 *For new cell types, optimize the sonication duration (try 200-300 sec).*
52. Remove 1 µL and run the sample on TapeStation. If the sample looks under-sonicated, sonicate the remainder for additional time until the desired fragment size is achieved. *Due to broad fragment size distribution, TapeStation tends to overestimate the average fragment size. Fragment Analyzer provides a more accurate size estimate. We usually verify sonication efficiency using Fragment Analyzer during optimization to avoid over-sonication based on TapeStation profiles alone. See ‘NOTES’ section for an example*.
53. After sonication, transfer the entire volume (∼130 µL) to a fresh 1.5-mL microcentrifuge tube and perform DNA purification as described in Steps 44-50 using 1.8X (234 µL) NGS beads.
54. In the final step, elute into 150 µL of H_2_O and recover the eluate. Measure the DNA concentration using Qubit BR Kit.

#### B. Multi-way Module

##### Gel Extraction

*Gel extraction is performed using the GeneJet Gel Extraction Kit (Thermo Scientific, K0692) with some modifications*.

55. Prepare a 170 mL of 1% agarose gel by combining 5 adjacent wells of an 18-well gel comb per sample.
56. Add 15-20 6X Gel Loading Dye to the sample (from Step 42), mix well and load into the combined well. *Adapt Steps 55 and 56 based on the available gel electrophoresis setup to be able to accommodate larger input volumes (∼170 µL)*.
57. Run the sample for 45-50 min at 120 V. *Monitor the run and adjust run time accordingly. Since the desired fragment range is 500-800 bp, the ideal duration should allow sufficient resolution for efficient gel excision while minimizing the gel volume to maximize the yield*.
58. Cut the gel slice containing 500-800 bp fragments into multiple parts using a clean scalpel on a transilluminator.
59. Transfer the gel slice parts into multiple pre-weighed 2-mL microcentrifuge tubes with a maximum of 1 g per tube and record the weight of each tube.
60. Add Binding Buffer to the gel slice in each tube at a 1:1 [volume:weight] ratio.
61. Incubate the gel mixture at 60°C for 20-30 min or until completely dissolved, then vortex thoroughly. *Mix by inversion every 5-10 min during incubation*.
62. Transfer the solubilized gel solution (up to 800 µL) to a spin column. *If total volume > 800 µL, the solution can be added in stages*.
63. Centrifuge for 1 min at 14.000 x g and RT. Discard the flowthrough.
64. Repeat Steps 62 and 63 until the entire volume has been transferred.
65. Add 100 µL of Binding buffer and centrifuge for 1 min at 14.000 x g and RT. Discard the flowthrough.
66. Add 700 µL of Wash Buffer and centrifuge for 1 min at 14.000 x g and RT. Discard the flowthrough.
67. Centrifuge for 1 min at 14.000 x g and RT (dry spin).
68. Place the spin column onto a 1.5-mL microcentrifuge tube and add 75 µL of Elution Buffer (pre-heated at 65°C). Incubate for 1-2 min at RT and centrifuge for 2 min at 10.000 x g and RT.
69. Repeat elution by adding another 75 µL of Elution Buffer (pre-heated at 65°C). Incubate for 1-2 min at RT and centrifuge for 2 min at 10.000 x g and RT. Discard the spin column.
70. Run the sample on TapeStation or Fragment Analyzer to assess the fragment distribution and measure the DNA concentration using Qubit BR Kit.

##### SAFE STOPPING POINT

*Store the sample at -20°C for long-term storage or at 4°C if to be used within a few days*.

## IV. Biotin Enrichment and Library Preparation

### Streptavidin Purification

71. Transfer 25 µL of Dynabeads MyOne Streptavidin T1 (pre-equilibrated to RT) to a 1.5-mL microcentrifuge tube and add 1 mL of 1X TBW. Mix by inversion, magnetize and discard the supernatant.
72. Repeat the wash with 1X TBW twice for a total of 3 washes.
73. Resuspend the beads in 150 µL of 2X BW and add the sample from Step 54 (pair-wise module) or Step 70 (multi-way module).
74. Incubate for 30 min (up to overnight) at RT on a nutator/rotator at gentle speed.
75. Spin down briefly, magnetize, and discard the supernatant.
76. Resuspend the beads in 1 mL of 1X TBW. Mix by inversion and incubate for 5 min at RT and 600 rpm on a thermomixer. Spin down briefly, magnetize, and discard the supernatant.
77. Repeat Step 76.
78. Wash the beads with 500 µL of 10 mM Tris (pH-8) while still magnetized. Discard the supernatant.
79. Resuspend the beads in 100 µL of ddH_2_O and split into two 1.5-mL microcentrifuge tubes as independent reactions.

### End Preparation and Adapter Ligation

*Library preparation is performed using the NEBNext Ultra II DNA Library Prep Kit for Illumina Kit (NEB, 7645) with some modifications*.

--- for each reaction ---

80. Add 7 µL of End Prep Reaction Buffer and 3 µL of End Prep Enzyme Mix. Mix by pipetting.
81. Incubate for 30 min at 20°C and 600 rpm on a thermomixer.
82. Inactivate the reaction by incubating at 65°C for 30 min without shaking. Then, immediately transfer to ice.
83. Add 2.5 µL of NEB Illumina Adapter, 30 µL of Ultra II Ligation Master Mix and 1 µL of Ligation Enhancer. Mix by pipetting. *If preparing a master mix, add the adapter separately to each sample to avoid adapter dimer formation*.
84. Incubate for 30 min at 20°C and 600 rpm on a thermomixer.
85. Add 3 µL of USER enzyme. Mix by pipetting.
86. Incubate for 15 min at 37°C and 600 rpm on a thermomixer.
87. Add 1 mL of 1X TBW. Mix by inversion and incubate for 2 min at RT and 600 rpm on a thermomixer. Spin down briefly, magnetize, and discard the supernatant.
88. Repeat Step 87.
89. Wash the beads with 500 µL of 10 mM Tris (pH-8) while still magnetized. Discard the supernatant.
90. Resuspend the beads in 57 µL of dd. H_2_O and split into two PCR tubes (28.5 µL each) as independent reactions. *At this point, each sample will have been split as 4 reactions*.

## SAFE STOPPING POINT

*Store the sample at -20°C or 4°C for a few days though it is recommended to proceed immediately with indexing PCR*.

### Indexing PCR

91. Add 16.5 μL of PCR mix to the reaction and mix well by pipetting. 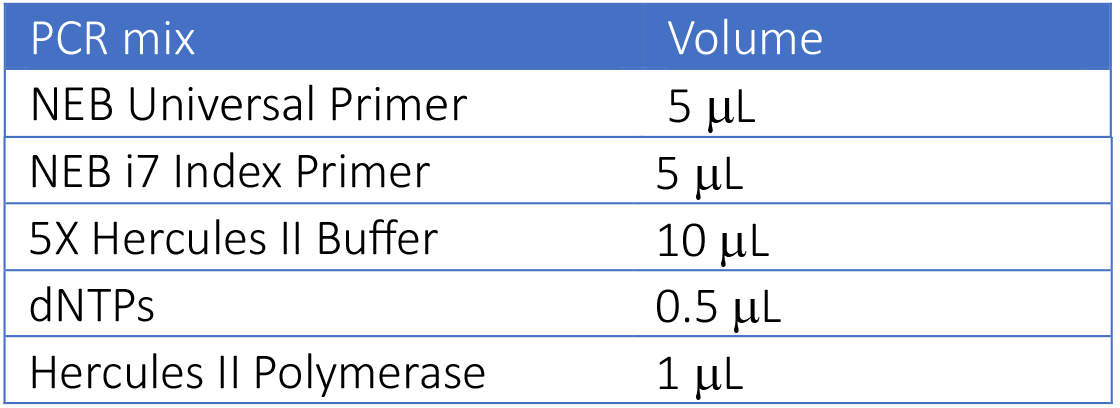 *Prepare a master mix (excluding the NEBNext i7 index primer) scaled to the number of reactions/samples. Add Hercules II Polymerase to the master mix just before use*. *Use the same i7 index primer for all split reactions from the same sample, or different i7 indices if the split reactions are intended to be used as technical replicates across multiple capture panels simultaneously*. *Since NEB has discontinued the sales of NEBNext Multiplex Oligos for Illumina (E7335; E7500; E7710; E7730), this step can alternatively be adapted to using NEBNext Multiplex Oligos for Illumina (96 Unique Dual Index Primer Pairs) (NEB, E6440)*.
92. Perform PCR for a total of 10 cycles using following settings: 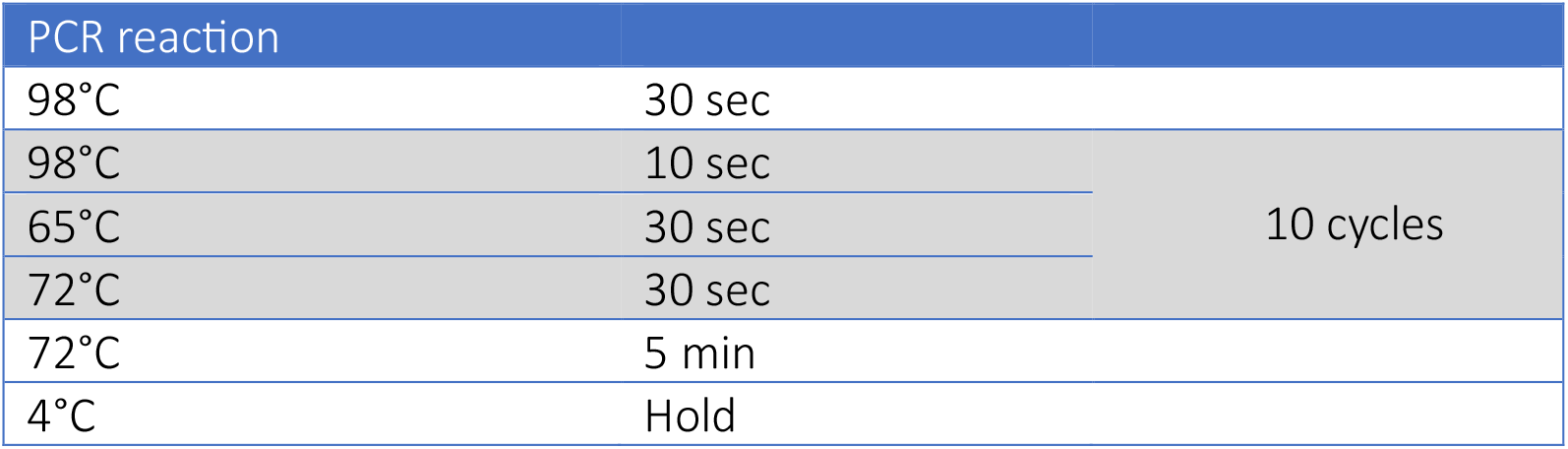
93. At this point, pool all the split reactions from the same sample and perform DNA purification as described in Steps 44-50 using 0.9X NGS beads.
94. In the final step, elute into 200 µL of dd. H_2_O and assess the sample on TapeStation or Fragment Analyzer and measure the DNA concentration using Qubit BR Kit.

## SAFE STOPPING POINT

*Store the sample at -20°C for long-term storage or at 4°C if to be used within a few days*.

### V. Pre-capture Biotin Clearing

*This is done to improve the on-target capture efficiency during the targeted capture enrichment in the next section*.

95. Transfer 25 µL of Dynabeads MyOne Streptavidin T1 (pre-equilibrated to RT) to a fresh 1.5-mL microcentrifuge tube and add 500 µL 1X TBW. Spin down briefly, magnetize, and discard the supernatant.
96. Repeat the wash with 1X TBW twice for a total of 3 washes.
97. Resuspend the beads in 200 µL of 2X BW and add the sample from Step 94.
98. Incubate at RT for 30 min on a nutator/rotator at gentle speed.
99. Magnetize, carefully collect the supernatant and transfer to a fresh 1.5-mL microcentrifuge tube. Discard the beads. *Leave a few µL of supernatant behind to avoid pipetting beads, which may result in contamination*.
100. Magnetize and transfer the supernatant again to a fresh 1.5-mL microcentrifuge tube.
101. Perform DNA purification as described in Steps 44-50 using 0.9X NGS beads.
102. In the final step, elute into 100 µL of dd. H_2_O and transfer the cleared eluate to a fresh 1.5-mL microcentrifuge tube.
103. Run the sample on TapeStation or Fragment Analyzer to assess the library quality and measure the DNA concentration using Qubit BR Kit.

## SAFE STOPPING POINT

*Store the sample at -20°C for long-term storage or at 4°C if to be used within a few days*.

### VI. Capture Enrichment

#### A. iMCC/iMCCu/mwMCC

*For iMCC/iMCCu/mwMCC, skip Steps 132-168*.

*Capture Enrichment is performed using KAPA HyperCapture Reagent kit (Roche, 9075828001) and is based on Downes et al. (Nature Protocols, 2022) with a few modifications*.

**1**^**st**^ **Capture**

##### Multiplexing of Indexed Libraries

104. Pre-heat a vacuum concentrator to 45°C.
105. In a hybridization tube, pool 2 µg from each of 6 uniquely indexed libraries. *Alternatively, pool 1 µg from each of 12 uniquely indexed libraries. If the total volume exceeds 12 µg, split equally into up to 12 hybridization tubes, with a maximum of 12 µg per tube. For all further washes volumes are given “per library” i*.*e. per 2 µg of DNA*.
106. Add 30 µL of COT Human DNA (1 mg/mL stock; 5 µL per library). *Replace COT Human DNA with species-specific COT DNA depending on the input DNA source. If a species-specific COT DNA is not available, it can be replaced with KAPA Hybrid Enhancer Reagent (Roche, 09075763001)*.
107. Dry the library pool using the vacuum concentrator with tube lids open.

##### Hybridization

108. Add 40.2 µL of Universal Enhancing Oligonucleotides (6.7 µL per library), 84 µL of Hybridization Buffer (14 µL per library) and 36 µL of Hybridization Component H (6 µL per library). Mix carefully by pipetting. Spin down briefly and incubate at RT for 2 min.
109. Add 27 µL of diluted capture oligonucleotides (4.5 µL per library; 1-5.8 nM final individual oligonucleotide concentration) to the hybridization mixture and mix carefully by pipetting. Briefly spin down the mixture. *The single-stranded DNA oligonucleotides (120 bp) for a specific panel (iMCC/mwMCC) or a tiled region (iMCCu) are ordered as a pool and diluted to the desired final individual oligo concentration. The oligonucleotides are also effective at lower concentrations (e*.*g*., *2*.*9 nM per individual oligonucleotide). However, using higher concentrations helps with improving capture efficiencies in our protocol (See Supplementary Note 1)*.
110. Perform hybridization for 68-72 h in a thermocycler using the following settings: 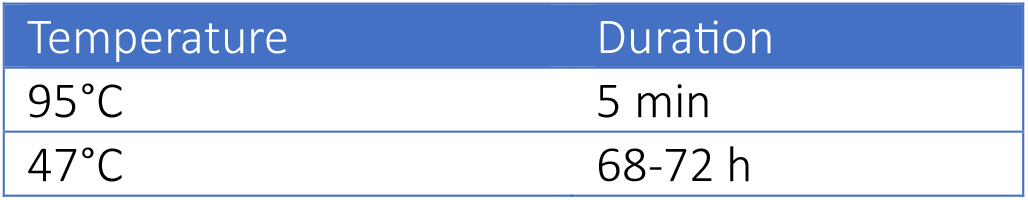

##### Streptavidin Binding and Capture Washes

Set a thermomixer to 47°C. Prepare the required wash buffers.

Streptavidin Binding and Capture Washes Set a thermomixer to 47°C. Prepare the required wash buffers.
| Buffer | Buffer [final] | Buffer volume | H<sub>2</sub>O volume |
| --- | --- | --- | --- |
| 2.5X Bead Wash Buffer | 1X | 400 µL | 600 µL |
| 10X Stringent Wash Buffer | 1X | 150 µL | 1,35 mL |
| 10X Wash Buffer I | 1X | 93 µL | 837 µL |
| 10X Wash Buffer II | 1X | 60 µL | 540 µL |
| 10X Wash Buffer III | 1X | 60 µL | 540 µL |

*Place the stringent Wash Buffer and 330 µL of Wash Buffer I at 47°C, and the rest at RT*.

111. Transfer 300 µL of Dynabeads M-270 Streptavidin (pre-equilibrated to RT; 50 µL per library) to a fresh 1.5-mL low-binding microcentrifuge tube.
112. Add 300 µL of 1X Bead Wash Buffer (50 µL per library) and mix by vortexing. Spin down briefly, magnetize, and discard the supernatant.
113. Repeat Step 112 for a total of two washes.
114. Resuspend the beads in 300 µL of 1X Bead Wash Buffer (50 µL per library). Magnetize and discard the supernatant.
115. Once the hybridization (Step 110) is complete, open the thermocycler lid, briefly spin down the tube, and put them back in the thermocycler. While keeping the tube in the thermocycler, transfer the entire content to the beads from the previous step. *Work quickly during transfer from thermocycler at 47°C as it is critical to minimize off-target binding*.
116. Incubate for 45 min at 47°C and 600 rpm on a thermomixer. *Gently flick the tube every 10 minutes during incubation to avoid settling of the beads*.
117. Add 300 µL of pre-heated 1X Wash Buffer I (50 µL per library) to the beads and bound DNA. Vortex gently for 10 seconds. Briefly spin down, magnetize and discard the supernatant.
118. Immediately transfer the tube to the thermomixer. Resuspend the beads in 600 µL of pre-heated 1X Stringent Wash Buffer (100 µL per library) and mix gently by pipetting while keeping the tube on the thermomixer.
119. Incubate for 5 min at 47°C and 600 rpm on a thermomixer. Spin down briefly, magnetize, and discard the supernatant.
120. Repeat Steps 118 and 119 once with 1X Stringent Wash buffer for a total of two washes. *Throughout Steps 115-120, minimize time at RT and keep samples at 47°C as much as possible to reduce off-target binding*.
121. Resuspend the beads in 600 µL of RT 1X Wash Buffer I (100 µL per library). Vortex for 10 seconds and incubate at RT for 1 minute. Spin down briefly, magnetize, and discard the supernatant.
122. Resuspend the beads in 600 µL of RT 1X Wash Buffer II (100 µL per library). Vortex for 10 seconds and incubate at RT for 1 min. Spin down briefly, magnetize, and discard the supernatant.
123. Resuspend the beads in 600 µL of RT 1X Wash Buffer III (100 µL per library). Vortex for 10 seconds and incubate at RT for 1 min. Spin down briefly, magnetize, and discard the supernatant.
124. Resuspend the beads in 240 µL of dd. H_2_O (40 µL per library) and split into 12 PCR tubes (20 µL per tube).

##### Post-capture PCR Amplification

125. Add 30 µL of PCR mix to the reaction and mix well by pipetting. 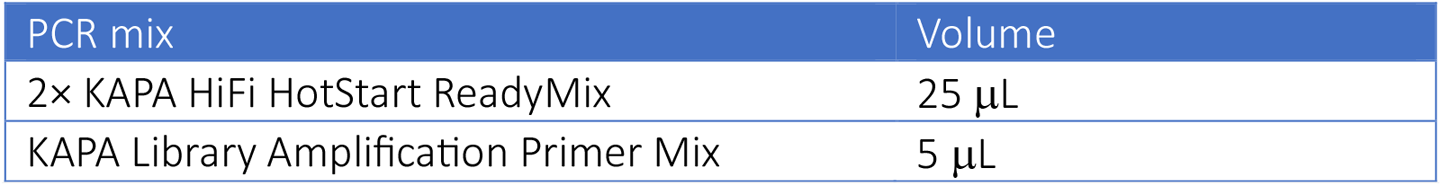 *Alternatively, at Step 124, resuspend the beads in 240 µL of dd. H*_2_*O and split equally into 2 tubes. Add 180 µL of PCR mix to each tube, then split into 6 PCR tubes (50 µL per tube, 12 tubes total)*.
126. Perform PCR for a total of 8 cycles using the following settings: 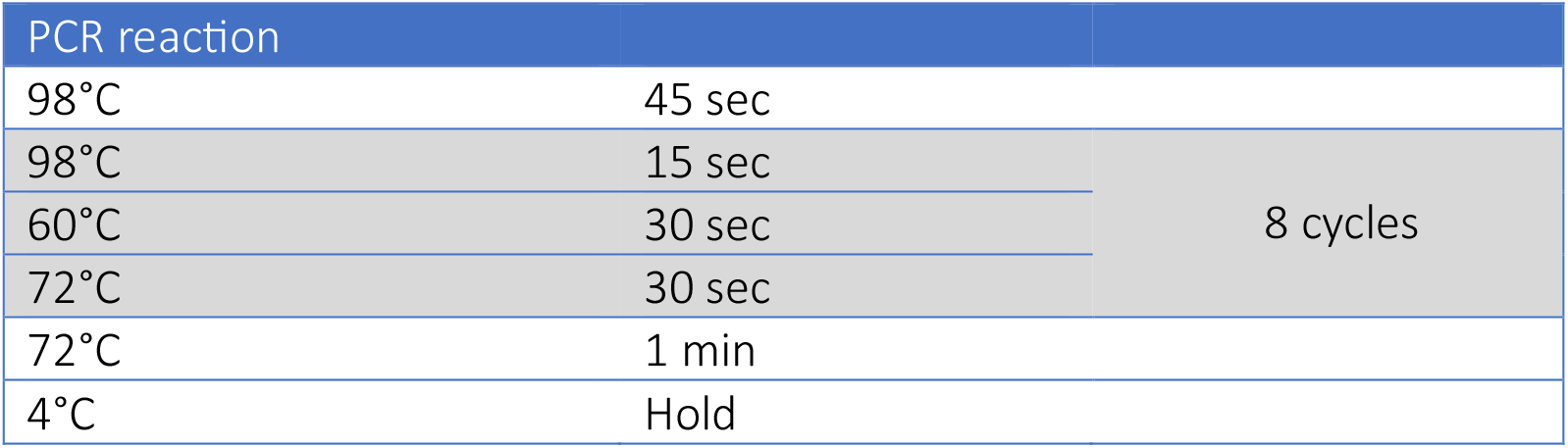
127. Pool 2-4 PCR reactions (100-200 µL) into a 1.5-mL microcentrifuge tube and perform DNA purification as described in Steps 44-50 using 1.8X NGS beads.
128. In the final step, elute into 40-80 µL of dd. H_2_O and transfer the cleared eluate to a fresh 1.5-mL microcentrifuge tube. *If the beads are difficult to resuspend due to large volumes used when processing more libraries, increase the elution volume accordingly*.
129. Run the sample on TapeStation or Fragment Analyzer to assess the final library quality and measure the DNA concentration using Qubit HS Kit.

## SAFE STOPPING POINT

*Store the sample at 4°C and proceed with the 2nd capture the following day*.

**2**^**nd**^ **Capture**

130. Perform hybridization for 24 h by following Steps 104-110. *If the yield after the 1*^*st*^ *capture is less than 2 µg, maintain single library volumes, or adjust the volumes according to the total input concentration (2 µg per library)*.
131. Perform Streptavidin binding, capture washes and PCR amplification (7 cycles) by following Steps 111-129 and adjust the volumes based on the previous step. In the final step, elute into 50 µL of dd. H_2_O and transfer the cleared eluate to a fresh 1.5-mL microcentrifuge tube.
132. Run the sample on TapeStation or Fragment Analyzer to assess the library quality and measure the DNA concentration using Qubit HS Kit.

## SAFE STOPPING POINT

*Store the sample at -20°C for long-term storage*.

*Optional: Repeat the targeted capture enrichment using the same indexed samples to increase library complexity. Final libraries from independent enrichments can be pooled prior to sequencing or sequenced independently and the FASTQ files merged afterwards*.

### B. iTMCC

*For iTiled-MCC, skip Steps 104-132*.

*Capture Enrichment is performed using the Twist Standard Hyb and Wash Kit (Twist Bioscience, 116614), Twist Universal Blockers (Twist Bioscience, 100578), and Twist Binding and Purification Beads (Twist Bioscience, 100983) as described in the manual and Aljahani et al. (Nature Communications, 2022) with slight modifications*.

#### Multiplexing of lndexed Libraries

133. Pre-heat a vacuum concentrator to 45°C. Pre-set a thermocycler to 95°C (with lid temperature set to 105°C). Pre-set a thermomixer to 65°C. On ice, thaw Twist Custom Probe Panel, Twist Hybridization Mix, Twist Hybridization Enhancer, Twist, Universal Blockers, and a species-specific COT DNA.
134. In a microcentrifuge tube, pool equal amounts of uniquely indexed libraries for each condition targeting 2 µg of total DNA per condition. *Example: For 6 replicates each of a control and a depletion condition, with a target of 2 µg total DNA per condition, pool 333*.*3 ng from each of the 6 control libraries, and 333*.*3 ng from each of the 6 depletion libraries*. *If the amount of library is not enough, use a smaller amount, however, this may result in decreased library complexity*.
135. Split into required number of hybridization tubes with a maximum of 4 µg of library pool per hybridization tube (2 µg of library pool per reaction). Briefly spin down the tube(s). *The volumes of reagents and buffers mentioned in the manual are originally intended for up to 4 µg of library pool per reaction. However, in our protocol, we use no more than 2 µg of library pool per reaction, with a maximum of 2 reactions (4 µg total) per hybridization tube*. *If using 2 reactions per hybridization tube, then double the volume of all reagents and buffers accordingly*.
136. Dry the library pool using the vacuum concentrator with tube lids open.

#### Hybridization

For each reaction per tube

137. Once dried, add 5 µL of Blocker Solution and 7 µL of Universal Blockers and keep on ice until needed.
138. Heat the Twist Hybridization Mix at 65°C until all precipitate is dissolved, then cool down to RT for 5 min.
139. Prepare the probe solution in a 0.2-mL microcentrifuge tube. 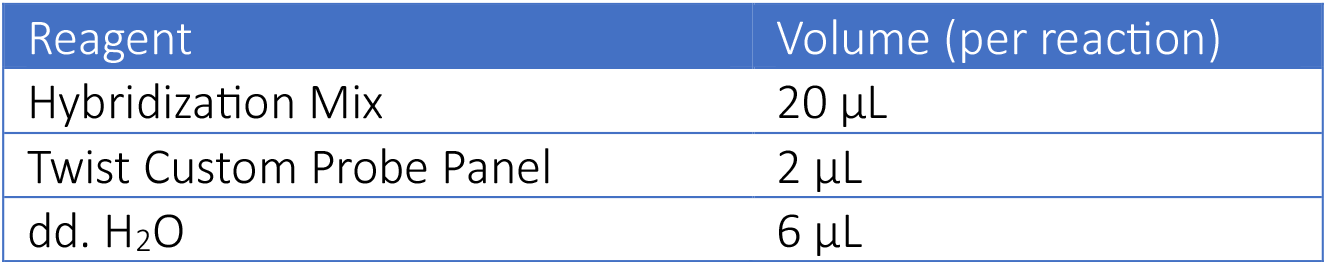 *Hybridization mix is viscous, so pipette slowly for accurate volume transfer*.
140. Incubate the probe solution at 95°C for 2 min in the thermocycler, then immediately transfer to ice or 4°C for 5 min. Next, leave the probe solution at RT for 5 min.
141. During the previous step, while the probe solution is on ice for 5 min, after 3 min passes, simultaneously start incubating the tube containing resuspended library pool (from Step 137) at 95°C for 5 min in the thermocycler, then leave at RT for 2 min, so that both solutions complete the incubation at approximately the same time.
142. Vortex and briefly spin down the probe solution, then transfer to the resuspended library pool.
143. Spin down briefly and add 30 µL of Hybridization Enhancer to the top of the reaction. *Do not vortex or mix by pipetting. Spin down briefly to ensure no bubbles*.
144. Incubate the hybridization mixture for 16 h at 70°C on a thermocycler (with lid temperature set to 85°C).

#### Streptavidin Binding and Capture Washes

For each reaction per tube

145. Pre-heat Binding Buffer, Wash Buffer 1, and Wash Buffer 2 to 48°C until all precipitate is dissolved. Then equilibrate Binding Buffer and Wash Buffer 1 to RT and leave Wash Buffer 2 at 48°C
146. Vortex the Streptavidin Binding Beads (pre-equilibrated to RT) until fully mixed. Transfer 100 µL to a fresh 1.5-mL microcentrifuge tube.
147. Add 200 µL of Binding Buffer to the beads and mix by pipetting. Magnetize and discard the supernatant.
148. Repeat Step 147 twice for a total of three washes.
149. Resuspend the beads in 200 µL of Binding Buffer.
150. Once the hybridization (Step 144) is complete, open the thermocycler lid, briefly spin down the tube, and put them back in the thermocycler. While keeping the tube in the thermocycler, transfer the entire content to the beads from the previous step. *Work quickly during transfer from thermocycler at 70°C as it is critical to minimize off-target binding*.
151. Mix well and incubate for 30 min at RT and 600 rpm on a thermomixer.
152. Spin down briefly, magnetize, and discard the supernatant including any remaining Hybridization Enhancer. *If some Hybridization Enhancer still remain visible after removing the supernatant, it will not affect the final capture product*.
153. Resuspend the beads in 200 µL of Wash Buffer 1. Mix by pipetting.
154. Briefly spin down the tube and transfer everything to a fresh 1.5-mL microcentrifuge tube. Magnetize and discard supernatant. *This is done to reduce the background from non-specific binding to the tube surface*.
155. Resuspend the beads in 200 µL of Wash Buffer 2. Mix by pipetting.
156. Incubate for 5 min at 48°C on a thermomixer. Spin down briefly, magnetize, and discard the supernatant.
157. Repeat Steps 155 and 156 twice for a total of three washes.
158. After the final wash, spin down briefly, magnetize, and remove final traces of supernatant.
159. Resuspend the beads in 45 µL of dd. H_2_O. Mix by pipetting and leave on ice until ready to proceed.
160. When ready, mix the beads by pipetting, if settled. Split into two 0.2-mL microcentrifuge tubes (∼22.5 µL each). *Alternatively, use one half and store the other at -20°C for future use*.

#### Post-capture PCR Amplification

161. Add 27.5 µL of PCR mix to each reaction and mix well by pipetting. 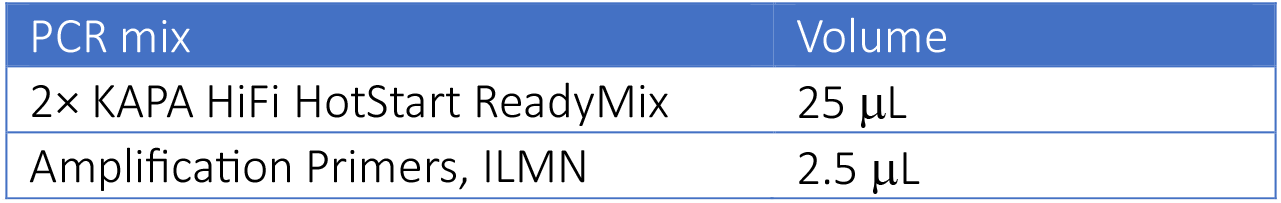 *TWIST Kit is supplied with Amplification Primers, ILMN. Alternatively, 5 μL KAPA Library Amplification Primer Mix (Roche, 07958978001) can be used*.
162. Perform PCR for a total of ‘N’ cycles using following settings: 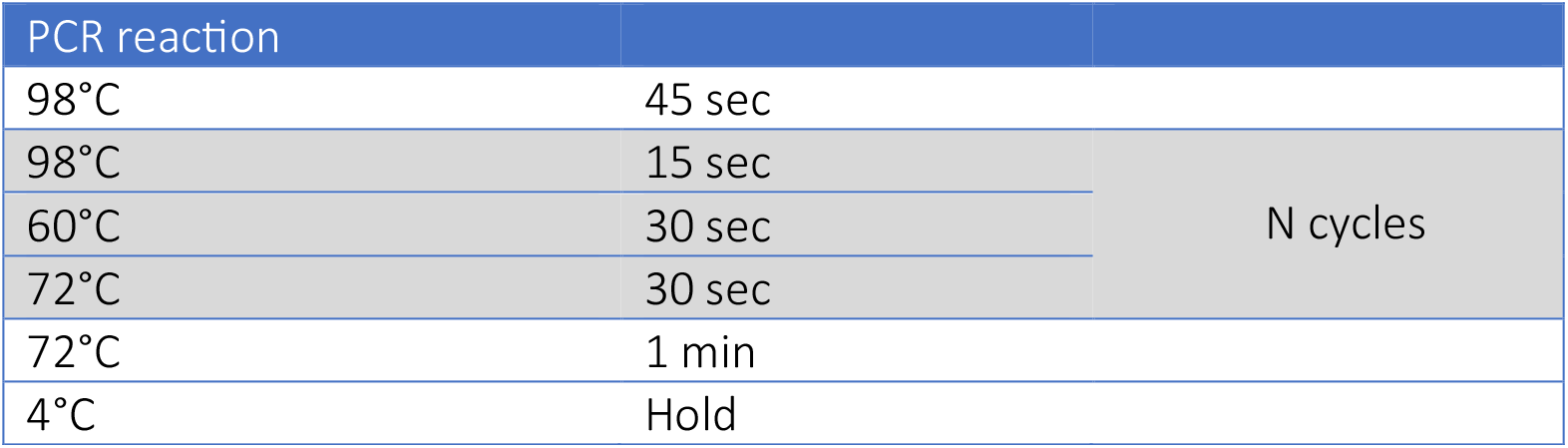 *The number of cycles depends on the panel size*. 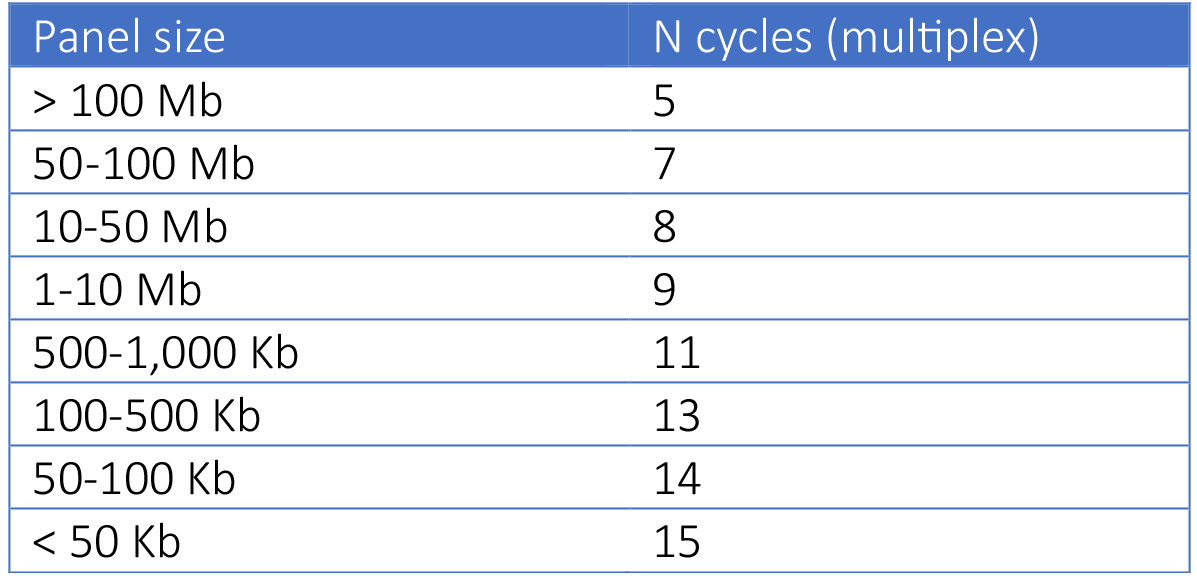
163. Pool all PCR reactions into a 1.5-mL microcentrifuge tube and perform DNA purification as described in Steps 44-50 using 1.8X NGS beads.
164. In the final step, elute into 100 µL of dd. H_2_O and transfer the cleared eluate to a fresh 1.5-mL microcentrifuge tube.
165. Run the sample on TapeStation or Fragment Analyzer to assess the final library quality and measure DNA concentration using Qubit HS Kit.

## SAFE STOPPING POINT

*Store the sample at -20°C for long-term storage*.

### VII Sequencing

Assess the final library on Fragment Analyzer and measure DNA concentration using Qubit BR Kit.

#### A. Pair-wise Module (iMCC/iTMCC/iMCCu)

Sequence libraries using a 300-cycle kit (e.g., Illumina, cat. no. 20100984) as 150 bp paired-end reads.

#### B. Multi-way Module (mwMCC)

Sequence libraries using a 600-cycle kit (e.g., Illumina, cat. no. 20100985) as 300 bp paired-end reads.

##### NOTES

▪ Digitonin permeability test Prepare 1% digitonin stock (5 mg digitonin in 500 µL of DMSO). Test a range of final digitonin concentrations: 0,001%, 0.0025%, 0.005%, 0.0075% and 0.01%.
  1. Trypsinize (if adherent) and centrifuge cells for 5 min at 300 x g and RT. Resuspend the cell pellet in media. Count cells and split into aliquots of 5×10^6^ cells each to test different digitonin concentrations.
  2. Add digitonin at desired final concentrations and mix by inversion.
  3. Incubate for 10 min at RT.
  4. Take 10 µL cells and mix with 10 µL Trypan Blue. Observe under microscope.
  5. Select the lowest concentration which results in complete cell permeabilization.

##### Sample

HCT-116 cells

Digitonin [final] concentration

Cells permeabilized

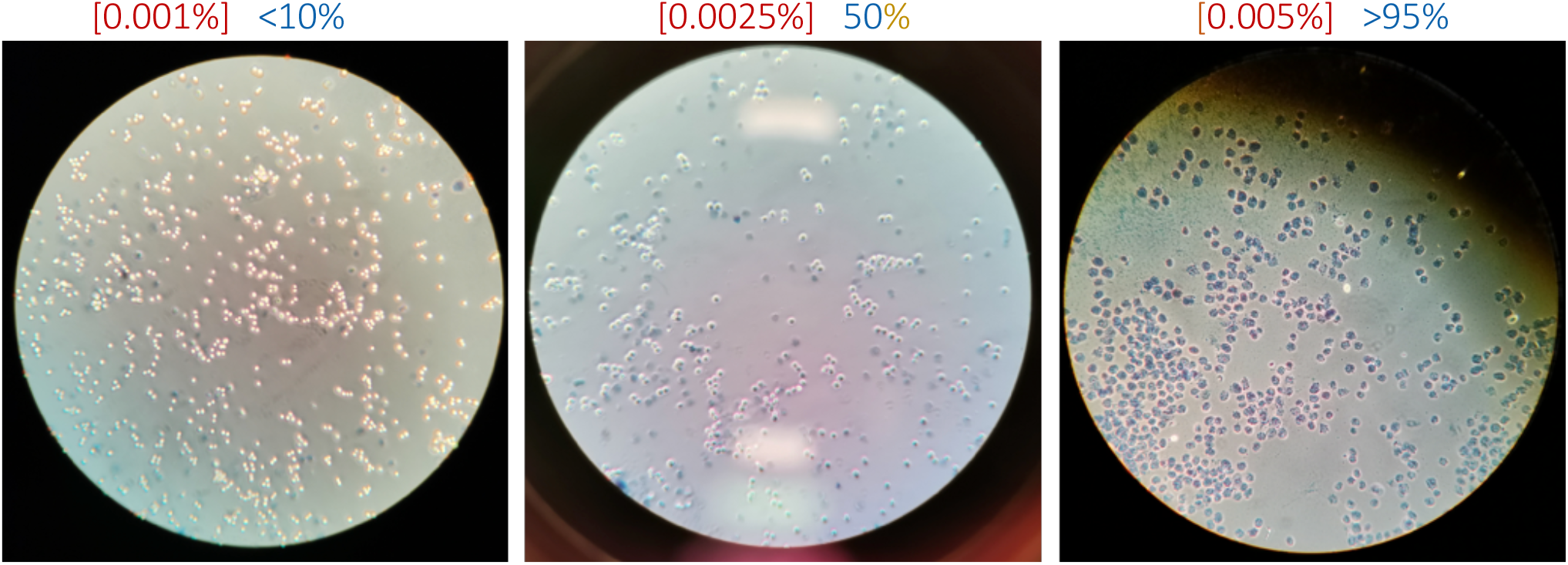

▪ Oligonucleotide design for capture enrichment Biotinylated capture oligonucleotides are designed using a python-based oligonucleotide tool (https://oligo.readthedocs.io/en/latest/). For iMCC and mwMCC, which target single viewpoints, 120 bp oligonucleotides are designed to center on the nucleosome-depleted region of the target locus. For tiled approaches, 70 bp (iTiled-MCC) or 120 bp (iMCCultra) oligonucleotides are designed by tiling across the entire region of interest, with consecutive oligonucleotides overlapping by 50%.
▪ Exemplary desired sample profiles Digestion and Ligation (all methods): 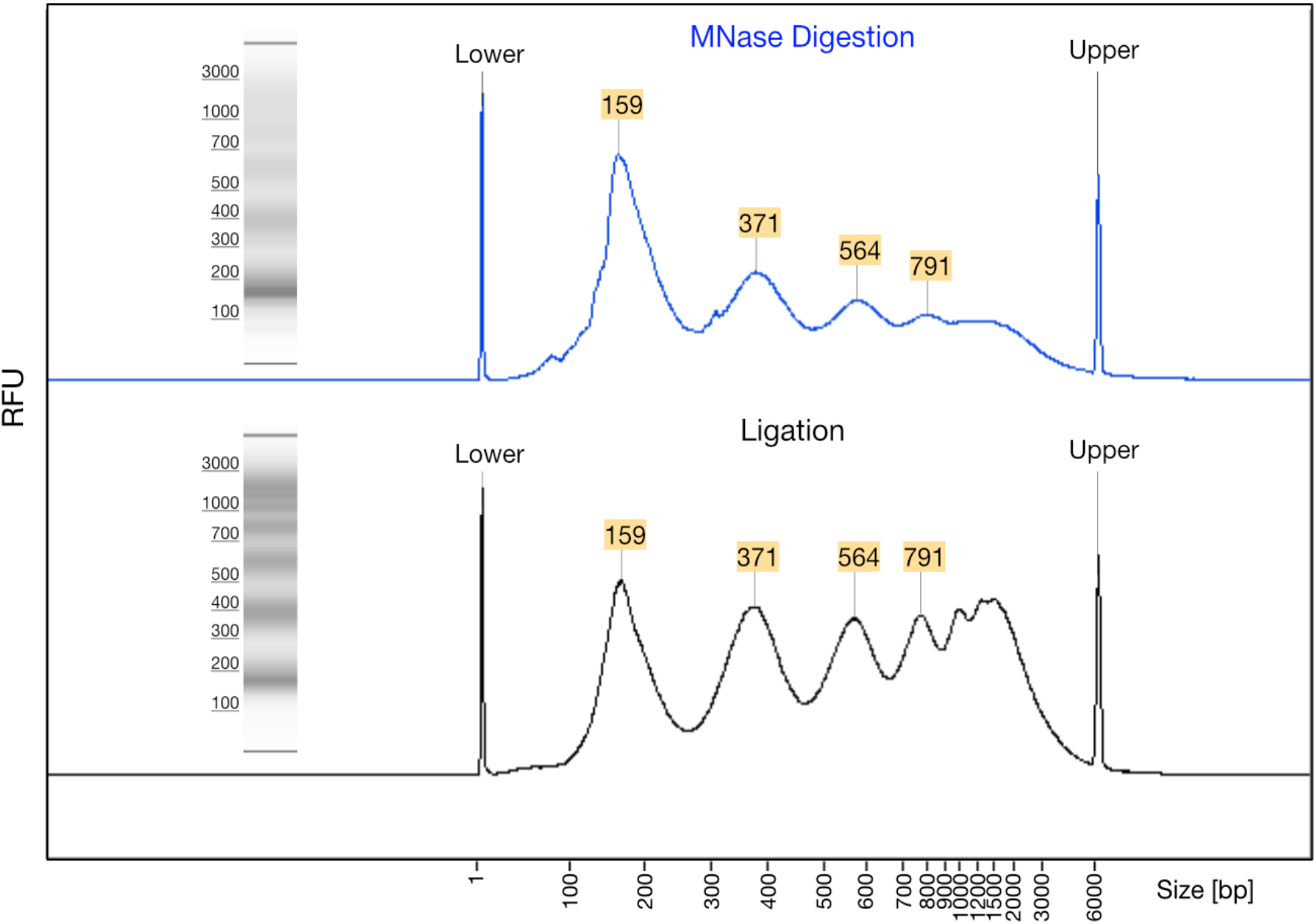 iMCC/iMCCu/iTMCC: 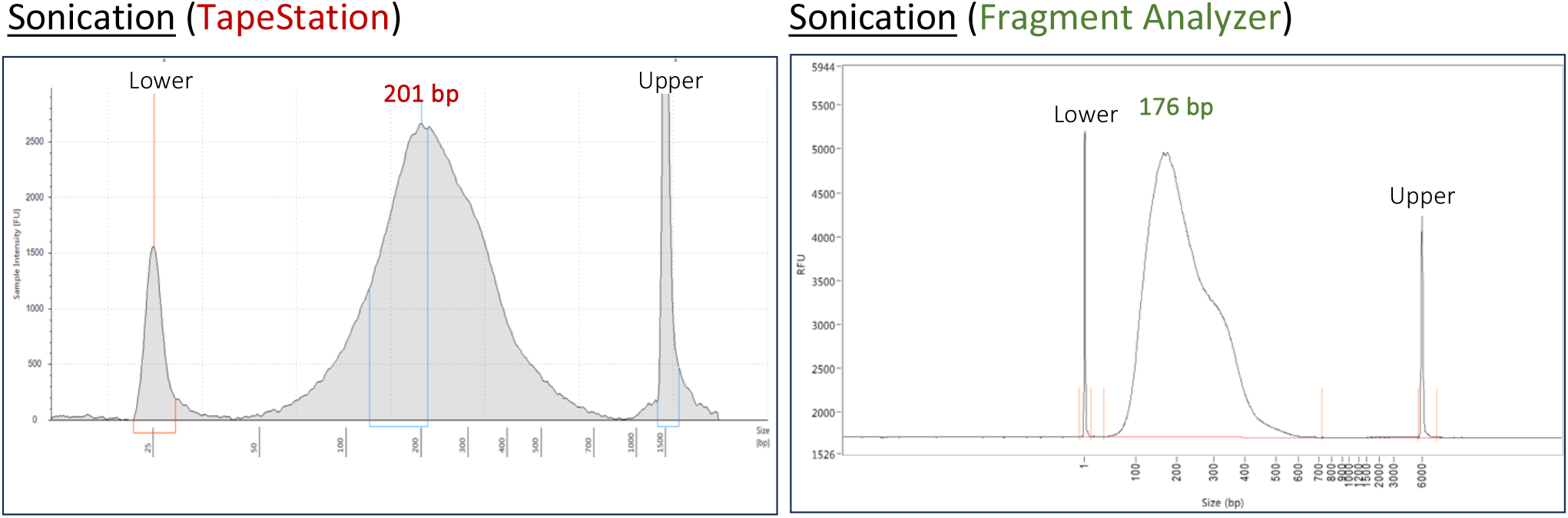 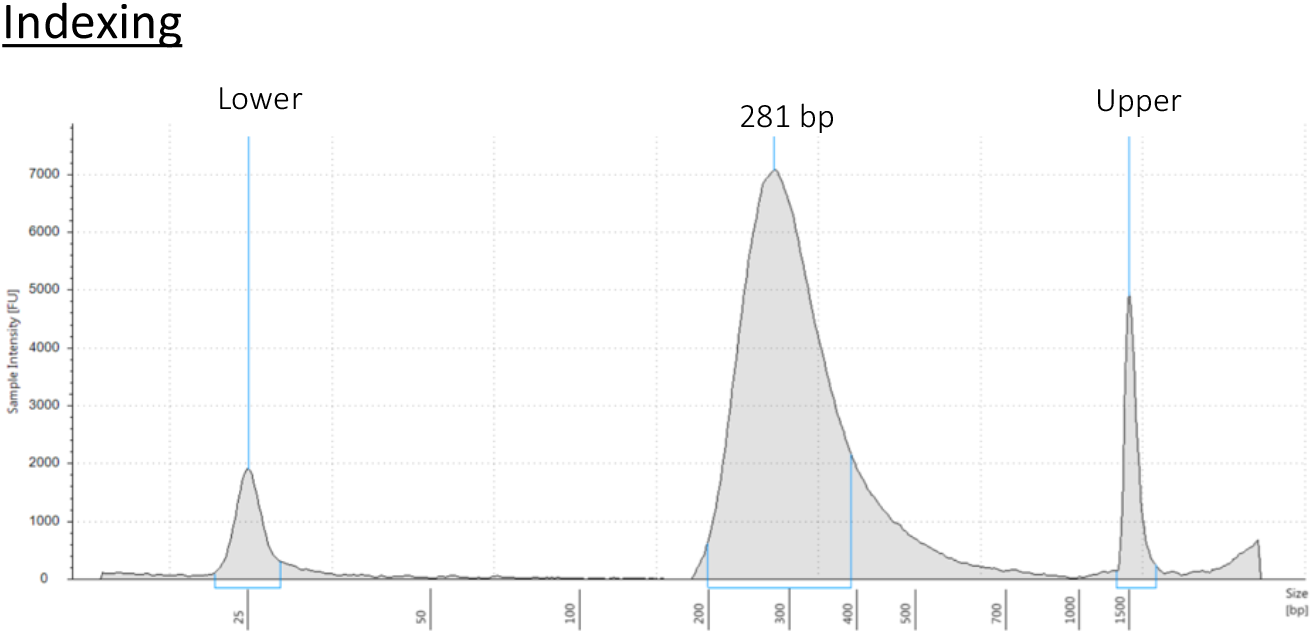 *The profiles after the Pre-Capture Biotin Clearing and Capture Enrichment should be similar to the Post-Indexing profiles*. mwMCC: 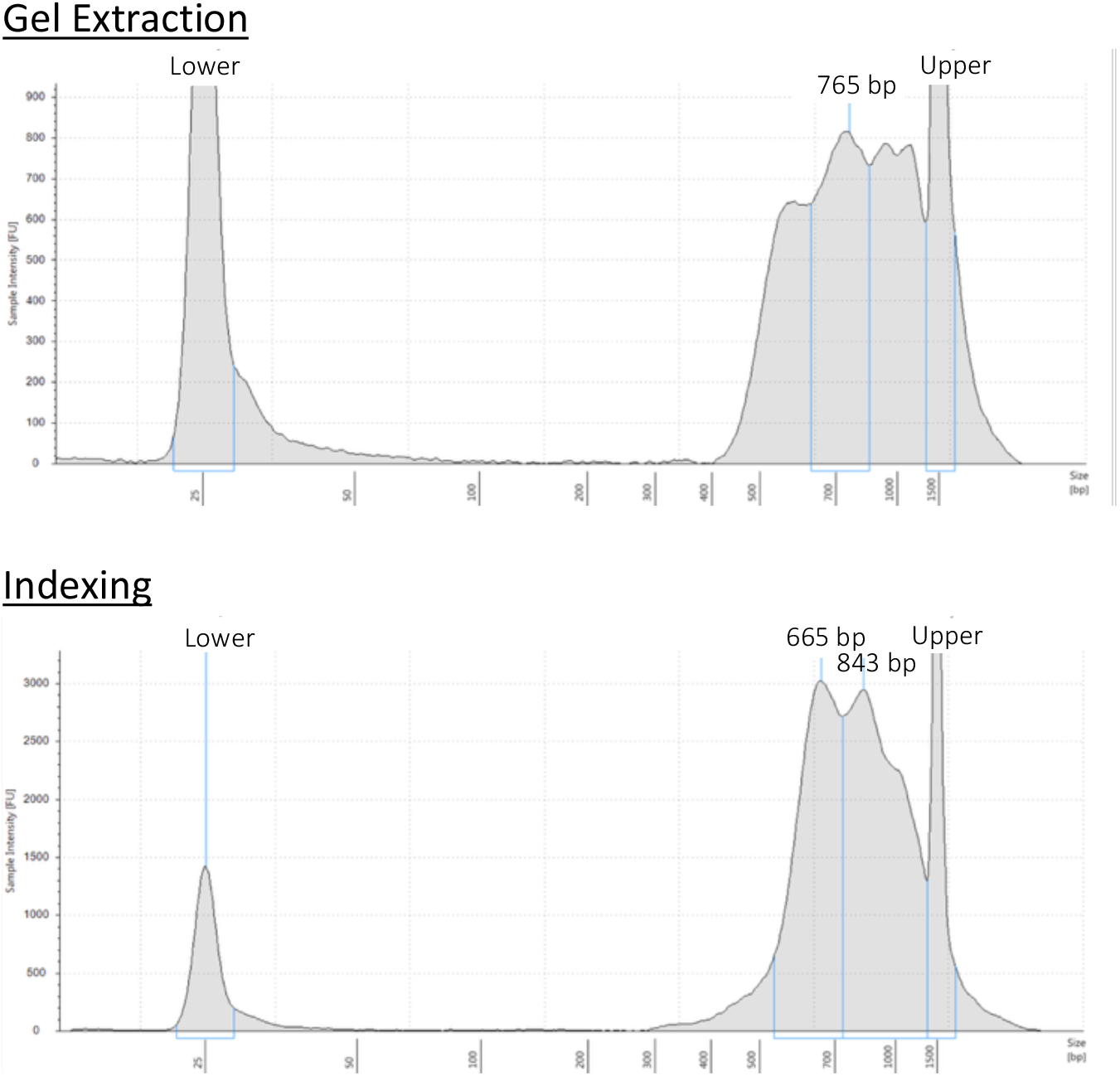 *The profiles after the Pre-Capture Biotin Clearing and Capture Enrichment should be similar to the Post-Indexing profiles*.

